# MAPLE: a Modular Automated Platform for Large-scale Experiments, a low-cost robot for integrated animal-handling and phenotyping

**DOI:** 10.1101/239459

**Authors:** Tom Alisch, James D. Crall, Dave Zucker, Ben de Bivort

## Abstract

Genetic model system animals have significant scientific value in part because of large-scale experiments like screens, but performing such experiments over long time periods by hand is arduous and risks errors. Thus the field is poised to benefit from automation, just as molecular biology did from liquid-handling robots. We developed a Modular Automated Platform for Large-scale Experiments (MAPLE), a *Drosophila-handling* robot capable of conducting lab tasks and experiments. We demonstrate MAPLE’s ability to accelerate the collection of virgin female flies (a pervasive experimental chore in fly genetics) and assist high-throughput phenotyping assays. Using MAPLE to autonomously run a novel social interaction experiment, we found that 1) pairs of flies exhibit persistent idiosyncrasies in affiliative behavior, 2) these dyad-specific interactions require olfactory and visual cues, and 3) social interaction network structure is topologically stable over time. These diverse examples demonstrate MAPLE’s versatility as a general platform for conducting fly science automatically.

## Introduction

Genetic model organisms are used to advance our biological understanding in numerous areas including disease and its treatment, basic cell biology, neuroscience and behavior. Species like *Mus musculus, Caenorhabditis elegans* and *Drosophila melanogaster* have advantages including rapid reproduction, ease of rearing, and deep genetic toolkits composed of strains with varying genotypes and transgenic alterations that permit rapid, mechanistic inquiries. To take advantage of these toolkits, screen experiments quantify the phenotypes of hundreds (Vitaterna et al., 1994), thousands (Kain et al., 2012), tens of thousands (Ayroles et al., 2015; Buchanan et al., 2015; Churgin et al., 2017) or even hundreds of thousands of individual animals (Robie et al., 2017). With the ongoing improvement and widespread adoption of high-performance machine vision phenotyping (Branson et al., 2009; Dankert et al., 2009; Kabra et al., 2012; Kimura et al., 2014), the time needed to manually handle experimental animals remains the bottleneck limiting data collection. Moreover, the repetitive labor of screening can be a deterrent to experimentalists. Systems that automate animal-handling thus have the potential to speed scientific findings, and can offer parity in high-throughput breadth to the high-throughput depth of modern molecular and neurobiological experiments — e.g., RNAseq (Macosko et al., 2015) or whole-brain imaging (Ahrens et al., 2013).

The advent of liquid-handling robots has radically changed the face of molecular biology, enabling techniques and experiments that had long been imagined, but were too complex, lengthy, or tedious to have been previously realized. We looked for an analogous system for *Drosophila*, but came up short. A number of animal-handling platforms can dispense or convey small organisms, such as the commercially successful BioSorter. However, as these systems require a liquid medium and are limited to smaller animals, they cannot manipulate adult fruit flies. There are very large scale systems that work with adult flies on a vial-by-vial basis, in the form of “fly flipping” robots. However, these take up a whole room, generally in a core facility, and cost hundreds of thousands of dollars to purchase and maintain, and are therefore inaccessible to most labs. And while there are many examples of high-throughput phenotypic assays in *Drosophila* (Branson et al., 2009; Kabra et al., 2012; Kain et al., 2012; Buchanan et al., 2015; Geissmann et al., 2017), the systems that aren’t single-purpose still require human intervention to load and unload individual flies. Some researchers have used flies' natural tendency to climb up (negative gravitaxis) to isolate individuals for behavioral analysis (von Reyn et al., 2014), imaging (Medici et al., 2017), or microsurgery (Savall et al., 2015). But dependence on this particular behavior fundamentally caps throughput, artificially selects for a subset of a population, and limits eligible genotypes. An alternative approach, actively conveying flies with airflow (Macmillan et al., 2014) permits moving animals on demand, and opens the door for increased throughput.

Here, we present an automated platform that is high-throughput and flexible enough to assist in conducting diverse experimental protocols in *Drosophila*. Due to its **<u>modular</u>** design, the system can **<u>automate</u>** diverse phenotyping assays in flies (e.g., loading of individual fruit flies for circadian rhythm (Pfeiffenberger et al., 2010; Geissmann et al., 2017) experiments), as well as aid with lab chores (e.g., collecting virgin female flies for genetic crosses or passaging individual flies in controlled culture conditions for longevity assays). In addition to facilitating fly science, this flexible system has diverse potential applications for studying non-model organisms (e.g., automated, longitudinal behavioral monitoring within bumble bee colonies (Crall et al. 2015) in multiple colonies in parallel). The physical **<u>platform</u>** of this instrument integrates animal husbandry and phenotyping, permitting end-to-end experimental protocols. Its low cost and scalability permit **<u>large-scale experiments</u>** that take advantage of the many benefits insect model organisms offer, such as huge fly genetic libraries containing thousands of lines (Jenett et al., 2012; Thibault et al., 2004). We highlight MAPLE’s utility and versatility with two particularly time- and manual labor-intensive tasks: rapidly collecting virgin female flies, and large-scale longitudinal measurement of fly social networks and behavior.

## Results

### MAPLE physical implementation

With those high-level goals in mind, we designed the MAPLE system with the following design constraints: 1) it features a large, flat experimental workspace with room for multiple flexibly-configurable experimental modules, 2) this workspace is physically open for user convenience, and transparent on the top and bottom for in situ optical phenotyping, 3) multiple end-effectors can move throughout the workspace to handle flies and manipulate experimental modules, 4) it features failsafe mechanisms so that users can leave it unattended without worrying that it would damage itself or experimental modules, and 5) it is relatively inexpensive and scalable.

MAPLE (Fig 1A, S1) was built using extruded aluminum rails to support x-, y-, and z-carriages mounted on linear rails in a Cartesian configuration. We employed the CoreXY system (Moyer, 2012), which reduces the mass of the moving part of the X/Y gantry by fixing the stepper motors on the frame. Although, for the speeds at which we run MAPLE, which are roughly 80% as fast as human hands conducting experiments (Movie S1), mounting the y-axis stepper motor on the x-axis carriage would likely not reduce performance.

**Figure 1.**
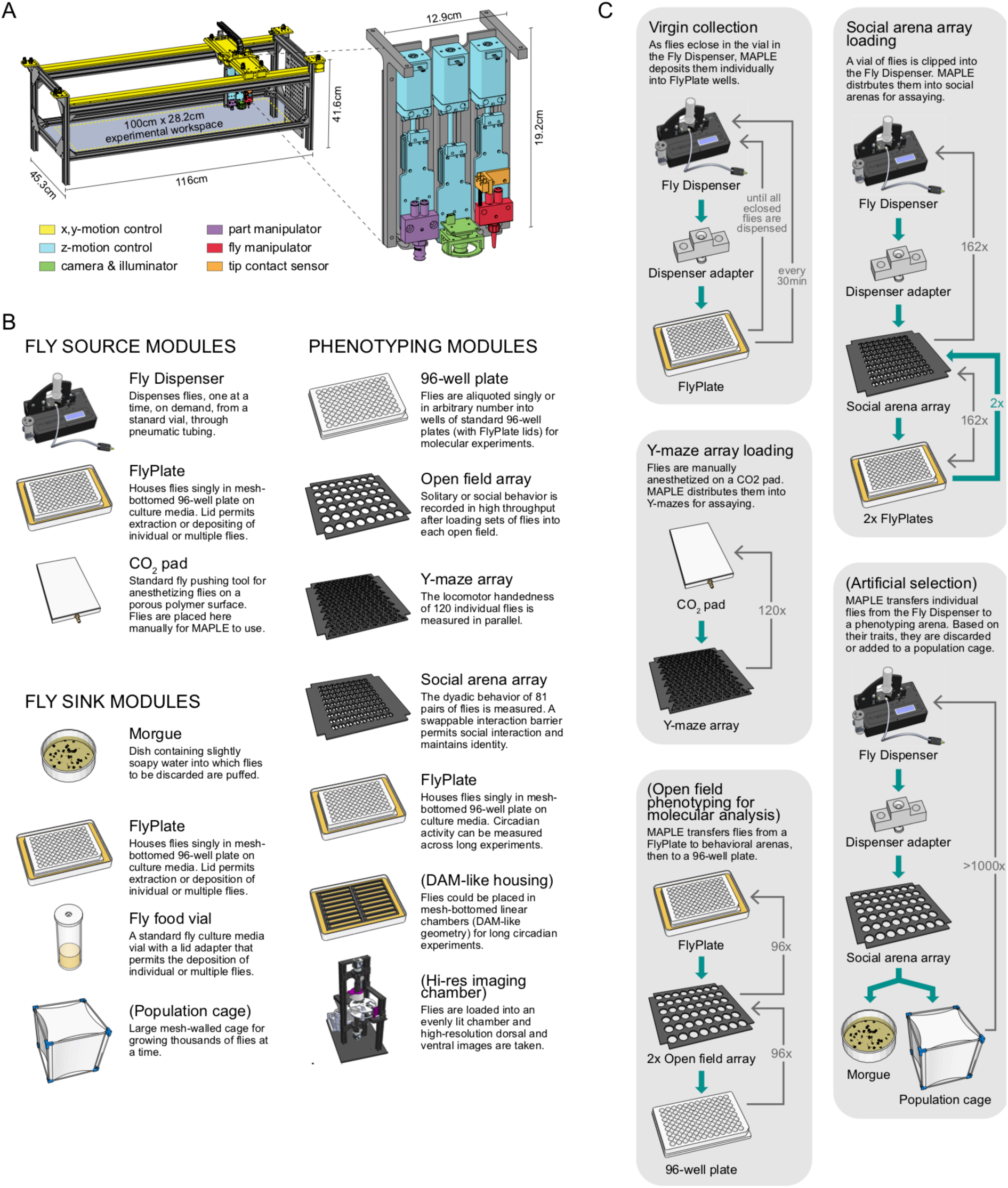
MAPLE hardware, modules and experimental procedures. —A) Schematic of MAPLE with workspace dimensions indicated and an expanded schematic of the end effectors. Colors indicate different robotic systems. B) Illustrations and brief descriptions of experimental MAPLE modules. Fly source modules can provide flies into the MAPLE system. Fly sink modules can receive flies. Phenotyping modules are used to collect experimental measurements. See Supplementary methods for more extensive descriptions. Modules with names in (parentheses) are under development. C) Simplified flowcharts illustrating a selection of tasks MAPLE is capable of performing using different combination of modules. Green arrows indicate the flow of animals through the task, thin grey arrows the motion of the MAPLE end effectors. Tasks with names in (parentheses) are hypothetical and illustrate the scope of possibilities. This figure is also a target for an augmented reality view of MAPLE. Print the figure in portrait orientation on letter paper, place the printout on a horizontal surface like a bench or desk, and view it in scan mode in the “Augment” mobile app to interact with a to-scale rendering of MAPLE.

In terms of our design constraints: 1) the experimental workspace measures 100 × 28.2 × 7.5 cm on the (x-, y-, and z-axes respectively). Its floor is clear acrylic with cable pass-throughs. Locating brackets (laser cut out of 6mm acrylic) were affixed to an interchangeable acrylic surface with the same footprint as the floor (a “workspace plate”), allowing experimental modules to be precisely and repeatably positioned within and removed from the workspace (Fig S2). Interchanging workspace plates allows rapid reconfiguration of the workspace for different experimental procedures. 2) The sides, top, and bottom of MAPLE are open or made of clear acrylic, permitting the optical phenotyping of flies in experimental modules at all time points other than when the end-effector carriages are above the modules. 3) The end-effector assembly comprises 3 independent z-axes, each featuring a single tool (Fig 1A; Movie S2): a small-part manipulator for picking up (using vacuum) experimental module components like plastic lids; a USB digital camera with LED illumination for acquiring high-resolution images for machine-vision; and a fly manipulator for handling awake or anesthetized flies using vacuum (Fig S3). 4) All motion axes have physical limit switches and/or software limits preventing overtravel. The fly manipulator end effector, which is rigid and must align precisely with experimental modules at different heights, is equipped with a collision-detection switch to halt z-motion before the robot is damaged. This sensor can also be used to detect the height of rigid module components. It is safe to leave MAPLE unattended (Movie S3). 5) MAPLE components cost approximately $3500 (Supplementary Document 1). Its bill of materials, assembly instructions (Supplementary Document 2), and code libraries (see Methods for links) have been made public under open source licenses. MAPLE measures 43.5 cm in the y-dimension permitting mounting in standardized 19” rack systems, so multiple robots can be arranged compactly.

### MAPLE Experimental Modules and Software

We have fabricated and used a number of experimental modules to test the notion that with the right diversity of reconfigurable modules, MAPLE can implement a great diversity of protocols. These modules fall into three categories (Fig 1B): fly source, fly sink, and phenotyping modules. Fly source modules are repositories from which flies can be removed in a controlled fashion and transferred into downstream modules. These include the Fly Dispenser (FlySorter LLC), a small device that outputs single flies, on demand, from a standard plastic vial pre-loaded with many flies. It can be triggered to dispense a fly via a serial command sent over USB. A standard CO_2_ pad with a porous polyethylene surface, used to anesthetize flies manually at the start of a MAPLE session, serves as a source of flies. Using machine vision, MAPLE is capable of recognizing flies’ positions on the pad. FlyPlates (FlySorter LLC) are modified 96-well plates in which the floor has been replaced with a metal mesh, allowing flies stored in the wells to feed on fly media below the plate. The lid features a nylon mesh with triangular flaps cut above each well, which allow an aspirator tip or the fly manipulator end-effector to enter the well, retrieve or deposit a fly, and leave the well without permitting the fly to escape. Because flies can be deposited in this module, it also falls in the category of fly sink modules.

Fly sink modules are destinations into which flies that have been handled by MAPLE can be deposited. In the case of the FlyPlate, deposited flies can be later removed. Other fly sink modules are one-way, including a morgue, a dish of slightly soapy water or 70% ethanol covered with a nylon mesh lid in the style of the FlyPlate. Standard fly culture media vials with nylon mesh loading adapters can be used to collect many flies after MAPLE handling for long-term storage.

The last category of modules is phenotyping modules, which produce experimental data. We have created a number of phenotyping modules with a focus on collecting behavioral data, including arrays of circular open field arenas, arrays of Y-shaped mazes for measuring locomotor handedness (Buchanan et al., 2015), and social arena arrays, in which pairwise social interactions can be monitored. All of these phenotyping modules can receive flies in one of two different ways, depending on whether the flies are anesthetized or not. In a traditional experimental style (Ayroles et al., 2015; Buchanan et al., 2015), MAPLE can pick up anesthetized flies from the CO_2_ pad module with the fly manipulator, pick up the plastic lid covering a behavioral arena with the small part manipulator, drop the fly in the arena, and replace the lid. In a MAPLE-optimized experimental style, awake flies are retrieved from the Fly Dispenser or FlyPlate and then transferred directly into the behavioral arena by sliding a multi-position loading port into place above the arena, dropping the fly, and then sliding the port so it is inaccessible (Fig S4 and examples below).

Phenotyping modules are not limited to collecting behavioral data. For example, flies can be loaded into standard 96-well plates outfitted with nylon mesh lids. From there, they are ready for molecular protocols, including use by liquid-handling robots. Flies in FlyPlates, which have access to food, can be used for circadian assays (Tataroglu et al., 2014), longevity (Stearns et al., 2000), or pharmacological experiments (Gasque et al., 2013). Modules such as the Fly Dispenser can be situated outside MAPLE’s frame, receiving and delivering flies via air-flow tubing. 3d-printed adapter blocks connect these tubes into the MAPLE workspace and are situated with locating brackets. Thus, all modules have some component that is placed within the MAPLE workspace, even if the bulk of the device is outside.

The modularity of MAPLE’s hardware is reflected in its software as well. Each experimental procedure (examples in Fig S5) is associated with a Python experimental script file. These files call on functions (Fig S6) that 1) implement common multi-step robot actions (like retrieving a fly from a behavioral arena), 2) implement low-level elemental robot actions, 3) are specifically associated with the modules used in a particular experiment, and 4) mediate the remote control of MAPLE over the internet. This software architecture permits the rapid scripting of new experimental protocols at a relatively high level.

### MAPLE is flexibly reconfigurable

To demonstrate the robot’s efficacy at running experiments and accomplishing lab tasks, we implemented a number of experimental procedures (Fig 1C). These include: 1) collecting virgin flies for genetics crosses by dispensing flies as they eclose and then distributing them into individual wells of FlyPlates where their isolation preserves their virgin status indefinitely; 2) loading flies into Y-shaped arenas to measure their locomotor biases, the time-consuming step of a routine assay in our lab; 3) loading flies from the Fly Dispenser into the FlyPlate wells for long-term culturing and circadian phenotyping; and 4) loading flies into social arenas to measure their pairwise interactions (Movies S2, 4).

While MAPLE was conceived for conducting experiments and lab tasks with fruit flies, its modularity, open-source design, and hierarchical software architecture facilitates uses for different species. As an example, we adapted the basic MAPLE design for high-throughput imaging of uniquely identified workers within colonies of the Common Eastern bumble bee *(Bombus impatiens)*. In the Bee Experimental Ethology Colony Hardware (BEECH) system (Fig 2), up to 12 colonies of ~50 bumble bees each are housed in acrylic colony boxes featuring a dark nest chamber (Movie S5) and a circadian-lit foraging chamber, where bees are supplied with nectar and pollen. This MAPLE-derived 2-dimensional Cartesian robot moves a multi-camera end-effector from colony to colony, recording high-resolution video of the nest and foraging chambers (Movies S5, 6), which permits automated tracking of individual bees using machine vision (Crall et al., 2015) for up to several weeks. Beyond replacing MAPLE’s z-axis assembly with multiple cameras and extending all three dimensions of MAPLE’s frame to accommodate the 12 colony box modules, BEECH employs identical frame architecture and motion-control, demonstrating the versatility of the MAPLE design.

**Figure 2.**
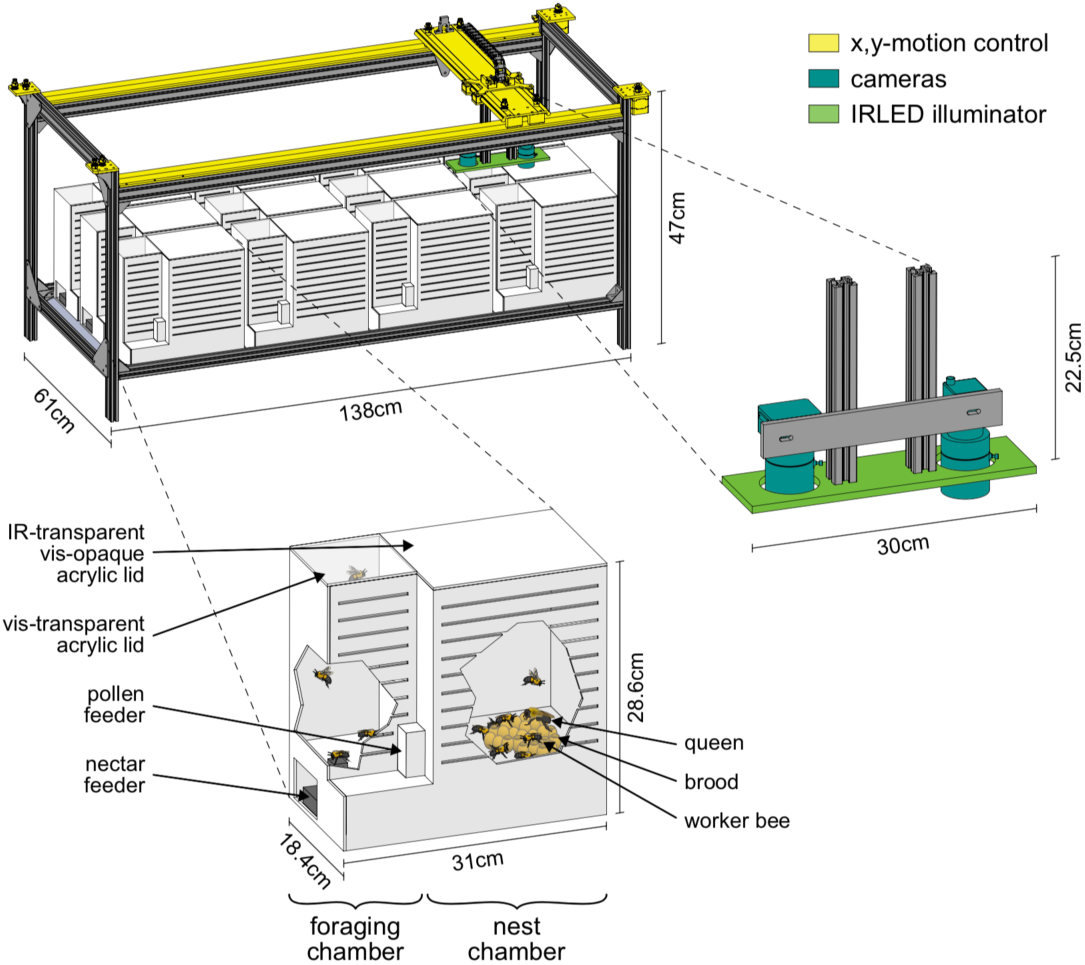
Bee Experimental Ethology Colony Hardware (BEECH): a MAPLE-derived robotic platform for imaging bumble bee behavior. —A MAPLE-derived Cartesian robot (top left) was used in an automated system for recording bumble bee behavior in multiple colonies over long periods of time. The end effectors of this robot were IR-sensitive cameras (right) for recording digital video of bees behaving. Bees were housed in acrylic colony boxes (bottom) with a dark nest compartment and circadian-lit foraging compartment where they, respectively, reared developing young and collected nectar and pollen from feeders.

### MAPLE-handled flies exhibit normal behavior

To confirm that handling by MAPLE did not damage flies or introduce discrepancies compared to experiments conducted manually, we compared the locomotor performance of flies handled by MAPLE and unhandled flies. First, we manually aspirated 96 anesthetized flies into a FlyPlate. After 2 hours of rest, half the flies (48) were subjected to repeated manual removal and replacement back into their well using MAPLE, while the remaining flies (48) were left unhandled (Fig 3A,B). After this handling procedure, the entire plate was imaged in a backlit motion tracking rig (Buchanan et al., 2015). Handling and imaging was then repeated an additional 4 times to test for cumulative effects. Flies handled by MAPLE were statistically indistinguishable from unhandled flies both in the fraction that were active across imaging sessions (Fig 3C; *b* = −0.02, *t*(477) = −0.098, *p* = 0.92 by multinomial logistic regression) and in their mean walking speed across imaging sessions (Fig 3D; F(4, 376) = 0.51, *p* = 0.73 by mixed-effects ANOVA). Second, we measured the locomotor bias of individual flies in Y-shaped mazes (Ayroles et al., 2015; Buchanan et al., 2015) configured with multi-position loading ports. Awake flies were loaded into these mazes from FlyPlates by either manual aspiration or MAPLE-handling (Fig 3E,F). The across-individual distribution of walking speeds was statistically indistinguishable between the manual and MAPLE-handled group (Fig 3G; *p* = 0.61 by KS-test). Likewise, the across-individual distribution of turning bias (the tendency of individuals to turn left or right at the choice point in the center of the Y-maze) was indistinguishable between handling treatments (Fig 3H; *p* = 0.74 by KS-test). The same was true for comparisons of manual and MAPLE-handled behavior in semi-circular arenas (Fig S7).

**Figure 3.**
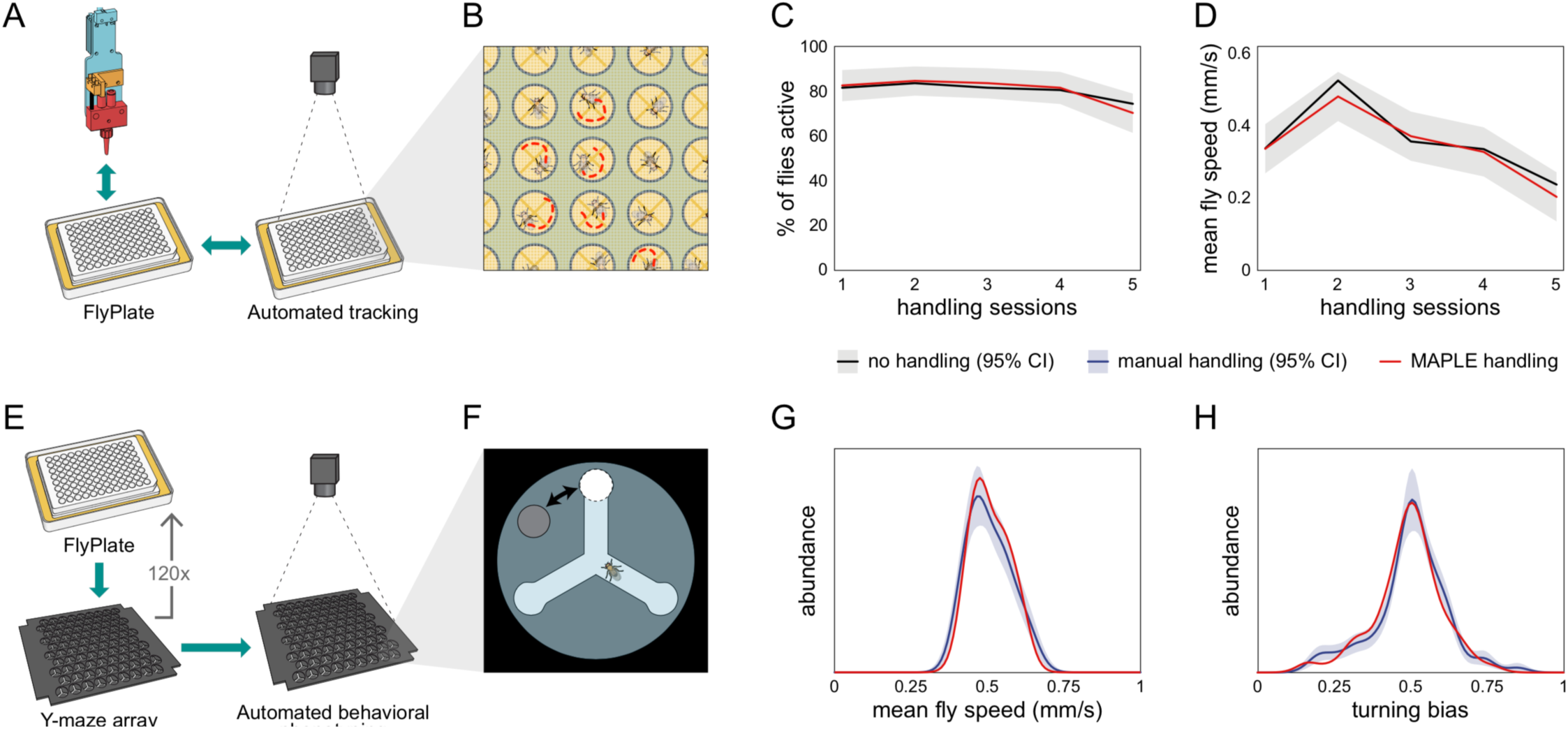
Behavior of flies manipulated by MAPLE is comparable with manual controls. —A) Diagram of repeated MAPLE handling procedure. Flies were removed from and replaced back into wells of a FlyPlate. In between these handling events their activity was assessed by automated tracking. B) Illustration of automatic tracking of flies in a FlyPlate through the nylon mesh covering the wells. C) Percentage of flies exhibiting supra-threshold activity (mean speed > 0.1 mm/s) across handling sessions. Red line is MAPLE-handled flies, black line is matched unhandled flies. Gray area denotes 95% CI around unhandled flies. D) Fly mean speed across handling sessions. E) Illustration of MAPLE-assisted handedness phenotyping procedure. F) Y-maze arena adapted for MAPLE use with a multi-position loading port allowing the deposition of awake flies into the behavioral arena. G) Distribution of fly mean speeds in MAPLE and manually loaded experiments. Distributions shown are kernel density estimates (KDEs). Gray area is 95% CI around manually loaded flies as estimated by bootstrap resampling of KDEs. H) Distributions of fly turning bias (# right turns / # total turns) in MAPLE and manually loaded experiments.

### MAPLE-assisted virgin picking is more efficient than manual methods

Next, we tasked MAPLE with performing a tedious task that consumes great amounts of time in essentially all *Drosophila* labs: collecting virgin females for genetic crosses. Female *D melanogaster* will not mate with males for approximately 6 hours after they eclose from the pupal case. In the first portion of this interval, they have morphological characteristics (puffy abdomens, translucent cuticle, and visible meconium in the gut) that correlate with their young age and are reliable, but conservative, indicators of virginity. In traditional manual virgin-picking, only females with these morphological correlates are collected. This means that many virgin females lacking morphological correlates in the latter portions of the 6-hour no-mating window are discarded, unless practitioners collect virgins from a stock vial or bottle at regular intervals at least 3 or 4 times a day.

MAPLE has the potential to recover 100% of females as virgins by isolating them quickly after they eclose, and storing them individually to preclude mating (Movie S7). To implement this procedure, we devised a simple custom fly media vial in which the lower portion containing fly food is detachable. Parental generation flies lay eggs in this food, or food from another vial containing larval flies can be transferred into the custom vial. When flies in this experimental generation climb onto the walls of the vial and pupate, the food-containing portion is manually detached, and replaced with an empty vial-bottom (Movie S7). This pupae-containing, foodless vial is then placed in the Fly Dispenser. Every 30 minutes, all newly eclosed are dispensed into MAPLE, which distributes them individually into the wells of a FlyPlate (Fig 4A).

**Figure 4.**
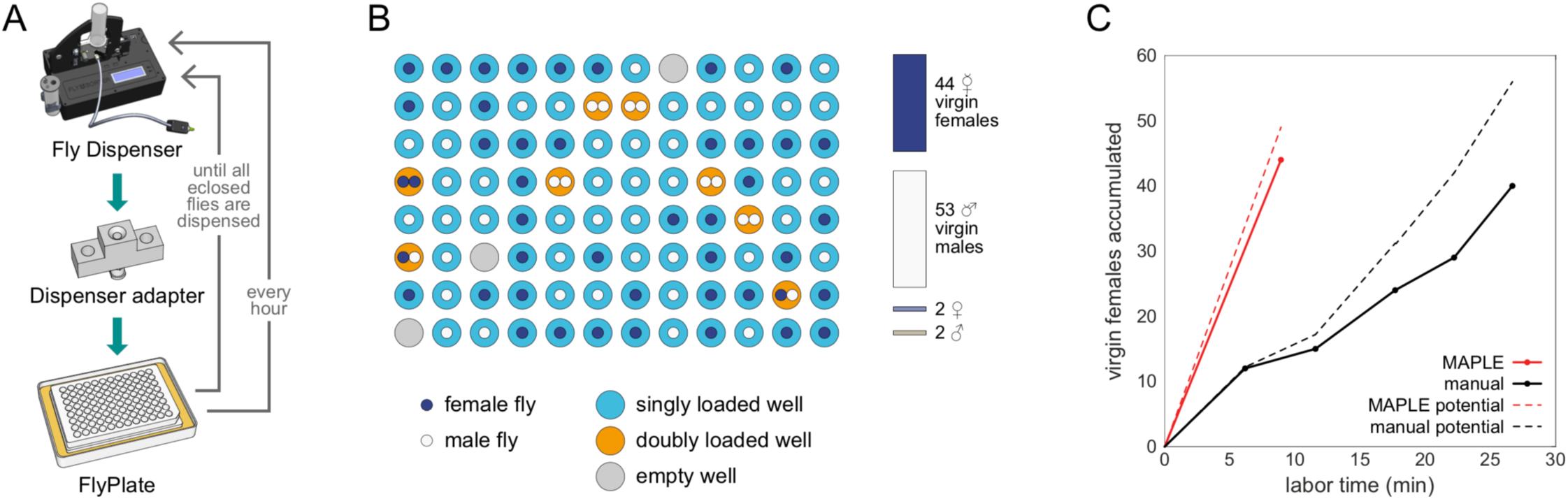
MAPLE performance at collecting virgin female flies for genetic crosses. —A) Diagram of modules and procedure employed in MAPLE virgin-collecting. B) Representative illustration of the well content of a full FlyPlate after 3 days of virgin collecting. Outer circle color indicates the number of flies loaded. Inner circles indicate the sex of deposited flies. C) Comparison of cumulative number of virgin females collected versus manual labor time required in MAPLE and conventional manual virgining procedures. Dotted lines indicate the maximum potential virgins that might have been collected in each approach. Thus the difference between the dotted and solid line indicates females that were “lost” and not retained as virgins.

This process was not without error: at rates of ~3% a well was left empty, or ~8% a well was loaded with two flies. A representative outcome is shown in Fig 4B. To recover only the virgin females, fully loaded FlyPlates were removed from their media base, brought to traditional CO_2_ fly-pushing pads at dissecting microscopes, the flies were anesthetized through the wire mesh floor, and then the plate and pad assembly was inverted, leaving the flies on the CO_2_ pad in their respective positions from the wells. The sex of each fly was determined by eye, and females that had been stored either alone, or with no males were collected. In a head-to-head comparison of typical manual virgin-collecting (picking at the beginning and end of the work day from 5 vials) versus MAPLE virgin-collecting, virgin females were procured at a rate of 1.5/min (including the time needed to bring vials from the incubator, anesthetize them, etc.) and 4.9/min using MAPLE (including the time to set up the pupa vial, load the dispenser, manually sex and sort the flies, etc.) (Fig 4C). In the future, we anticipate MAPLE will be able to sex flies without human intervention using the high-resolution imaging module, further decreasing manual work required.

### MAPLE reveals persistent fly social interaction networks

Lastly, we set out to see if MAPLE could conduct experiments that would be daunting by traditional manual methods. Specifically, we set out to measure social interaction networks (SINs; Schneider et al., 2012; Pasquaretta et al., 2016) between pairs of flies, and then determine if measures of pair-wise affiliation are preserved on the timescale of days (Fig 5). It is known from group behavioral experiments that individuals spend more time interacting with specific other individuals (Simon et al., 2012). Because it is challenging to perfectly maintain individual identity through such experiments, as well as retrieval, storage, and finally retesting, it is unknown if dyad-specific affiliative measures are stable over time. To resolve this uncertainty, we devised a new high throughput assay to measure affiliative behavior between pairs of flies (Fig 5B). This consisted of a 9x9 array of adjacent, approximately semi-circular arenas separated by an interchangeable barrier. Individual flies can be loaded into each arena-half by MAPLE using a multi-position loading port. We made 4 versions of the interchangeable barrier: an open-clear barrier was made of clear acrylic with grooves to connect the half-arenas, permitting flies to see and easily smell each other; a solid-clear barrier permits flies to see each other but impedes airflow; an open-black barrier permits airflow but blocks visual cues; and a solid-black barrier blocks both airflow and visual cues.

**Figure 5.**
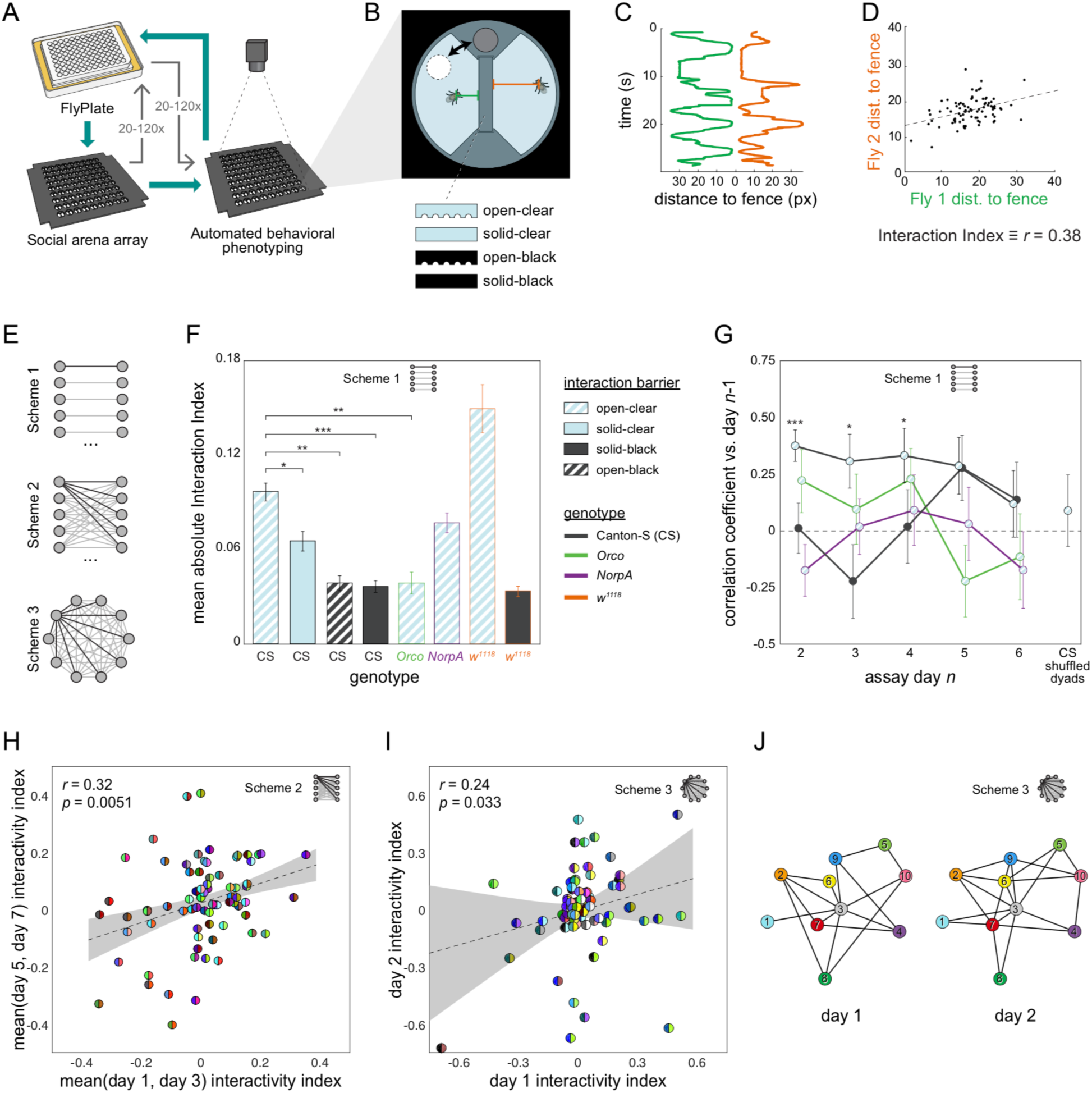
Persistent fly social networks measured using MAPLE. —A) Diagram of MAPLE-assisted social interaction behavior phenotyping procedure. B) Schematic of one of 81 social arenas including interchangeable interaction barrier types. A multi-position loading port allows flies to be loaded or removed from either compartment independently. C) Individual flies’ distances to the interaction barrier over time in a representative session. D) An example dyad interactivity index, calculated as the correlation between individual flies’ interaction barrier distances. E) Diagram of pairing schemes employed to form dyads. F) Mean absolute interactivity indices across interaction barrier types and genotypes. Dyads were formed according to scheme 1. Patterns denote barrier type, outline colors denote genotype. Error bars correspond to +/- 1 SEM. Mean absolute interactivity indices differ significantly across conditions, F(7, 853) = 13.42, *p* < 0.001. Asterisks indicate pair-wise comparisons that are significant by *t*-test. *: significance at a = 0.05; **: significance at a = 0.01; ***: significance at a = 0.001. G) Pearson correlation coefficient between dyad interactivity indices measured on successive days, over 6 days. Dyads were formed according to scheme 1. Point patterns denote barrier type, line colors denote genotype. Error bars correspond to +/- 1 SEM. Asterisks denote significant two-tailed z-tests. H) Scatter plot of interactivity indices of Canton-S (wild type) flies in arenas with open-clear barriers from measurements made across 2–4 day intervals. Dyads were formed according to scheme 2. Gray area is 95% CI of the linear regression line. Circles represent dyads; semi-circles denote individual flies forming a dyad; colors denote fly identity. Pearson correlation coefficient is statistically significant, *r*(99) = 0.32, *p* = 0.005. N: 99 dyads, 20 flies. I) As in (H) for two groups of 10 virgin female Canton-S forming 45 dyads each according to scheme 3 and tested on successive days, *r*(78) = 0.24, *p* = 0.034. J) Visualization of a Social Interaction Network (SIN) of 10 Canton-S (45 dyads) in arenas with open-clear barriers. Connections denote dyads exhibiting absolute interactivity index values greater than the average of the absolute values of the 1st and 4th quartiles (threshold: 0.031). Threshold was identical for day 2. Colors and numbers indicate fly identity on both days.

As a measure of social affiliation, we determined an “interactivity index,” defined as the Pearson correlation coefficient of the distance of each fly to the barrier over time (Fig 5C,D). Either significantly positive or negative values of this index indicate social interaction. We found that male-female and female-female, although not male-male, dyads produced the same distribution of interactivity indices (Fig S8A). Most social experiments were performed with virgin female-female dyads, though for some control groups we also included male-female dyads.

We programmed MAPLE to load pairs of flies from the FlyPlate into social arenas according to three different dyad schemes (Fig 5E). These varied from high-throughput (testing 162 flies at a time), with each fly tested in just a single dyad, to low-throughput (10 flies total) but saturated with respect to all possible dyadic pairings. Using the high-throughput scheme, we first measured the interactivity indices of wild type (Canton-S) flies separated by open-clear barriers. The observed distribution was significantly different compared to shuffled controls that scramble dyadic pairings (Fig S8B; *p* > 0.0001 by KS-test), indicating that interactions are dyad-specific.

Dyads separated by barriers that limit sensory cues exhibited significantly lower mean absolute interactivity indices (Fig 5F), indicating that both olfactory and especially visual cues drive differences in dyadic interactions. This is consistent with reports that visual cues can mediate social modulation of behavior (Kim et al., 2012). Consistently, anosmic *Orco* mutant flies (Vosshall et al., 2011) showed a significant 65% reduced mean absolute interactivity index, even with open-clear barriers *(p* = 0.00048 by *t*-test). Blind *NorpA* mutant flies (Kim et al., 1995) did not differ significantly from wild type flies in mean absolute interactivity index *(p* = 0.26 by *t*-test). Flies with mutations in the *white* gene *(w^1118^)* have reduced visual acuity (Markow & Scavarda, 1977) but showed a 50% increase in absolute interactivity index which was statistically significant *(p* = 0.013 by *t*-test), suggesting that social partner visual recognition does not require fine acuity (Justice et al., 2012). The increased inter-dyad variability seen in *w^1118^* animals may be consistent with increased inter-individual variability in phototactic preference exhibited by this genotype (Kain et al, 2012). In a control experiment, *w^1118^* flies separated by solid-black barriers had interactivity indices indistinguishable from Canton-S flies separated by solid-black barriers *(p* = 0.48 by *t*-test).

Lastly, we set out to determine if social behavior is stable over long periods of time by measuring dyadic interactivity indices across days. Using our high throughput dyad scheme, we measured interactivity indices from 91 dyads on each of 6 consecutive days. Interactivity indices across Canton-S dyads were statistically significantly correlated (0.31 < *r* < 0.38, 0.0001 < *p* < 0.04) between day 1 vs. day 2, day 2 vs. day 3 and day 3 vs. day 4 (Fig 5G), but not days 4 vs.5 or 5 vs. 6. Thus dyad-specific affiliative behavior appears to be stable over days-long timescales, though only for the first days of experiments. In control experiments, Canton-S flies with solid-black barriers and *NorpA* and *Orco* mutant flies exhibited no significant correlation in dyad interactivity indices across days. Significant correlation in interactivity index across dyads was observed in our two SIN dyad schemes as well (Figs 5I-J) suggesting that SIN persistence can be measured in a variety of experimental formats. Other metrics of social interaction yielded qualitatively similar results to the interactivity index (Fig S9; Supplementary methods).

## Discussion

Genetic model systems, like fruit flies, have tremendous potential to reveal the mechanistic underpinning of human diseases and basic biological phenomena like behavior. Much of this potential comes from examining behavioral and morphological phenotypes from a wide variety of genetic variants. As taking full advantage of the power of these systems requires extensive labor, automation has the potential to accelerate the production of data and thus scientific discovery. We developed MAPLE, a modular, automated platform for large-scale experiments, to greatly expand automated fly-handling experimental capabilities (Fig 1). By design, MAPLE is versatile, scalable, and relatively inexpensive, with all components of the core fly-handling robot costing roughly $3500. We displayed the versatility of this system by adapting the MAPLE platform to automatically monitor the behavior of multiple bumble bee colonies over long periods (Fig 2). We have made the design and code for MAPLE open access in the hope that it, or descendant approaches will be widely adopted by our field to fuel the accumulation of scientific findings.

After demonstrating that MAPLE did not discernibly alter the baseline behavior of animals compared to manual handling (Fig 3), we developed a procedure by which MAPLE could help conduct a repetitive lab chore that nearly all *Drosophila* biologists are familiar with — collecting virgin females for genetic crosses. Using a Fly Dispenser, MAPLE collected individual animals as they eclosed from the pupal case and kept them in isolation, ensuring their virginity (Fig 4). To arrive at a final collection of virgin females, these singly-housed individuals were sexed by hand. Even with this manual step, virgins were procured at a much higher (~3x) rate with MAPLE’s assistance than without it.

Thanks to MAPLE’s capacity to handle many individual flies in combinatorially complex experimental designs, we demonstrated that fly social behavior depends on sensory cues and found that individual differences in dyadic social behavior — as well as derived social interaction network topography — remains stable over days (Fig. 5). Dependence of fly social behavior on visual (Mery et al., 2009; Simon et al., 2012) and olfactory cues (Schneider et al., 2012; Billeter et al., 2015), and stability of social networks on short timescales (Schneider et al., 2012) are known, but long-term stability of social networks is a novel finding. This illustrates that MAPLE 1) can replicate known results without introducing significant behavioral confounds and 2) extend our knowledge by conducting complex, long-term experiments that are challenging for human experimentalists.

Technologies that automate experiments will particularly facilitate the phenotyping of individual animals, an approach that reveals underappreciated intragenotypic variability, and is likely to become increasingly prevalent as the philosophy of precision-medicine (understanding the individual as an individual) spreads throughout the biological sciences. Our group has shown that individual flies generally exhibit very different behaviors from one another even if they are reared in the same environment and have the same genotype, and that these differences persist across days (Kain et al., 2012; Buchanan et al., 2015; Ayroles et al., 2015; Kain et al., 2015; Todd et al., 2017). MAPLE allowed us to show that this extends to dyad-specific social interactions in flies (Fig 5). To our knowledge, this is the first demonstration that dyad-specific social interactions are persistent on days-long timescales in flies.

Another potential advantage of using MAPLE for fly experiments is the option to avoid anaesthesia. The Fly Dispenser releases awake animals into the MAPLE system, and these can then be moved between the FlyPlate and behavioral arenas using the multi-position loading ports we developed. Avoiding anaesthesia has multiple benefits, including reducing the distortion of behavioral (Bartholomew et al., 2015) and physiological measurements (Colinet et al., 2012), which, depending on the form of anaesthesia, can last for hours or days (MacMillan et al., 2017). Even seemingly benign manual aspiration can disrupt the expression of sensitive phenotypes (Trannoy et al., 2015). Automated, anaesthesia-free animal-handling thus has the potential to allow biological measurements that would be otherwise impossible.

MAPLE’s modularity means that the platform's versatility and capabilities will expand in the future. New experimental modules can be integrated into the system with only a modest coding effort. As examples, we are developing fly sink modules for culturing thousands of animals in the style of population cages, facilitating selection experiments and experimental evolution (long, grueling procedures begging for automation). We are developing a high-resolution imaging source/sink module into which flies could be deposited, imaged in dorsal and ventral views, and released back into experimental workflows. This device is inspired by the Fly Catwalk system (Medici et al., 2015) but can be loaded with a fly on demand, rather than relying on flies to enter the imaging chamber on their own. The images from this module will allow machine-learning based classification of morphological phenotypes, like sex and eye color, as well as quantification of fluorescent proteins; key genetic markers in flies.

In addition to conducting tedious or complex experiments, MAPLE’s integrated format affords the opportunity to feed back the results of phenotypic assays into fly handling tasks. For example, MAPLE could be used to perform artificial selection, identifying individuals with a specific behavior or phenotype and placing males and females together in wells of a FlyPlate, vials, or population cage. Likewise, multigeneration genetic crosses will be possible with our high resolution imaging module. Our current behavioral phenotyping measures use simple motion tracking, but behavioral assays employing thermogenetic or optogenetic stimulation, or sophisticated stimulus control could be implemented through custom modules (yet to be designed, but easily incorporated into the MAPLE ecosystem). Even the venerable forward genetic screen could be automated. Unsupervised learning algorithms have been used to automatically phenotype flies (Berman et al. 2014; Todd et al., 2017), and with the capacity to store and access large numbers of flies individually, MAPLE could identify and isolate outliers within a population without human intervention. We believe MAPLE is compatible with all such modern, automated approaches to fly experimentation, and brings automated animal-handling one step closer to the revolutionary potential achieved by liquid-handling robots.

## Methods

CAD files for MAPLE can be found at *https://github.com/FlySorterLLC/MAPLEHardware*. Control software for MAPLE including scripts for the experiments described here can be found at *https://github.com/FlySorterLLC/MAPLEControlSoftware*. Raw data and analysis scripts can be found at *https://zenodo.org/record/1119131#.Wj7SYlQ-eRc*. These materials are also available at *http://lab.debivort.org/MAPLE*.

### MAPLE Technical Specifications

MAPLE’s frame is a rectangular prism constructed from extruded aluminum struts (Misumi HFS-5 series in various sizes and lengths) and brackets (Misumi 5 series). The principal axes of the robot are Cartesian, that is to say linear and mutually orthogonal. The longest axis - designated the X axis - comprises two supported linear rails (IGUS Drylin AWUM-12), each with two housed bearings (IGUs drylin OJUM-6-12). The total of four housed bearings support a single, wide linear rail (IGUS WS-10-120) that is the Y axis. A carriage made of two aluminum plates sandwiching four more bearing blocks (WJ200UM-01-10) slides along the Y axis. Suspended from this carriage is an assembly that houses the three independent Z axis slides (IGUS SLN-D740679-2). We deliberately chose sliding bearings (as opposed to ball bearing slides) for the three axes to avoid noise and vibrations that might confound behavioral experiments.

The first of the three end-effectors is a small part manipulator. Made from an off-the-shelf vacuum cup connected to an air manifold, it can pick up and deposit lids or other small components from the workspace. A high-resolution digital camera and lens (The Imaging Source DFM 72BUC02-ML and TBL 9.6-2 C 3MP) is mounted to the second Z axis. Moving this Z carriage up and down focuses the camera. The third motorized Z slide holds a custom aspirator to move flies. Several short lengths of small diameter tubing (sections of needle tips, McMaster 75165A553 and 75165A682) are fixed approximately 4 mm inside a blunt, Luer-lock needle (McMaster 6710A61), forming a barrier to flies but allowing air to flow. A custom-molded silicone rubber boot surrounds the fly manipulator, forming a better seal against apertures in modules and improving fly handling (Fig S3).

Stepper motors (1.5 A NEMA 17 60 mm bipolar stepper motor, MNEMA17-60 from RobotDigg.com) mounted to the main frame drive two 6mm-wide GT-2 belts arranged in a CoreXY configuration (Moyer, 2012), and these belts drive the motion in the X and Y directions. Each Z axis slide is driven by its own stepper motor, integral to the slide assembly. The maximum speed for the X and Y axes is 200 mm/s. The Z axes top out at 83.3 mm/s. Limit switches (Omron SS-5) are mounted to the frame for each axis, providing repeatable end stops for homing.

Motion control, as well as the control of auxiliary devices such as the solenoid valves and the LED illumination, is handled by a Smoothieboard v1 PCBA, running custom Smoothieware firmware (included in our Github repository). G-code commands are sent from a PC connected via USB, interpreted on the Smoothieboard, and translated into electrical signals sent to each stepper motor. Scripts containing each experimental protocol, along with a common set of frequently-used subroutines, were written in Python 2.7 (or Matlab 2016a for BEECH) and executed on PCs running Windows 7.

### Modules

Modules were made by a variety of fabrication techniques, particular laser-cutting of acrylic. Vector outlines of module components were made in Autodesk Inventor or Adobe Illustrator and printed in CorelDraw. Acrylic components were joined together using Plastruct plastic weld. Please see Supplementary Online Methods for specific descriptions of the MAPLE experimental modules.

### Fly Lines

All experiments were performed using Canton-S (wild type), *Orco, NorpA*, or *w*^1118^ lines. Mutant lines were homozygous. We raised flies on standard cornmeal media (BuzzGro from Scientiis) under 12 h/12 h light and dark cycle in an incubator at 25°C and 70% humidity. Flies were anesthetized using carbon dioxide (CO_2_) and housed in vials of 15 to 20 flies, unless otherwise specified. Five days post-eclosion, flies were aspirated into individual wells in the FlyPlate using CO_2_ and used for experimentation after at least 2 hours of recovery.

### General experimental procedures

All experiments were conducted between 9AM and 9PM. Flies were loaded into individual arenas by MAPLE with arenas empty after 2 iterations loaded manually using an aspirator. Flies were later removed from their arenas in an identical fashion. FlyPlates, social arenas, and y-maze arrays were filled with 96, 162, and 81 flies, respectively. Flies were assayed using diffused white LED backlighting (Buchanan et al., 2015) in a temperature (23°C) and humidity (41%) controlled behavioral observation room. Fly movement was tracked for 1 hour. Fly tracks were analyzed using a custom MATLAB script. For longitudinal assaying, flies were moved back into the FlyPlate after phenotyping and allowed to feed and rest over night or for 1h at minimum.

Experimental procedures for specific experiments are provided in the Supplementary Online methods.

### Behavior measurement

Fly movement was tracked at 29.9 fps using a custom real-time MATLAB script interfacing with a Firefly MV FMUV-13S2C USB-camera. Tracks were analyzed in MATLAB. In total, 3% of the data were discarded because flies were immobile as determined by mean speed thresholds.

### Statistics

Unless otherwise specified, all confidence intervals were computed by bootstrapping the data associated with individual flies or individual fly dyads (in the case of social interaction measurements) 1000 times, using custom MATLAB scripts. One standard deviation of the bootstrap estimates was our estimate of the standard error of the estimate. P-values reported were adjusted for multiple comparisons using the Bonferroni correction where applicable, and asterisks reflect post-correction significance.

## Acknowledgements

We are grateful to Jim Morris for assistance developing the Smoothieware firmware, Matt Zucker for ongoing software and hardware advice, Chris Stokes for assistance with early prototypes, Yong-Li Dich for help with hardware development, and Kyobi Kakaria for consultations, fabrication assistance, and manuscript feedback. Jess Kanwal, Katrin Vogt, and Armin Bahl provided helpful manuscript feedback. Claire Guérin helped with bee colony box design and BEECH assembly. B.d.B. was supported by a Sloan Research Fellowship, a Klingenstein-Simons Fellowship Award, and the National Science Foundation under grant no. IOS-1557913. J.D.C. was supported by a Winslow Foundation research grant. D.Z. was supported by the NIH under Award Number R43OD023302. The content is solely the responsibility of the authors and does not necessarily represent the official views of the National Institutes of Health.

## Contributions

B.d.B. and D.Z. conceived the robot idea. T.A., D.Z. and B.d.B. developed MAPLE hardware. T.A. and D.Z. developed MAPLE software. T.A. conducted the scientific experiments, which were conceived by T.A. and B.d.B. J.D.C., D.Z., and B.d.B. developed BEECH. T.A. and B.d.B. performed the data analysis. All authors contributed to the manuscript.

## Online Methods

### Software

Every module has an associated Python 2.7 module class file (all MAPLE control software, including module class files, are available at https://github.com/FlySorterLLC/MAPLEControlSoftware). These files provide the 3d coordinates of key points on each module, including the sites of any ports or adapters through which flies are conveyed, as well as the z-clearance above the module needed so that the end-effectors do not collide with the modules in the workspace. These classes can be instantiated in Python files that represent workspace configurations *(Examples/ExampleWorkspace1.py)*. Beyond the module class files, there is a master MAPLE class file *(robotutil.py)*, which 1) establishes the communications connections between the experiment-coordinating computer and the motion control card and camera in MAPLE and 2) contains functions for all low-level robot operations, like returning to the 0,0,0 home position, moving to an arbitrary position, opening or closing the solenoid valves that control the vacuum flow in the end effectors, or acquiring a photo from the end-effector camera. There is also a file *(commonFlyTasks.py)* that contains subroutines for common usage tasks, like the combination of end effector movements, vacuum and air valve engagement, and Fly Dispenser serial commands required to retrieve a fly from the Fly Dispenser adapter on the workspace. Files of an additional type, experimental scripts, implement the actual experimental procedures (e.g., Fig S5; Movie S4). Each of these scripts load the class files for the modules used in their respective experiments, and procedurally calls the low- and mid-level functions of robotutil.py to implement each procedure. See Fig S6 for a schematic of the software architecture. Lastly, to increase the convenience of using MAPLE, we implemented a remote control system in which representations of the current status of experiments (e.g., text reports and digital images) were posted to a dedicated email account. This account could also receive MAPLE commands by email to remotely trigger experimental procedures.

### Modules

Phenotyping modules used to investigate affiliative behavior and locomotor handedness, i.e., social arena and y-maze arrays, measured 30 cm × 30 cm (Fig 1), and were fabricated from sheet acrylic using a laser cutter. Fly source modules were custom made (CO_2_ pad) or purchased from FlySorter LLC. Fly Dispenser dimensions are 22 cm × 15 cm, CO_2_ pad dimensions are 25 cm × 15 cm, and FlyPlate dimensions are 16 cm × 10 cm.

### Workspace

The workspace refers to a 100 cm × 28.2 cm × 7.5 cm volume that can be accessed by all end-effectors. The bottom of this volume is a clear acrylic floor with 5 mm holes organized in a grid (10 cm apart) and cable pass throughs for easy organization of individual modules. The 5mm holes can be used to affix an interchangeable acrylic plate (a “workplate”) to the acrylic workspace floor using nylon thumbscrews. Workplates have the locating brackets which define the positions of modules in a particular experimental configuration. Thus, every experimental script is associated with a physical workplate that locates the modules used in its experiment.

### Fly Dispenser

The Fly Dispenser isolates and outputs individual flies from an attached vial. Repeated knocking motion (which mimics the tapping gesture that people use to knock flies down in a vial) causes flies to fall from the vial into a funnel. At the bottom of the funnel, a pair of motorized, soft foam wheels acts as a valve, and a photo interrupter detects when one fly has passed by. The wheels are stopped, preventing other flies from passing through the valve, and an air pump transports the isolated fly out a tube. The dispensing process can be remotely triggered and monitored by Python scripts via a USB serial interface.

### Dispenser adaptor

MAPLE interfaces with the Fly Dispenser through the dispenser adaptor logistics module. The dispenser adaptor is a 4 cm × 1.5 cm 3D-printed ABS block with two 5mm diameter plastic Luer lock tube sockets attached on opposite sides. The Dispenser handpiece connects to the bottom side of the dispenser adaptor, while the MAPLE fly manipulator end effector aligns to the opening on the top of the adapter.

### FlyPlate

The FlyPlate is a modified 96-well plate positioned on a food tray. Each well in the FlyPlate follows the 96-well plate standard for bottomless wells (7 mm in diameter and 10.9 mm deep). Wells have a stainless steel mesh floor that allows feeding but prevents escape. The plate lid has x-shaped laser-cut openings over each well, cut into a flexible nylon mesh, that allow MAPLE’s individual fly manipulator (or a handheld aspirator) to penetrate to remove or deposit flies. The openings close back up once the aspirator/manipulator tip has been removed, keeping flies securely housed. Flies had free access to standard cornmeal diet on the food tray placed below. Food was replaced every 2 days to maintain adequate moisture and freshness and remove eggs and first instar larvae.

### Morgue

The morgue fly sink module is a 10 cm diameter × 5 cm deep laser-cut acrylic cylinder covered by a detachable lid that allows quick disposal of its contents and is equipped with an nylon mesh adapter (in the style of the well coverings of the FlyPlate) that allows MAPLE to deposit flies into soapy water or ethanol that traps and euthanizes them.

### Fly food vial

The fly food vial is a standard 2.6 cm × 10 cm vial equipped with a detachable lid that facilitates MAPLE fly depositing. A standard fly culture media vial can be placed into the fly food vial.

### Arenas and arrays

#### Social arena array

Arenas used for affiliative behavior and circling bias experiments are circular in shape with a 30 mm diameter and a height of 3 mm (Fig 5). Arenas are covered by a multi-position loading port — a rotatable clear lid with a 3.5 mm diameter opening through which flies can be deposited and removed (Fig S4). Two equal-sized semicircular compartments are formed by a 1.5 mm thick interaction barrier. Interaction barriers refer to individually laser-cut blocks that can be placed into corresponding openings in the middle of the circular arena, allowing separation of flies into individual compartments. Barriers were laser-cut from either clear or black acrylic and were designed to be either solid or open. Solid barriers were flat on the bottom. Open barriers had 4 ~0.25 mm horizontal channels connecting the two compartments (Fig 5; Movie S9). Open barriers presumably facilitate the exchange of odor cues between compartments. Barriers could be made of clear acrylic, facilitating visual cues, or black acrylic. A social arena array comprises 81 circular arenas, with 162 semicircular arenas in total.

#### Y-maze array

Locomotor handedness was assessed using y-maze arenas (Buchanan et al., 2015). Individual arms of the symmetrical Y-shaped mazes are 15.5 mm long and 120° apart. Arm ends are circular (5.2 mm) in shape, making it easier for flies to turn around and permitting loading and unloading flies via multi-position loading ports (Fig 3, S4). Arenas are covered with identical lids as those in the social arena array. A y-maze array comprises 81 y-maze arenas arranged equidistantly in a 9 by 9 grid. All parts described were manufactured from either clear or black acrylic and cut into shape using a laser cutter.

### Specific Experimental Procedures

#### Activity and speed MAPLE handling control experiments

Flies were loaded into a FlyPlate in accordance with general experimental procedures. MAPLE removed every second fly from its individual well and released it back after a 1 second delay 10 times in a row (48 flies total MAPLE-handled). This procedure was repeated 3 times so that every second fly was handled 30 times in total after 1 hour (Fig 3A-D). Flies were then monitored according to general experimental procedures. The preceding steps constituted 1 handling session. There were 5 handling sessions lasting 10 hours in total.

#### Manual vs MAPLE-handling control experiments

In the first control experiment (Fig 3E-H) lies were loaded into the y-maze array according to general experimental procedures. MAPLE loaded a random arena compartment (81 flies total) of each arena in the social arena tray to prevent dyad-neighbors influencing circling behavior. Flies were observed as described in general experimental procedures. Flies were discarded into morgue fly sink module after phenotyping.

#### Virgin-picking procedure

Ten male and 20 female CS between 5 and 7 days post eclosion are placed in custom dispenser-type vials. These custom virgin-picking vials are 10 cm long × 4 cm in diameter open cylinders with a bottom that attaches by press-fit. Standard cornmeal diet was poured in the bottom portion and allowed to cool down prior to fly introduction. Flies were anaesthetized using CO_2_ and placed in the vial. After 2 days of egg-laying in the incubator (25°C), parental flies were discarded and the vial was placed back in the incubator. After 9 days, the bottom portion of the vial containing the food was removed from the container. The food was discarded and the container washed and reattached to the vial. The vial now only contained animals that pupated on the sides of the open cylindrical portion of the vial (Movie S7).

The vial was then placed into the Fly Dispenser and MAPLE’s virgin-picking subroutine was engaged. Every 30 minutes, the Fly Dispenser attempted to dispense any eclosed flies while MAPLE aligns its fly manipulator end effector to the fly dispenser adapter. (Fig S5A). If a fly was successfully dispensed, MAPLE deposited it into a FlyPlate in the workspace. If at any point no fly is dispensed, MAPLE and the Fly Dispenser paused for a 30 minute waiting period before a new dispensing attempt was made. Over 3 days, MAPLE continued to load newly eclosed flies into individual FlyPlate wells until pupa were exhausted (Movie S9). FlyPlate wells containing multiple flies were manually emptied with an aspiratory. The remaining flies were manually anesthetized using CO_2_ and their sex assessed under a dissecting microscope. After sexing, flies were returned to their individual wells. The FlyPlate fly source module including food tray was removed from the workspace and placed inside a sealed plastic container. After 10 days, the food tray was examined for larvae, eggs, and newly hatched flies. When none were found, the single flies MApLe placed into individual FlyPlate wells were considered virgins.

#### Virgin-picking MAPLE/manual comparison

To allow a fair head-to-head comparison between manual virgin collection and MAPLE-assisted virgin collection, our approach was to start and end both procedures in identical circumstances (bottles collecting flies, and vials containing virgin females, respectively). Two standard bottles containing 10 male and 20 female CS flies 5–7 days post eclosion were allowed to mate and lay eggs for 3 days. After 3 days, flies were removed and egg- and larvae-containing agar was transferred with a spatula among 5 standard vials and 1 custom dispenser-type vial. The amount of virgin females automatically picked by MAPLE was assessed according to the general virgin-picking procedure described above. Total time required was computed as the total time required for every step of the process, including preparation, manual sexing, and cleanup. Virgin female count, virgin female ratio, and time required for manual virgining was assessed by twice daily (9AM and 9PM) manual virgin-picking from 5 virgin-producing standard culture vials.

#### Social interaction paradigm validation

Social arenas were prepared by manually inserting the appropriate interaction barrier type into the arena array prior to introducing flies. Flies were loaded into social arena arrays according to general experimental procedures. To minimize behavioral confounds caused by disparate loading times, one social arena compartment (i.e., all the left compartments) was filled first. This ensured that flies in social arenas loaded earlier were allowed minimal additional time to familiarize with or habituate to their dyad-neighbor. Phenotyping arrays were moved into behavior-recording boxes according to general experimental procedures. After assaying, flies were manually removed from trays and discarded.

#### Social behavior day-to-day persistence (Scheme 1)

Flies were loaded into social arena arrays, assayed, and deposited into FlyPlates after phenotyping according to general experimental procedures. Dyads were randomly determined on the first day. Flies were phenotyped once per day. Fly identity and dyad composition was maintained throughout 6 assaying days. Compartments and arenas were loaded in a randomized fashion each day. On the 7th day, Canton-S flies in the open-clear interaction barrier condition were randomly placed in social arena compartments to form physically shuffled dyads to complement computational shuffling for resampling statistics.

#### Social behavior persistence (Scheme 2)

Flies were loaded into social arena arrays, assayed, and deposited into FlyPlates after phenotyping according to general experimental procedures. Each fly was randomly assigned to be part of 10 dyads on the first day. Flies were assayed 6 times per day for 7 days. Fly identity was maintained throughout the experimental duration. In total, each dyad was phenotyped 4 times. We averaged behavioral measures across the first and last 2 phenotyping sessions.

#### Social Interaction Network (SIN) persistence (Scheme 3)

Flies were born in the Fly Dispenser and deposited into FlyPlates according to general virgin-picking procedure. Two groups of 10 flies each were assayed and deposited back into FlyPlates after phenotyping according to general experimental procedures. Flies were assayed 6 times per day for 3 days to exhaust each possible dyad combination twice. Fly identity was maintained throughout the experimental duration.

### Social Interaction Analyses

#### Interactivity index

A dyad’s interactivity index was defined as the correlation of dyad-neighbors’ distances to the interaction barrier over the entire experimental duration (Fig. 5 B-D). Distance was defined as the euclidean distance between a fly’s centroid and the closest side of the interaction barrier.

#### Coincidental approaches

We defined a coincidental approach as an interval in which both flies in a dyad were located within 1 body-length (3 mm) of the interaction barrier on the same frame. This definition of social interaction yielded qualitatively similar results to the interactivity index (Fig S7). Distance to the barrier was defined as the euclidean distance between a fly’s centroid and the nearest side of the barrier. A coincidental approach was scored as a single event irrespective of its duration. For each subsequent coincidental approach to be valid, at least one fly was required to leave and re-enter the 3 mm zone. Coincidental approaches were normalized for dyad mean speed over the experimental duration. Dyad mean speed was the grand mean of both dyad-neighbors’ mean speed.

#### SIN connection threshold

In the graph representation of social interactions, edges between flies in the network were retained if the absolute value of their interactivity index was greater than the average of the absolute value of the first and fourth quartile of all dyads’ absolute interactivity indices. The same threshold was applied to both repetitions of the SIN measurement (Fig 5J).

## Supplementary Materials

### Contents

- 9 Supplementary Figures with Captions
- 8 Supplementary Movies with Captions
- Supplementary Document 1: MAPLE Bill of Materials
- Supplementary Document 2: MAPLE Build
- Instructions

### Supplementary Figures and Captions

**Figure S1.**
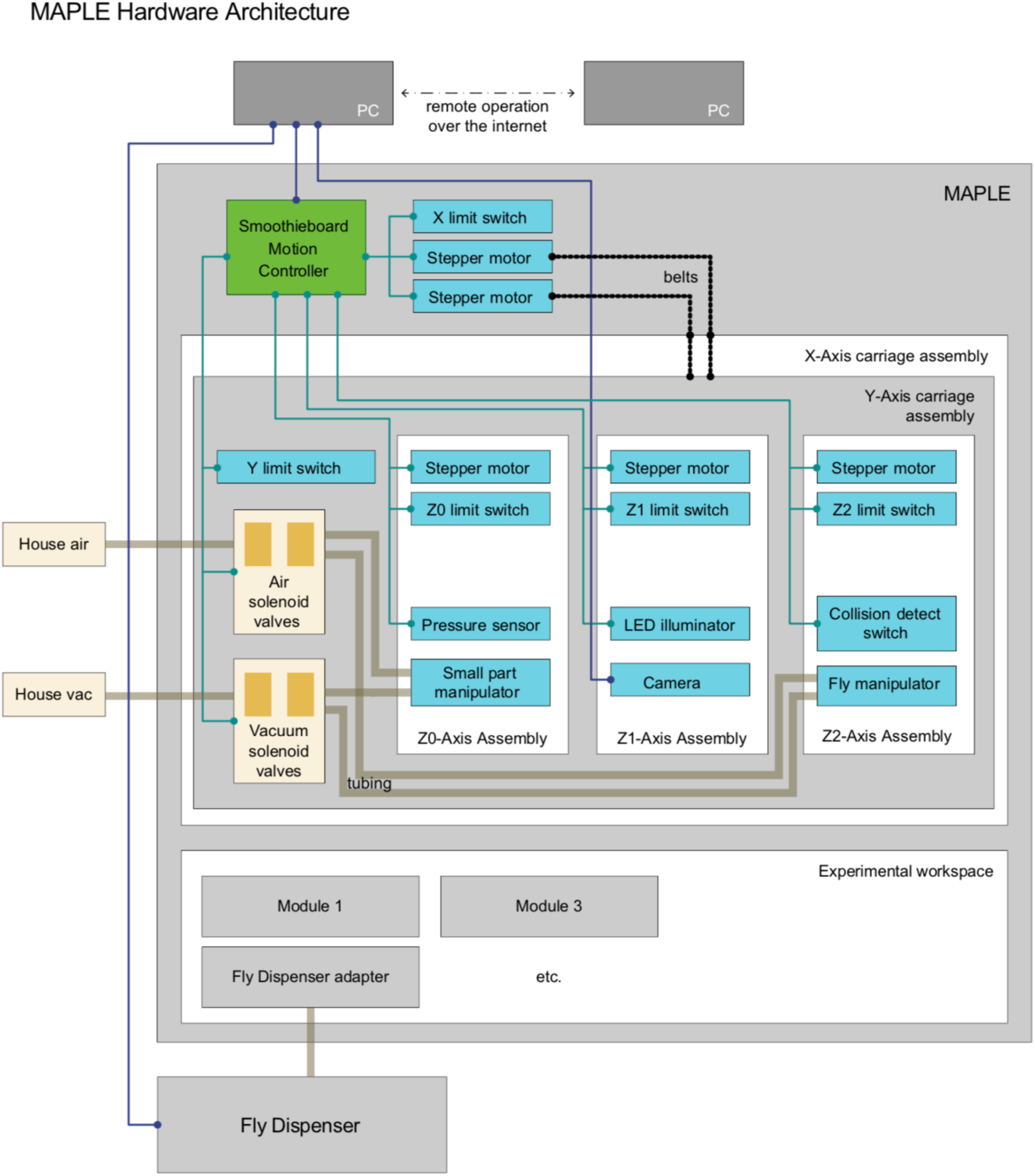
MAPLE hardware architecture. —MAPLE has two major components. First, animal-handling is mediated by a 5-axis Cartesian robot with three end effectors on independent z-axes. Two of these, the Small part manipulator and the Fly manipulator use positive and negative pressure pneumatics to manipulate module components like lids and individual flies, respectively. Secondly, handled animals move through an experimental workspace which contains reconfigurable experimental modules, and can interface with devices outside of MAPLE, e.g., the Fly Dispenser, using adapter modules.

**Figure S2.**
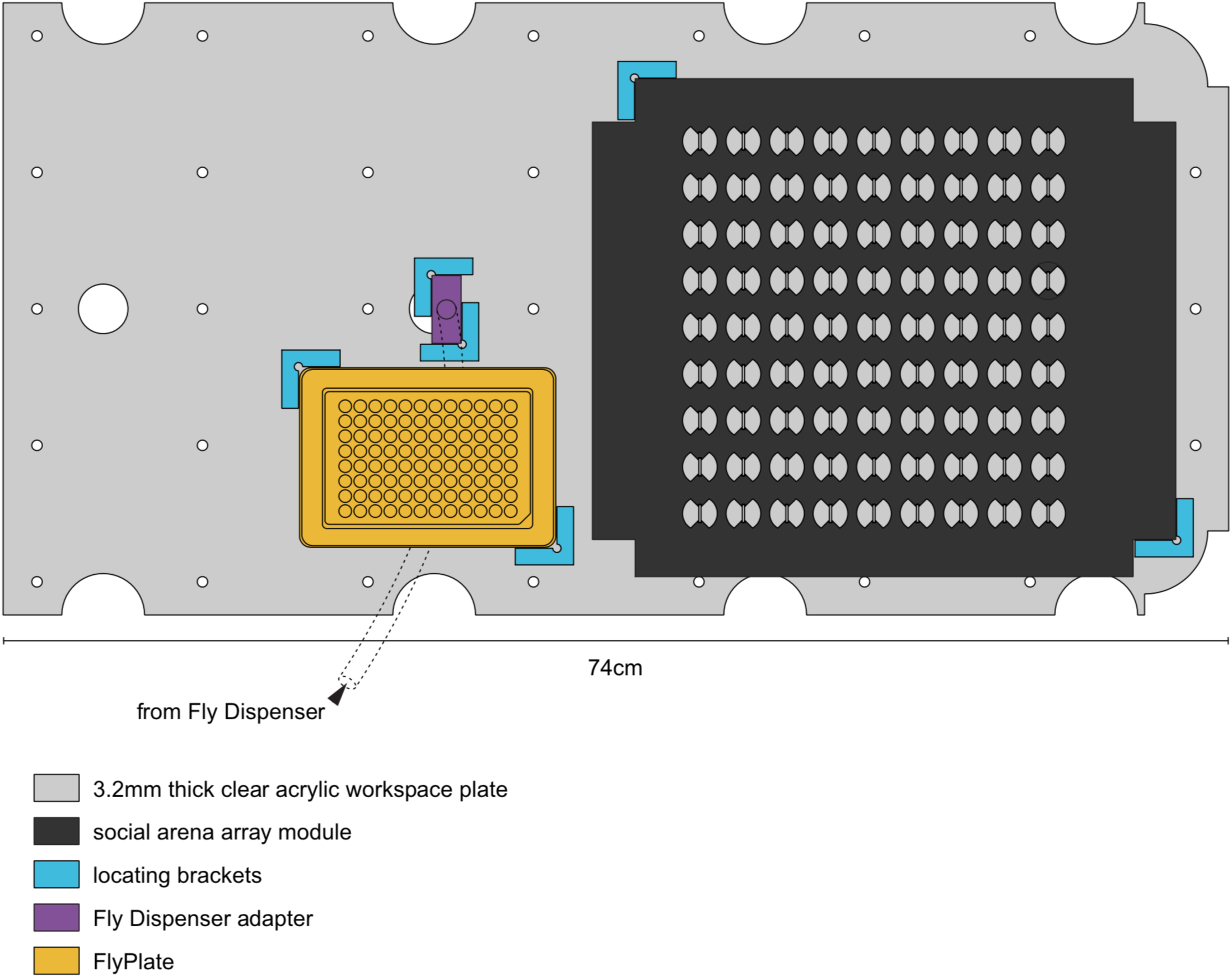
Example MAPLE experimental workspace. —To-scale illustration of the MAPLE experimental workspace. The workspace plate contains cable pass-throughs (large circular cutouts) and evenly spaced peg holes for registering it with the MAPLE floor and frame. This example workspace is configured for a social arena array experiment, wherein flies can be brought into MAPLE from the Fly Dispenser, through the Fly Dispenser adapter, into the FlyPlate, into the social arena array for behavioral phenotyping, and then potentially back into the FlyPlate, as we did in the longitudinal social interaction network persistence experiments (Fig 5G-J).

**Figure S3.**
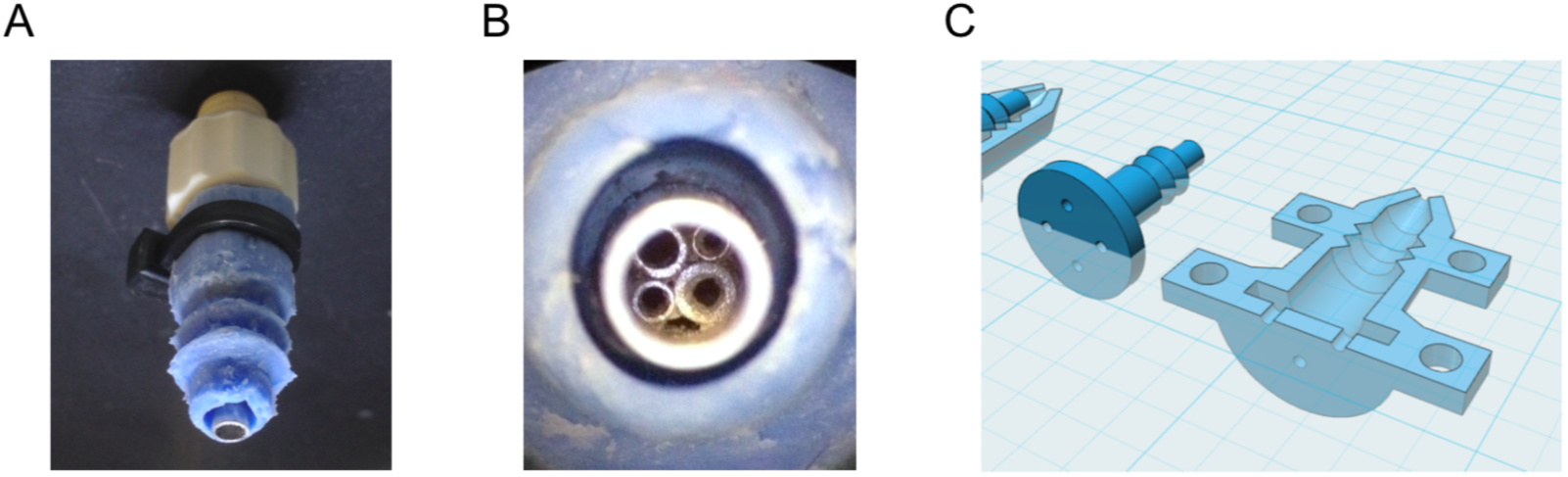
Fly manipulator end effector details. —A) Close-in photograph of the fly manipulator end effector, showing the luer socket fitting (white), pneumatic opening (steel, bottom) and bellows-style silicone cuff, which provides a better seal against multi-position loading ports or FlyPlate well tops for more effective removal of flies in compartments (blue). B) Macro view of the pneumatic opening, showing small diameter tubes inserted into the fly-manipulator tip to provide a surface against which flies being sucked into the manipulator would be caught, without greatly impeding airflow. C) CAD models of the 3D-printed mold that was used to make the silicone cuff.

**Figure S4.**
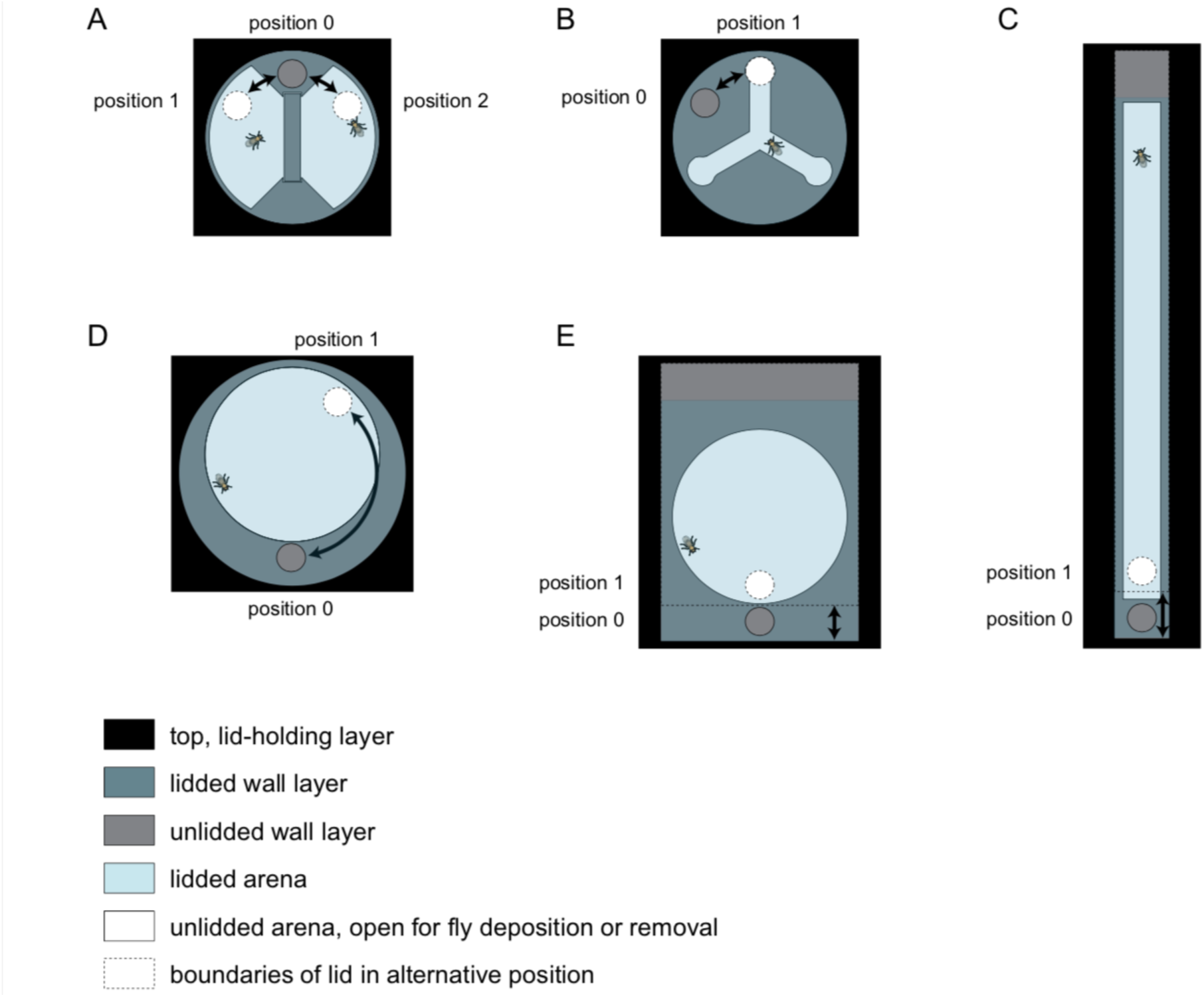
Multi-position loading ports. —Illustrations of arena lids that can be rotated or slid to position an opening over a behavioral arena, or over a portion of inaccessible wall, corresponding to configurations that are, respectively, open or closed for MAPLE loading or unloading. Arenas shown are: A) social arena, B) Y-maze arena, C) linear (Drosophila Activity Monitor style) arena, D) circular arena with rotating multi-position loading port, and E) circular arena with linearly sliding multi-position loading port.

**Figure S5.**
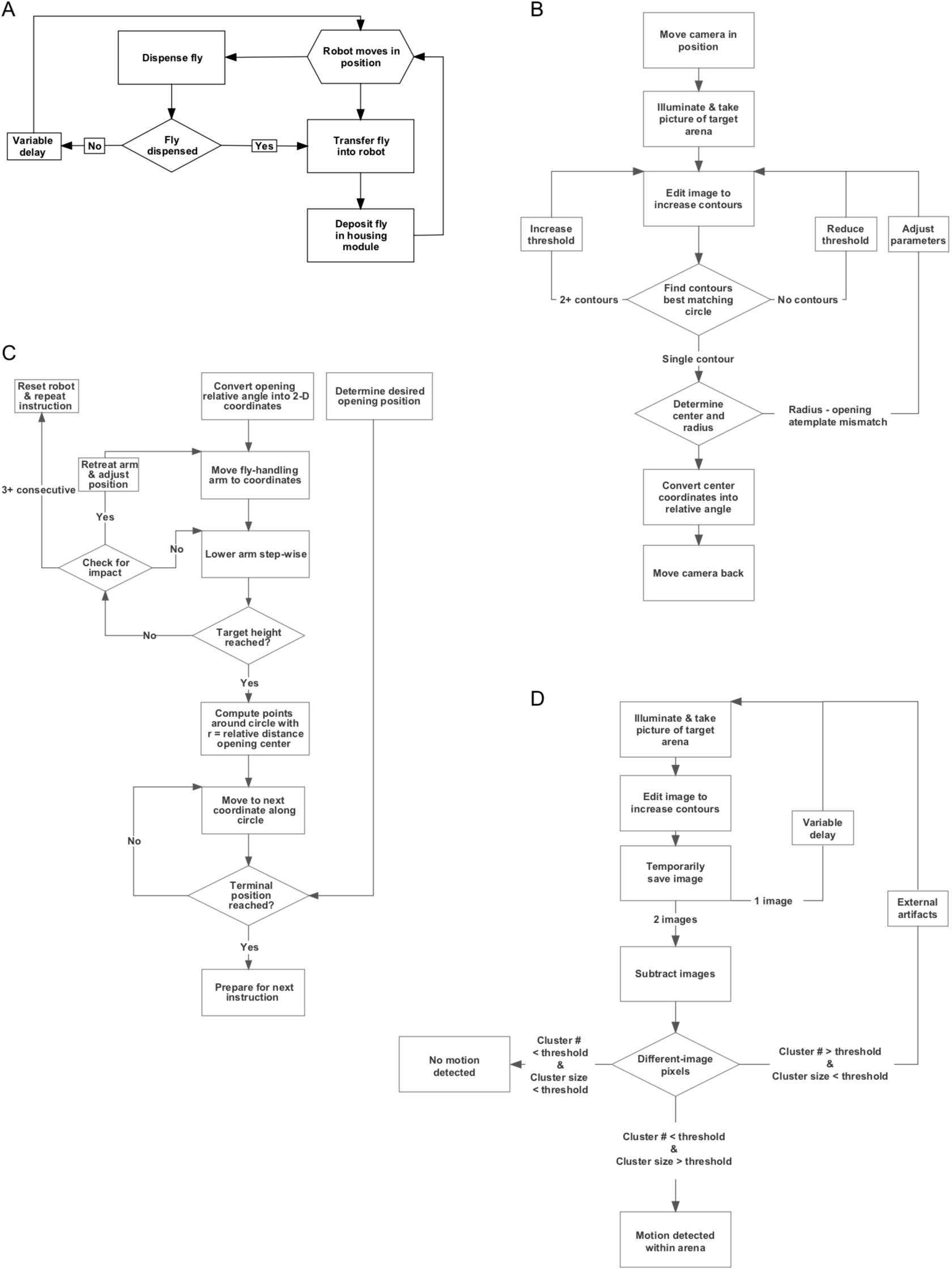
MAPLE procedure flowcharts. —A) Flowchart for a high-level MAPLE experimental procedure, the collecting of virgin females in a FlyPlate from the Fly Dispenser. High level functions such as this one depend on lower-level procedures, such as those illustrated in subsequent panels. B) Flowchart for detecting the opening of multi-position loading ports (Fig S4) using machine vision. C) Flowchart for using the fly manipulator end effector to put a multi-position loading port into the open configuration. D) Flowchart for using the camera end effector to detect if there is a moving fly within a behavioral arena.

**Figure S6.**
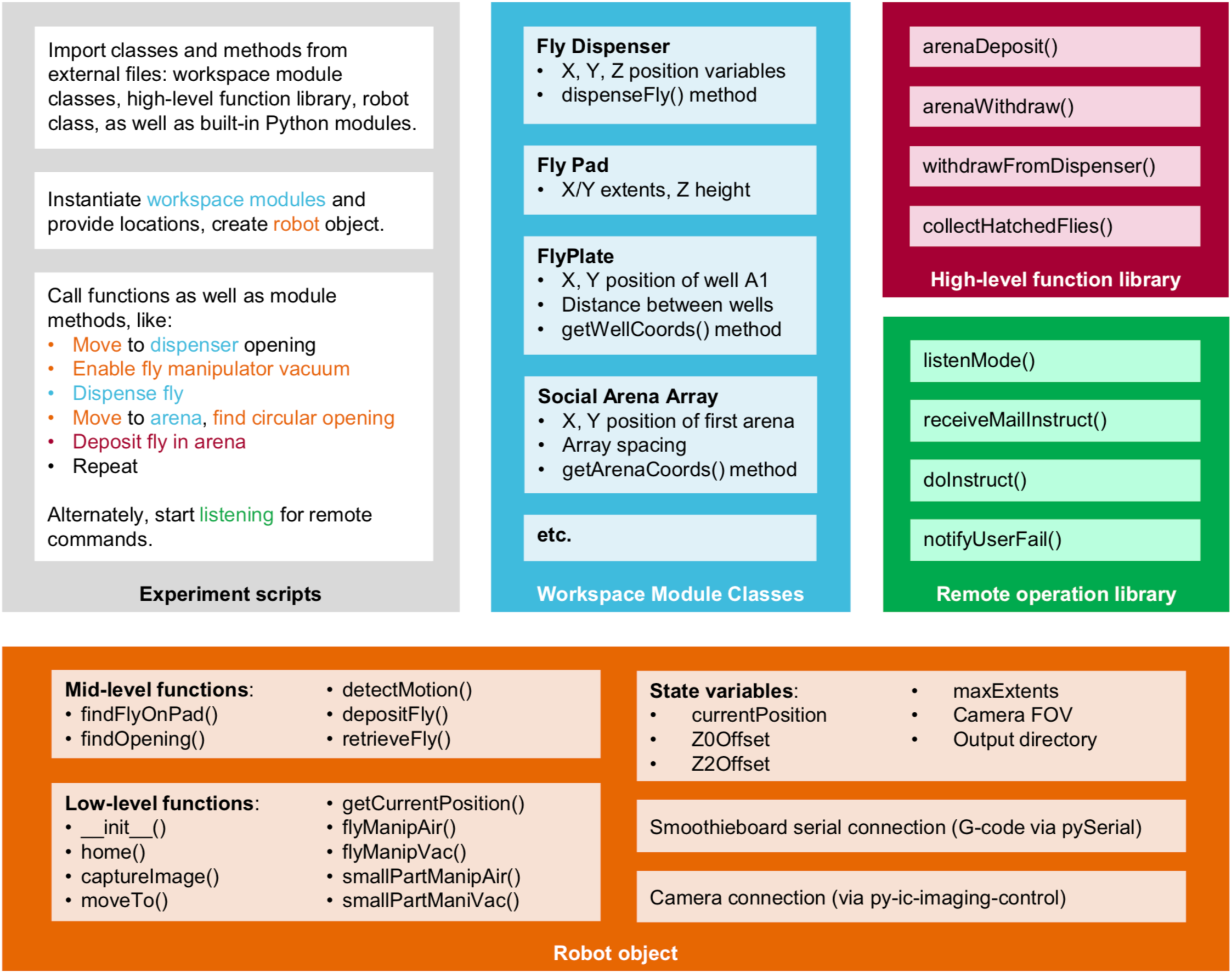
MAPLE software architecture. —Schematic of the organization of MAPLE software into experimental scripts, module class files, a high-level function library, a remote operation library, and the robot object which harbors functions for low-level robot operations. The generic experiment script indicates with color how each of these file types is called to generate a useful MAPLE procedure.

**Figure S7.**
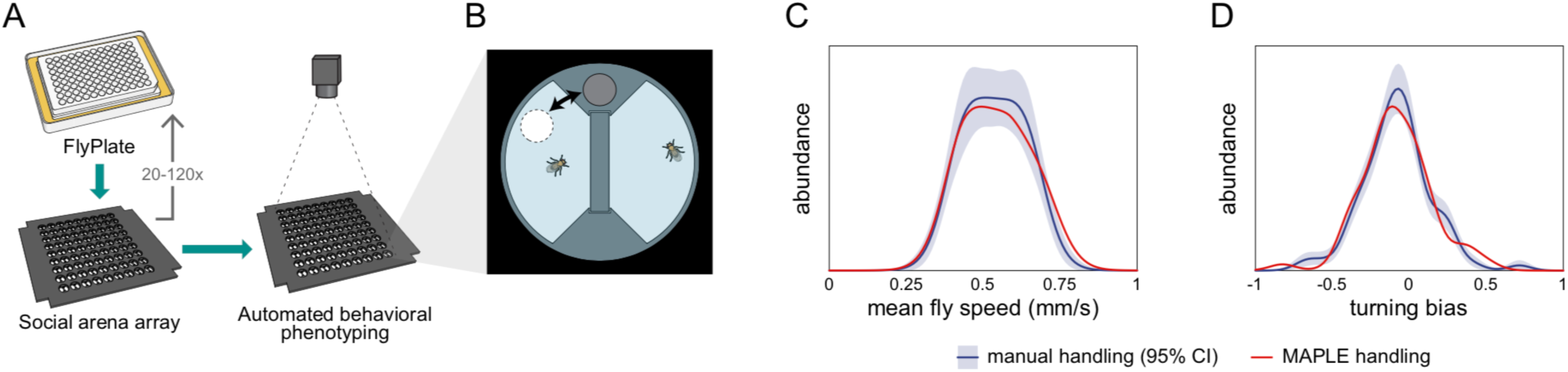
MAPLE-versus manual-handling in the social arena. —A) Experimental procedure by which MAPLE loaded flies from a FlyPlate into a social arena array for automated behavioral phenotyping in a back-lit imaging box. B) Flies were inserted into behavioral arenas via multi-position loading ports. C) The distribution of mean fly walking speeds in these arenas was statistically indistinguishable between MAPLE- and manually handled flies. *p* = 0.99 by KS-test. Distributions are determined by KDE. Red indicates MAPLE-handled flies. Dark blue is manually handled flies; light blue area is the 95% CI of the manually-handled KDE as determined by bootstrap resampling. D) The distribution of mean fly circling biases (tendency to turn clockwise or counterclockwise in their respective semi-circular arena compartment) was statistically indistinguishable between MAPLE- and manually handled flies, *p* = 0.99 by KS-test.. The circling bias metric was defined as ^ score (Buchanan et al., 2015), which measures the average circumferential component of motion, with −1 indicating purely clockwise motion, +1 purely counterclockwise, and 0 equal portions clockwise and counterclockwise motion or inward/outward motion. Colors as in (C).

**Figure S8.**
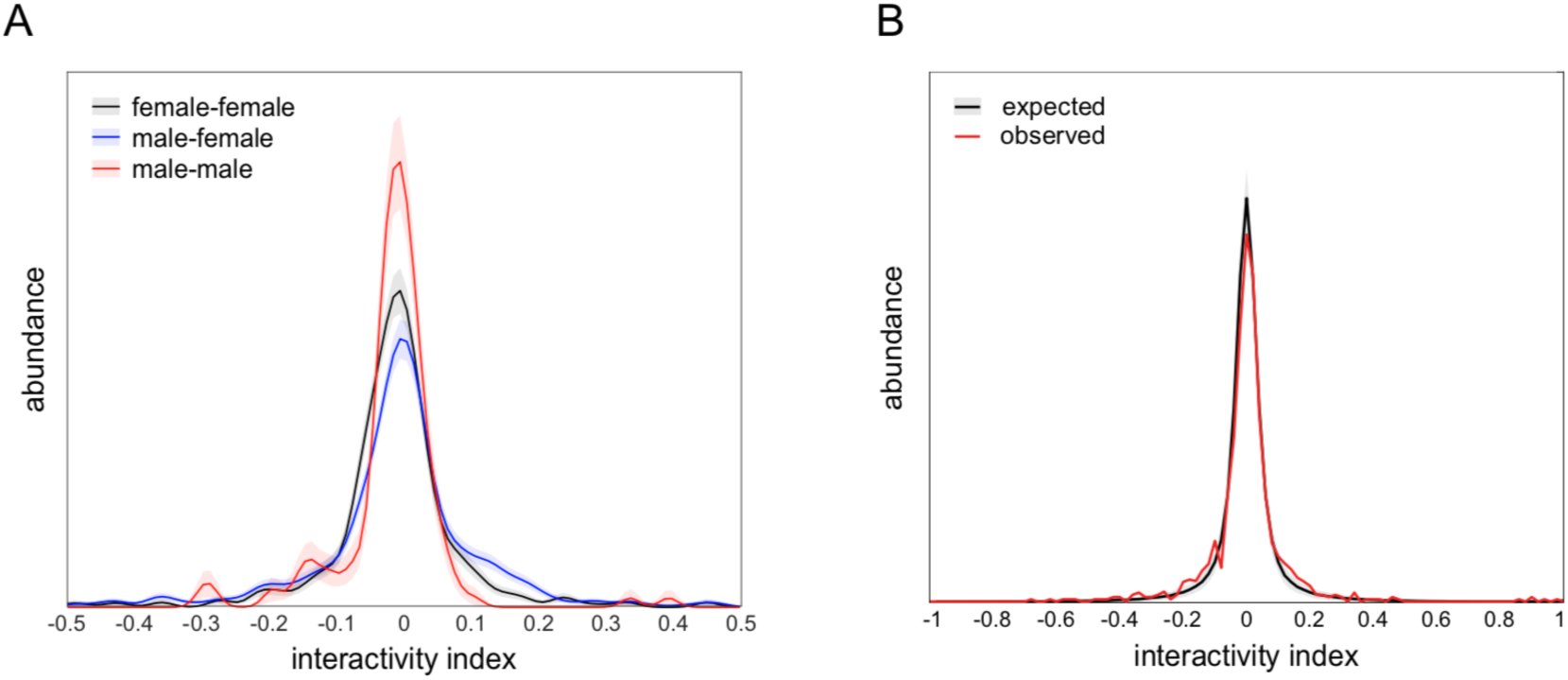
Interactivity index controls. —A) Interactivity index distributions of female-female (black), male-female (blue), and male-male (red) dyad compositions, visualized using kernel density estimation. Female-female distribution is not significantly different from male-female (*p* = 0.66 by KS-test) or male-male (*p* = 0.14 by KS-test) distributions. Male-female distribution is significantly different from male-male distribution (*p* = 0.0094 by KS-test). Distributions comprise 396, 474, and 132 dyads, respectively. Shaded regions indicate +/-1 standard error of the distribution estimate as determined by bootstrap resampling. B) Observed (black) and expected (red) distributions of interactivity index from male-female and female-female dyads (*n* = 870). The overdispersion of the observed distribution (standard deviation = 0.13) over the expected distribution (standard deviation = 0.096) indicates that dyads are exhibiting more strongly positive or negative interactivity indices than expected under the null hypothesis in which interactions between dyad partners are independent (F = 37.3, p < 0.0001 by Levene’s test; *p* < 0.0001 by KS-test; *p* < 0.0001 by *χ*^2^ test of variance). This null hypothesis was simulated in a resampling procedure by shuffling dyad partner identity and recomputing the interactivity index. Shaded region indicates +/-1 standard error of the distribution estimate as determined by bootstrap resampling. Observed distribution comprises 870 dyads. Bootstrapped shuffled dyad distribution was resampled 1,000 times.

**Figure S9.**
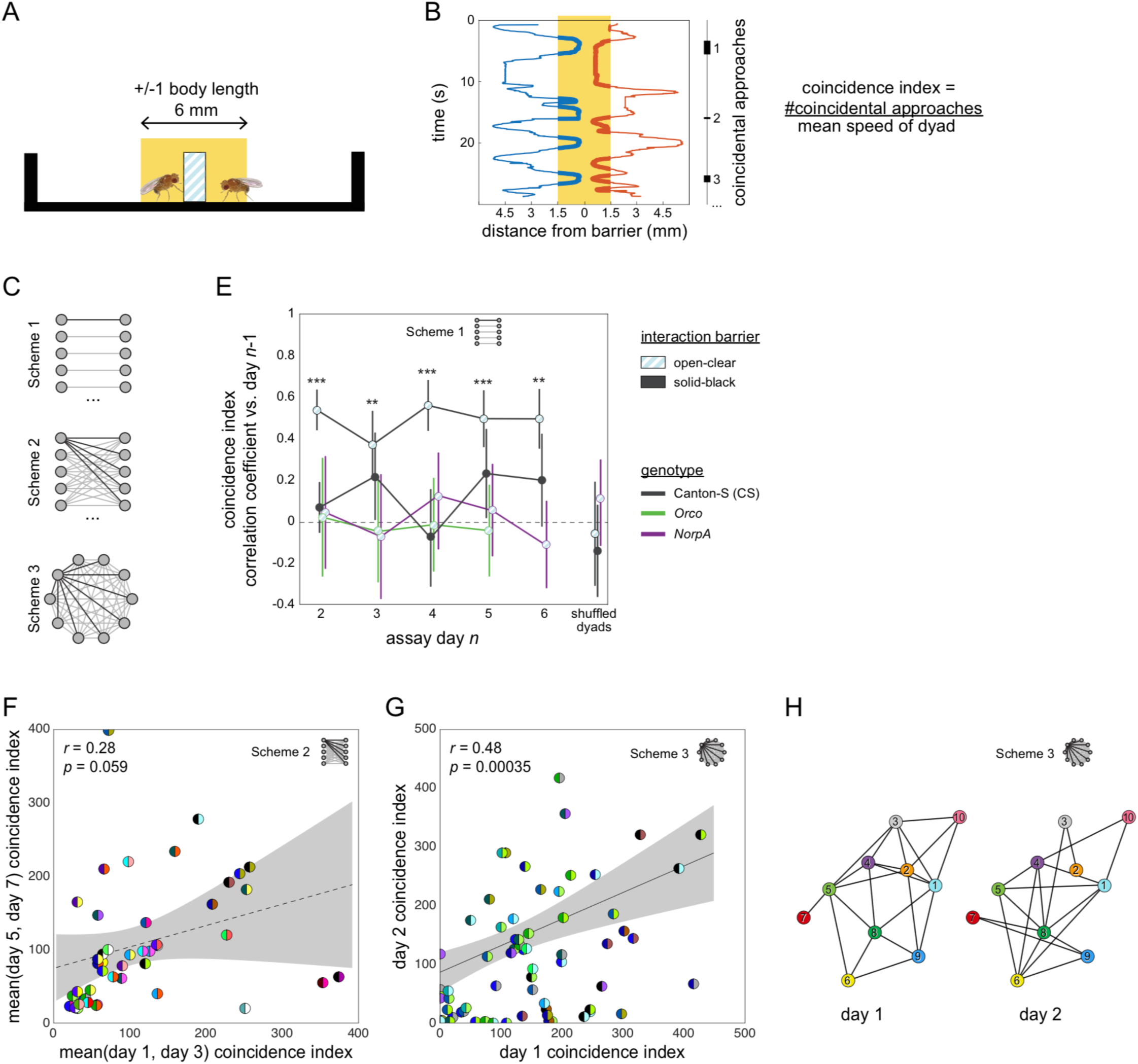
Social interactions and their persistence as measured by the coincidence index metric. —A) Schematic side view of a social arena showing in yellow the zone in which flies are scored as being at the barrier. B) Position (x-axis) vs time (y-axis) of two flies interacting in a dyad. Thick lines indicate timepoints when the flies are in the at-barrier-zone (yellow). Black bars at right indicate intervals in which both flies are at the barrier. Each of these intervals represents one “coincidental encounter”. Coincidence index is defined as coincidental encounters normalized for average dyad speed. C) Diagram of pairing schemes employed to form dyads. D) Pearson correlation coefficient between dyad coincidence indices measured on successive days, over 6 days. Dyads were formed according to scheme 1. Point patterns denote barrier type, line colors denote genotype. Error bars correspond to +/- 1 SEM. Asterisks denote significant one-sample two-tailed z-tests. E) Scatter plot of coincidence index of Canton-S (wild type) flies in arenas with open-clear barriers from measurements made across 2–4 day intervals. Dyads were formed according to scheme 2. Gray area is 95% CI of the linear regression line. Circles represent dyads; semi-circles denote individual flies forming a dyad; colors denote fly identity. Pearson correlation coefficient is not statistically significant, *r*(45) = 0.28, *p* = 0.059. F) As in (E) for two groups of 10 Canton-S flies forming 45 possible dyads each according to scheme 3 and tested on successive days, *r*(68) = 0.48, *p* = 0.00035. G) Visualization of a Social Interaction Network (SIN) of 10 Canton-S flies (45 dyads) in arenas with open-clear barriers. Connections denote dyads exhibiting coincidence indices greater than the average values of the 1st and 4th quartiles (threshold: 223). Threshold was identical for day 2. Colors and numbers indicate fly identity on both days.

### Supplementary Movie Captions

**Movie S1** (https://youtu.be/BMLy9QbgMLY) — *MAPLE in use* — MAPLE is situated on a standard experimental bench. The user controls its behavior through an attached PC. Real-time video shows that MAPLE moves at speeds similar to human fly experimentalists.

**Movie S2** (https://youtu.be/wxEgbYfif_M) — *Motion VR view from within MAPLE during an experimental procedure* — Time-lapse view from a wide-angle camera mounted on the x-axis assembly of MAPLE during a procedure to move flies from a 96-well plate into social arenas, providing a clear view of the action of all three z-assemblies and end effectors. Moments when MAPLE flashes multiple times successively in the same position reflect an algorithm to vary the exposure time for image acquisition to detect the opening of a multi-position loading port. The viewing angle can be adjusted during playback in certain video viewers like VLC 3.0 or the YouTube viewer.

**Movie S3** (https://youtu.be/bgtJA46egho) — *MAPLE can operate autonomously* — Wide-angle movie acquired by mounting the camera on the y-assembly as MAPLE conducts a social arena array loading procedure. After starting the protocol, confirming that the camera stays attached and trimming some zip-tie tails from the camera mount, the user can walk away while MAPLE continues the experiment.

**Movie S4** (https://youtu.be/F9KdnkGfkhI) — *Motion VR view from the MAPLE workspace floor* — Time-lapse view from a wide-angle camera mounted on MAPLE’s experimental workspace plate during a procedure to move flies from a 96-well plate into social arenas. Moments when MAPLE flashes multiple times successively in the same position reflect an algorithm to vary the exposure time for image acquisition to detect the opening of a multi-position loading port. The viewing angle can be adjusted during playback in certain video viewers like VLC 3.0 or the YouTube viewer.

**Movie S6** (https://youtu.be/crIb4ZfecYQ) — *Motion VR view from within a bumblebee colony in BEECH* — Wide-angle movie acquired by mounting the camera inside a bumblebee colony box and placing into the BEECH platform for imaging. BEEtag spatial barcodes are visible on the backs of individual bees. Once BEECH begins to move, its camera and IR illuminator effectors periodically come into view above the nest chamber (e.g., at 3:05; in this movie the normally visible-opaque next chamber roof has been replaced with clear acrylic. The viewing angle can be adjusted during playback in certain video viewers like VLC 3.0 or the YouTube viewer.

**Movie S5** (https://youtu.be/kDzjep15lAQ) — *BEECH recording behavior of multiple bumblebee colonies* — Timelapse of a real BEECH experiment in which the camera end effectors are moved between bumble bee colony boxes to record brief videos in each successively.

**Movie S7** (https://youtu.be/rfgrX_mQdmw) — *MAPLE-assisted virgin-collecting procedure* — Animation illustrating the procedure MAPLE can be used to more efficiently collect virgin female flies. See also Fig 4.

**Movie S8** (https://youtu.be/hntOpuU0AB4) — *Fly behavior in social arena assay* — Six social arenas are seen with a fly loaded into each of the 12 semicircular compartments. These compartments are separated by open-clear barriers which are made of clear acrylic with horizontal channels allowing flies to both see and smell each other.

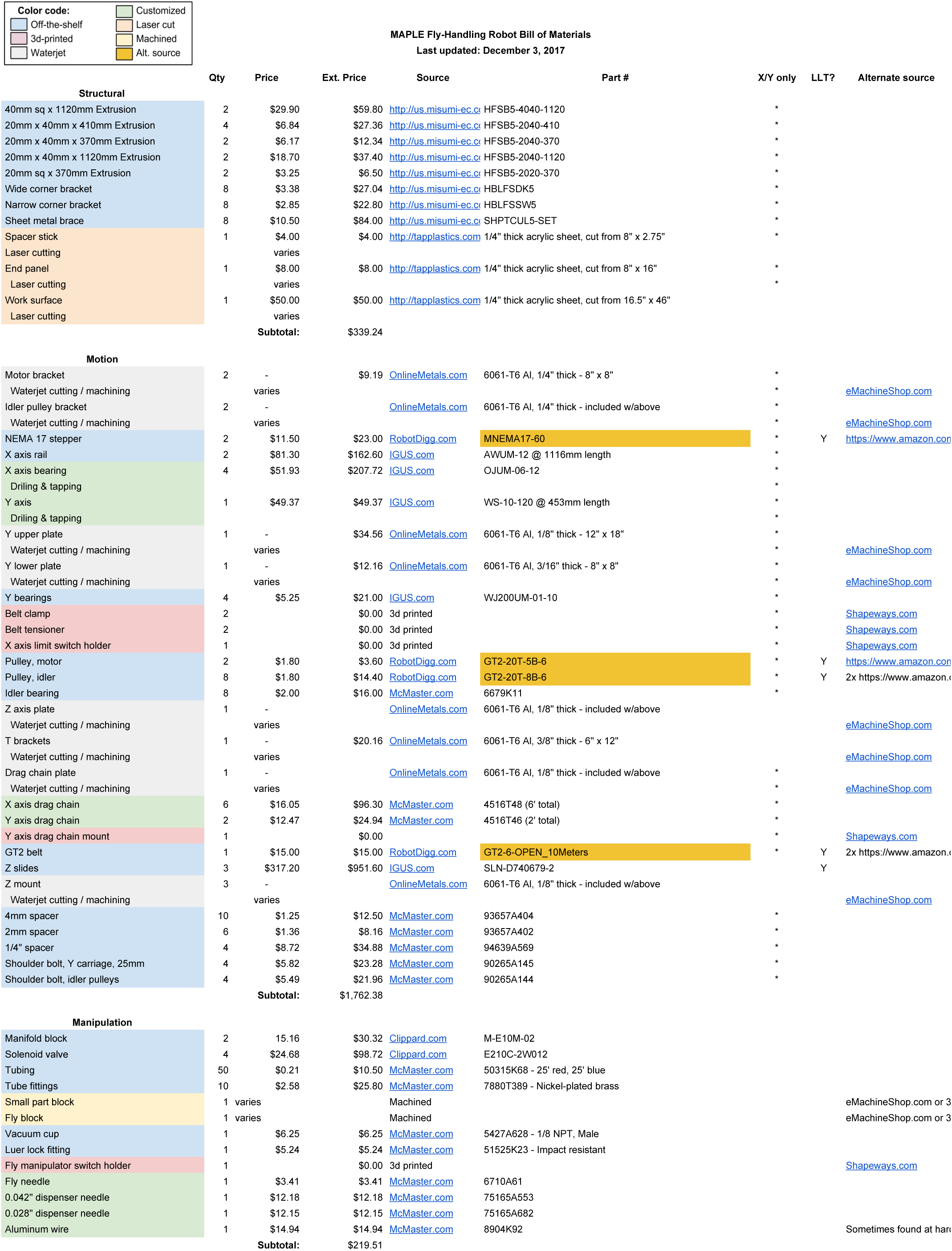

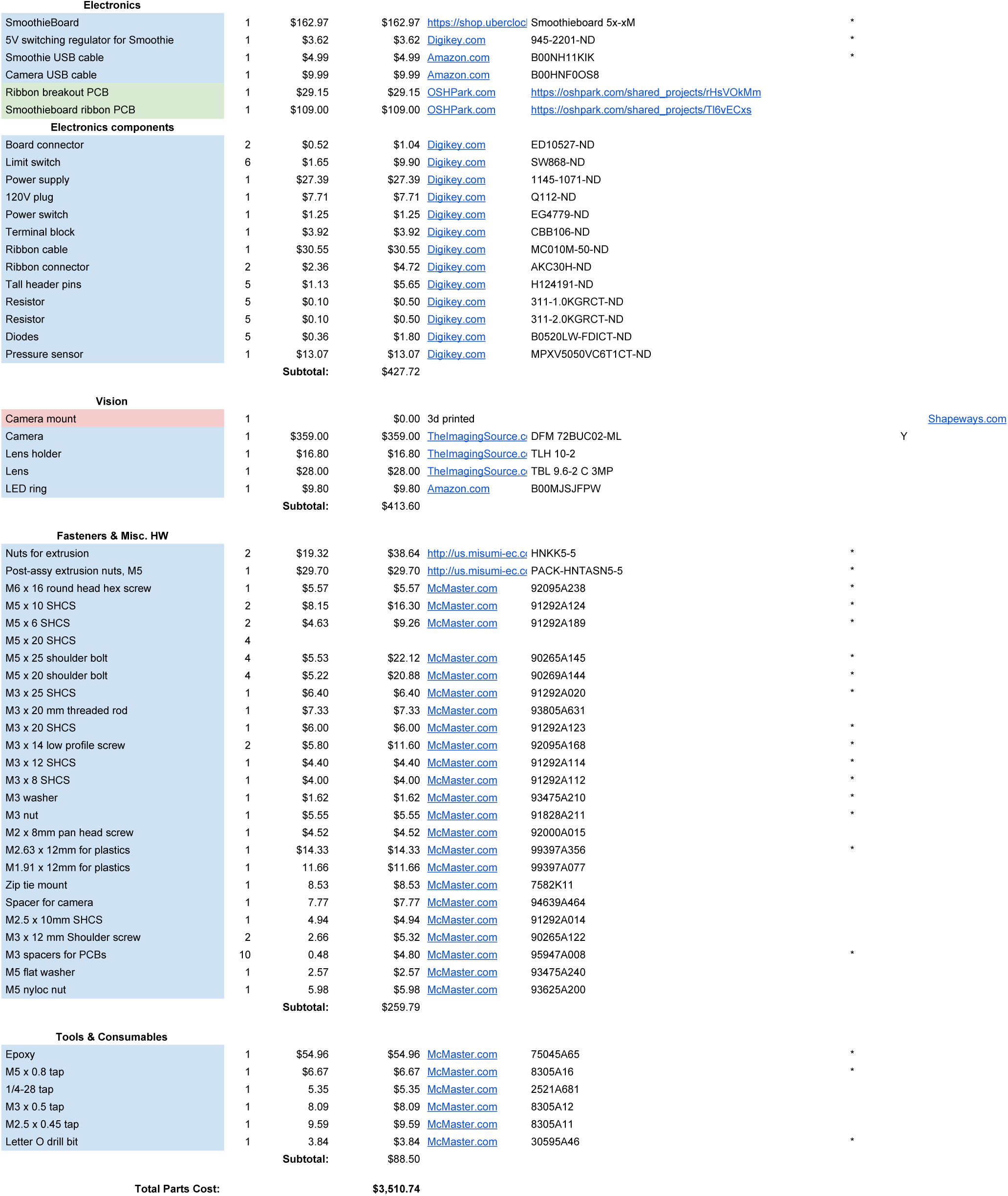

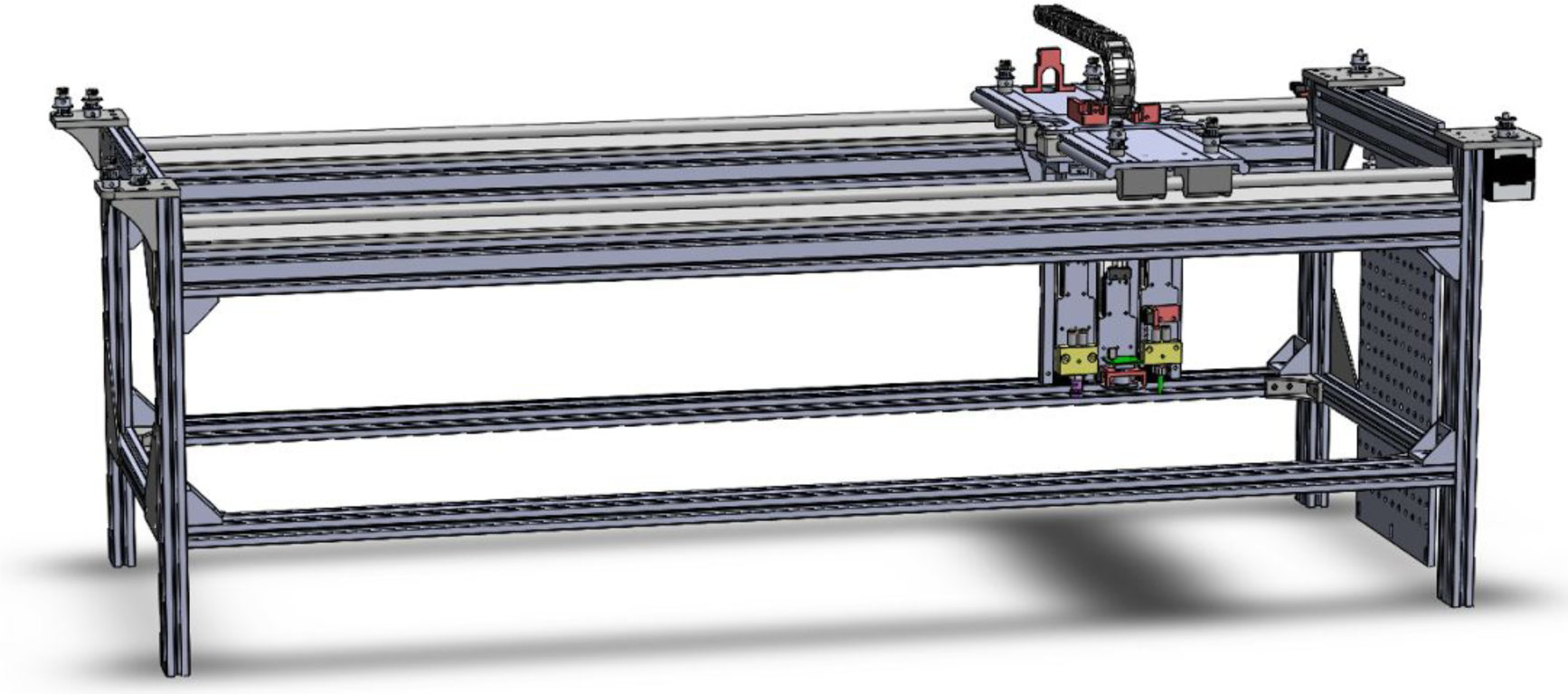

## MAPLE Build Instructions

Dave Zucker, <u>FlySorter LLC</u>

### Change Log

2016–12–15 - Version 2.0 - Initial build instructions

2017–07–27 - Version 2.1 - Doc updates (no design changes)

2017–11–25 - Version 2.2 - More details (no design changes)

MAPLE Build Instructions

Change Log
Notes
Documentation Sources & License
Pre-Assembly

Aluminum Extrusions
3D Printed Parts
Silicone Rubber Boot
Machined Aluminum Parts
Wateriet Aluminum Parts

Secondary operations:
Laser Cut Parts
X Axis Rails
X Axis Bearings
Y Axis
PCB Assembly

Smoothieboard
Smoothieboard-Ribbon PCB
Ribbon Breakout PCB
Frame Assembly
Y Axis Assembly
Assembling the Z Axes

Z0 - Small Part Manipulator
Z1 – Camera
Z2 - Fly Manipulator
Complete Z Assembly
Wiring & Tubing

Threading the Drag Chains

Large Drag Chain
Small Drag Chain
Power Supply
Smoothieboard
Z Assembly Air & Vacuum Tubing
Firmware
Alignment & Belt Tension
Testing

## Notes

These instructions will guide you through assembly of the MAPLE fly handling robot.

Before you begin, please note:

- Before you order parts or start 3D printing, read through this document thoroughly and make sure you understand what’s required.
- We assume familiarity with and/or access to a machine shop, hand tools, 3D printer, soldering station, and more. If you don’t have access to a 3D printer, there are online services that can provide parts, such as Shapeways.com, 3dhubs, and more. Similarly, eMachineShop.com will fabricate parts from CAD. But with that said, you will have to get your hands dirty at some point during this build, so it may not be for everyone.
- While we have made every effort to ensure the documentation is complete, we cannot be responsible for errors (though of course we will gladly try to help).
- The bill of materials for the MAPLE robot can be found <u>here</u> (https://docs.google.com/spreadsheets/d/1WS5JZ8TgN1G7WdbVoJIBsS1zV8HpRZNDN9ckr8WF8/edit?usp=sharing).
- Unfortunately, many of the parts (particularly fasteners) are sold in packs of 25, 50 or more, even if the assembly only uses a few. At this point, neither FlySorter nor the de Bivort Lab sells a ready-made kit of parts. The costs shown in the BOM do reflect purchasing full packs.
- Some fasteners are omitted from the CAD file. Their locations are indicated in this document.

## Documentation Sources & License

We have made the original CAD files (SOLIDWORKS 2016), STL files for 3D printing, and PDF drawings of all mechanical parts available on Github: https://github.com/FlySorterLLC/MAPLEHardware

If you do not have a license for SOLIDWORKS, you can still open the files using the free eDrawings Viewer (http://www.edrawingsviewer.com/).

The circuit boards in this project were designed using KiCAD EDA open source software (http://kicad-pcb.org/). The original design files are also in the Github repository.

All files are distributed under the GNU GPL v2 (see LICENSE file).

## Pre-Assembly

### Aluminum Extrusions

Tap the holes in one end of each of the four 410 mm long 20x40 extrusions with M5 threads:

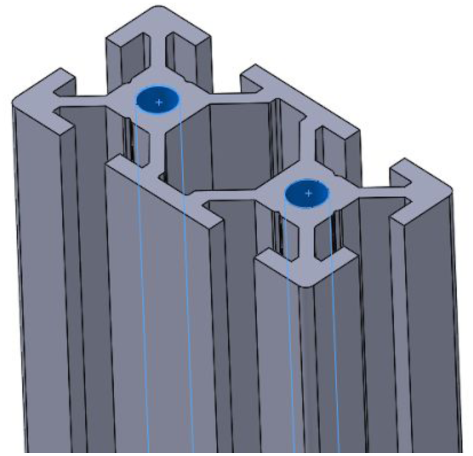

## 3D Printed Parts

Settings
- Maximum 0.2 or (preferred) 0.15 layer thickness
- Minimum 0.6 mm top and bottom thickness
- Minimum 0.8 mm shell thickness
- Minimum 25% infill

Print the following (STL files available in Github):

| Part | Quantity |
| --- | --- |
| BeltClamp-AD-FU | 1 |
| BeltClamp-AU-FD | 1 |
| BeltAdjuster | 2 |
| DragChain-YMount | 1 |
| LimitSwitch-Holder-XAxis | 1 |
| Z2-Carrier-Switch | 1 |
| IS-Camera-Holder | 1 |
| Bellows-cavity-half | 2 |
| Bellows-core | 1 |
| 2mm & 4mm spacer | As needed |

Use a 3.2 mm drill to open these holes to the proper diameter:

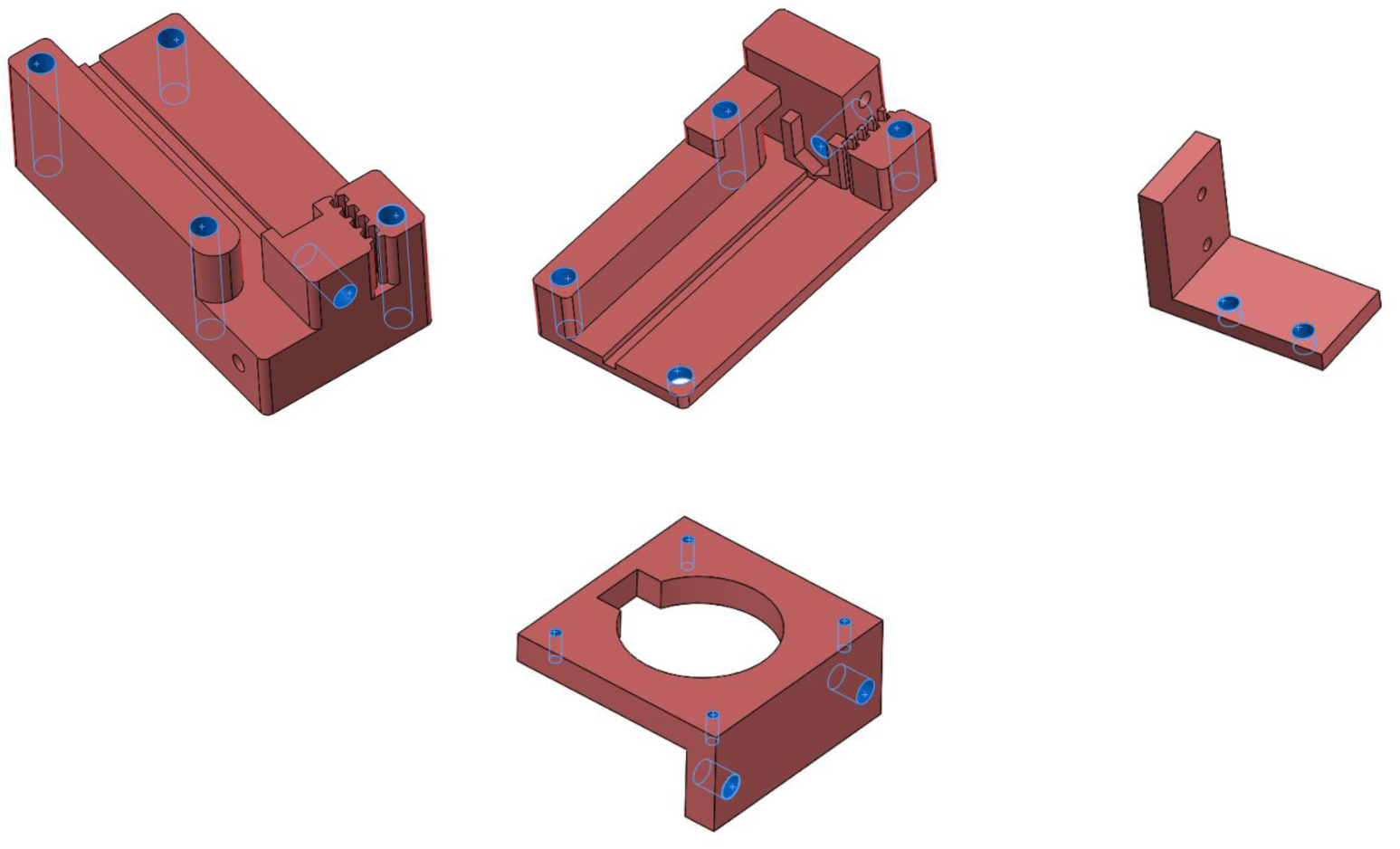

5.5mm drill:

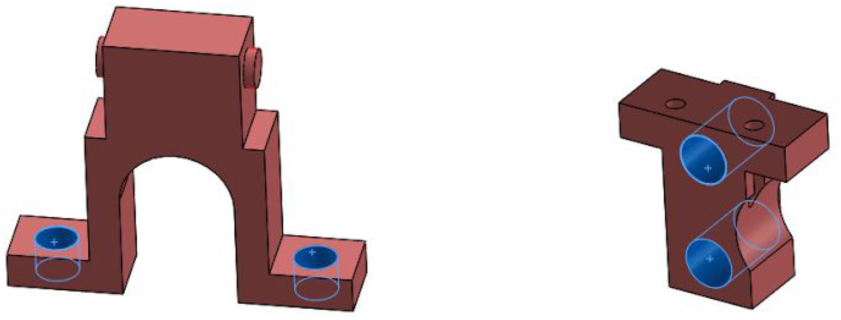

### Silicone Rubber Boot

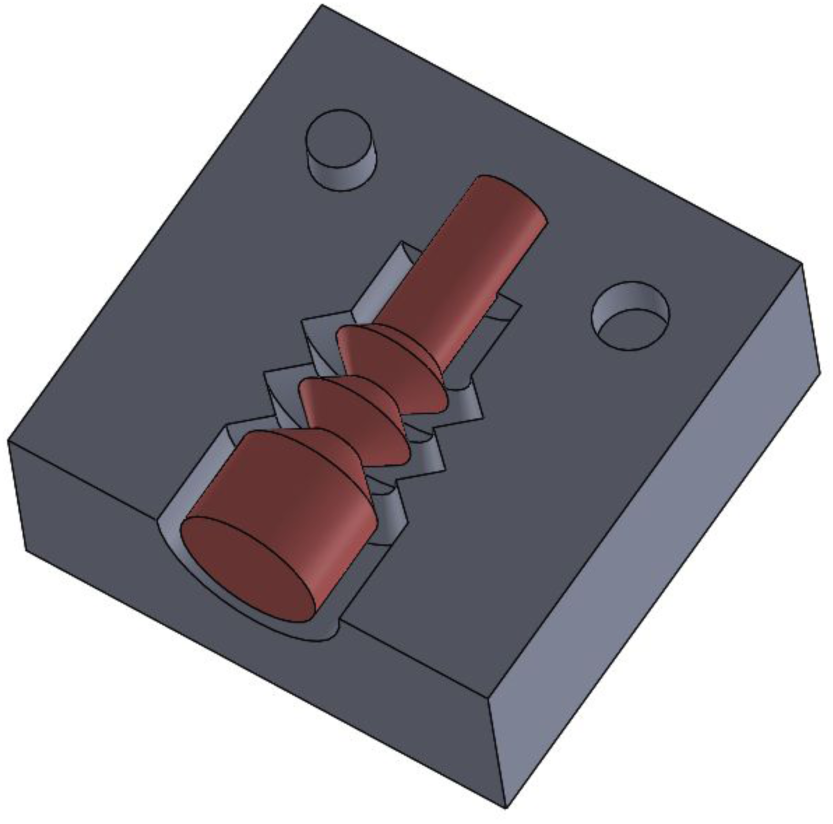

Coat the working faces of the three mold parts (two cavity halves and one core) in mold release (or vaseline, in a pinch). Clamp the two mold halves together, and fill the resulting cavity with liquid silicone (or some other elastomer), insert the core, and allow to cure. Trim off any excess, and remove from the mold.

### Machined Aluminum Parts

See PDF drawings for the small and large part manipulators in <u>Github</u>. It is also possible to 3D print these parts instead.

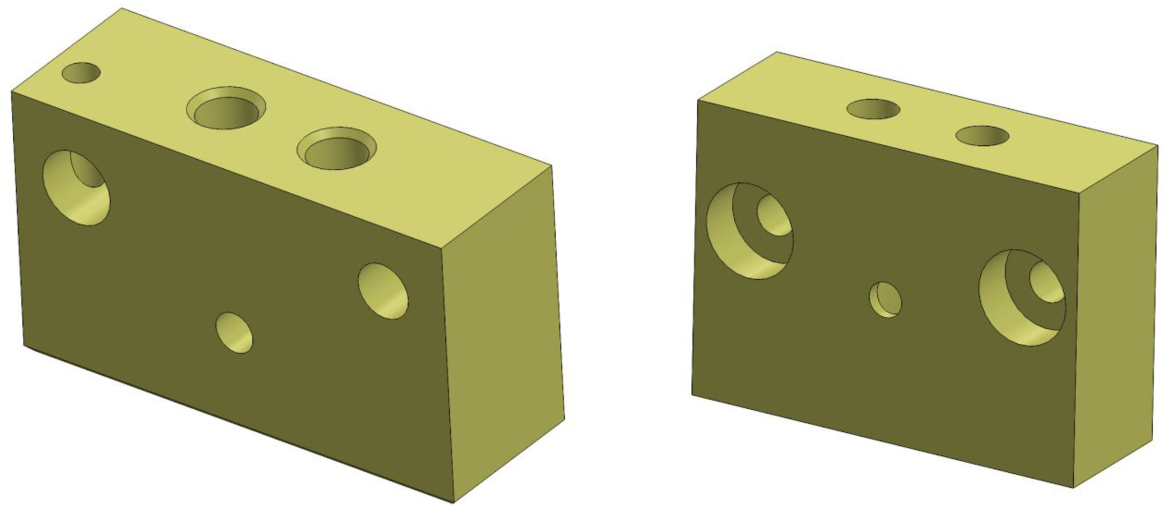

## Waterjet Aluminum Parts

See the DXF in the Drawings directory of the Github repository.

| Material | Part | Quantity |
| --- | --- | --- |
| 1/8" Aluminum plate | Upper-Y-Plate | 1 |
|  | DragChain-Plate | 1 |
|  | Z-Mount | 3 |
|  | Z-Plate | 1 |
| 3/16" Al plate | Lower-Y-Plate | 1 |
| 1/4" Al plate | Stepper-Plate | 2 |
|  | Idler-Plate | 2 |

If your waterjet parts are spot on, you should be able to tap holes without pre-drilling. Measure a couple and compare against the PDF/DXF to be sure.

### Secondary operations

Tap M5 threads (drill to 4.2 mm if holes are small):

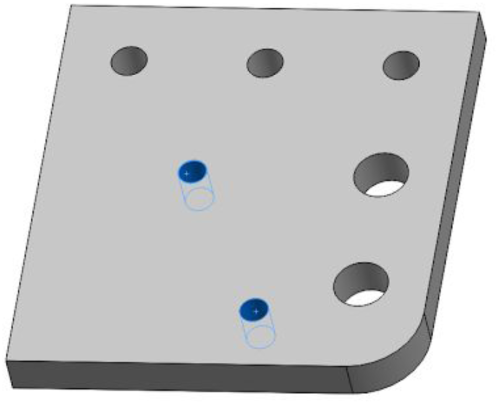

Tap M3 threads (drill to 2.5 mm if necessary):

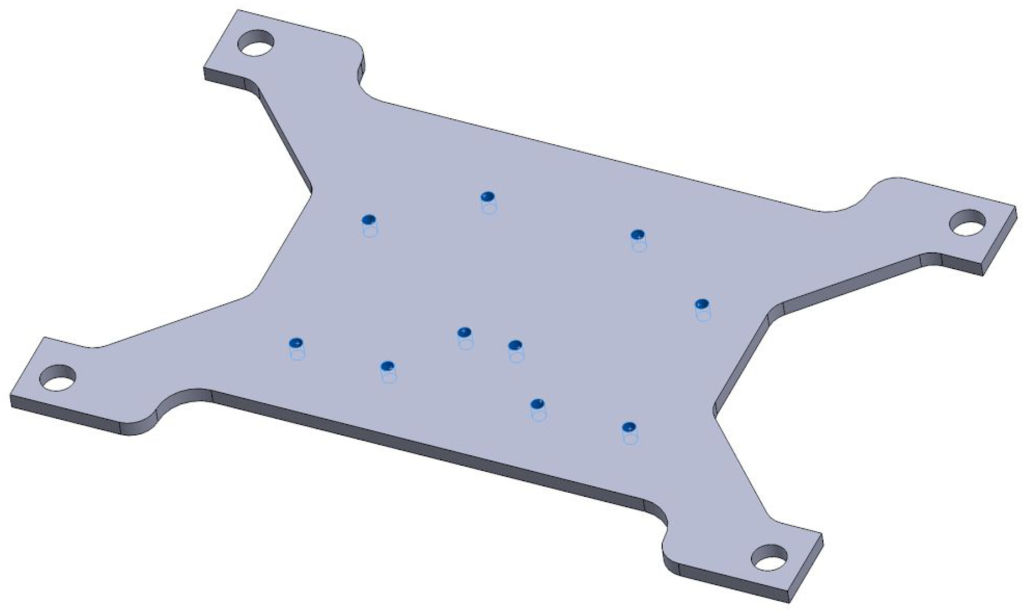

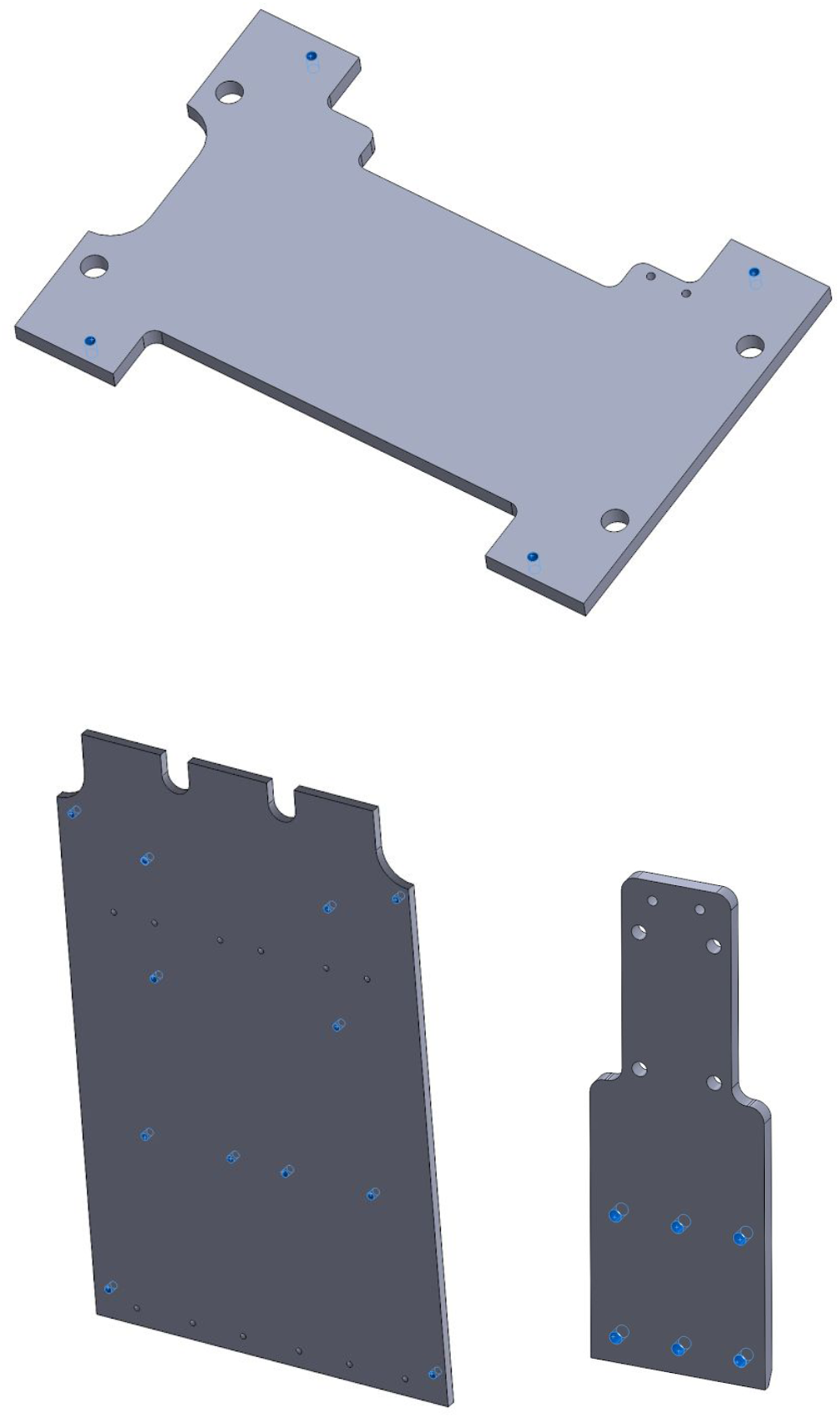

Tap M2.5 threads (drill to 2 mm if necessary):

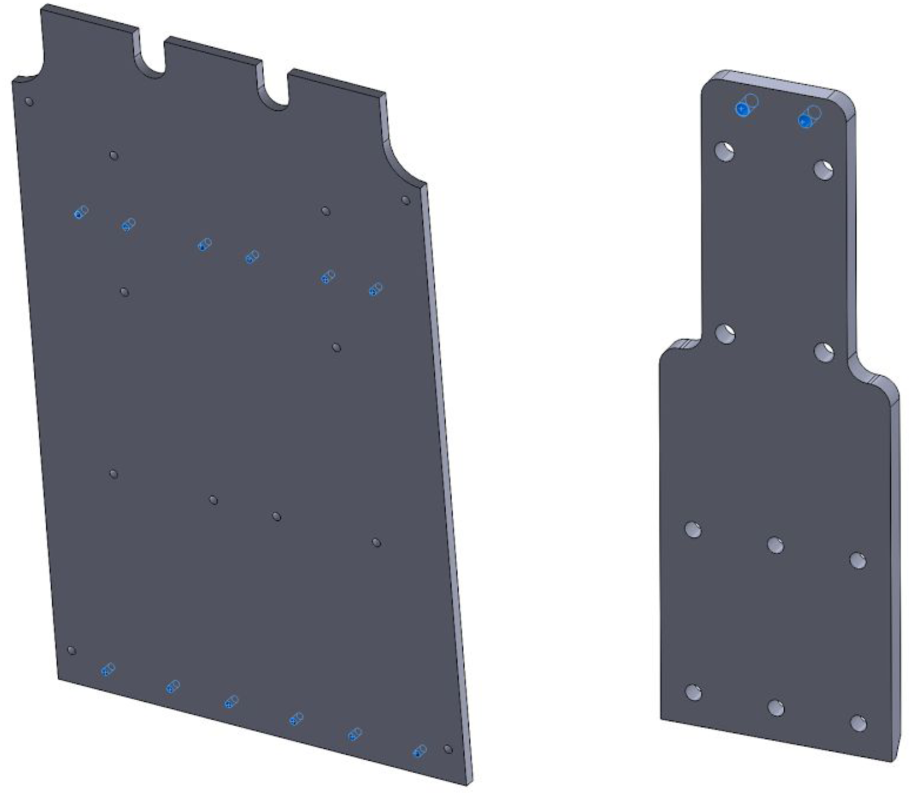

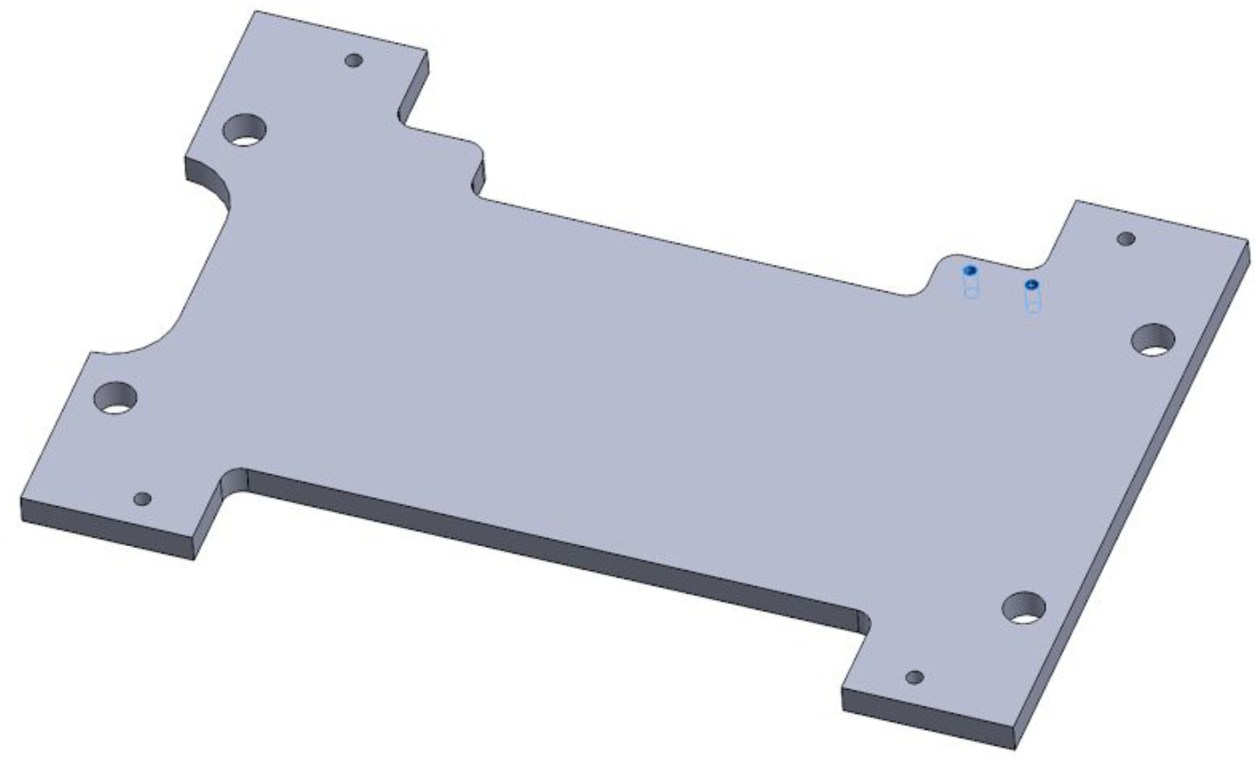

File / grind ~30° bevel:

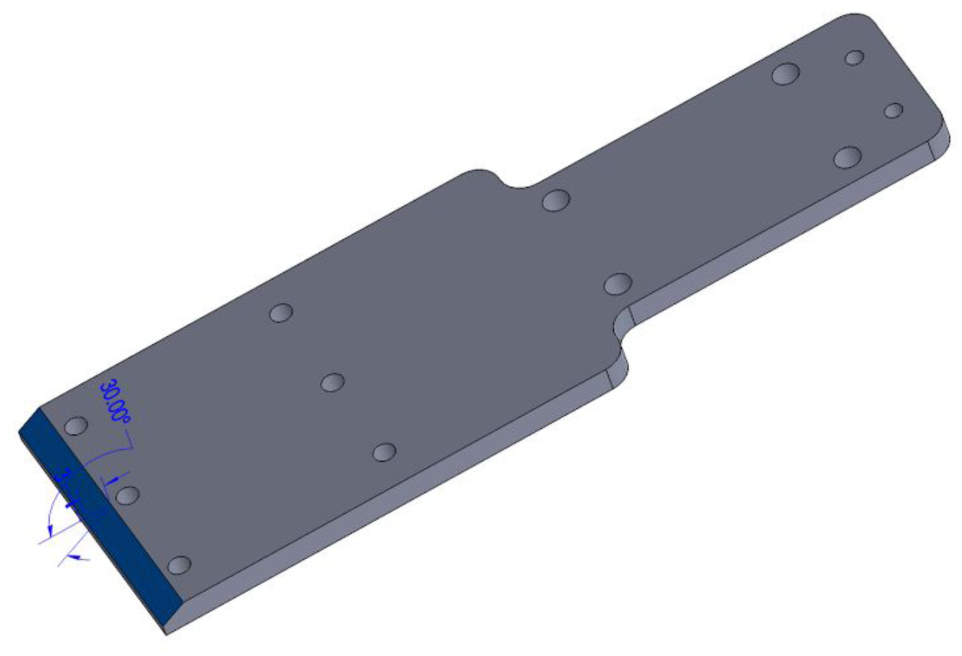

### Laser Cut Parts

Cut the spacer stick from %” or %” acrylic:

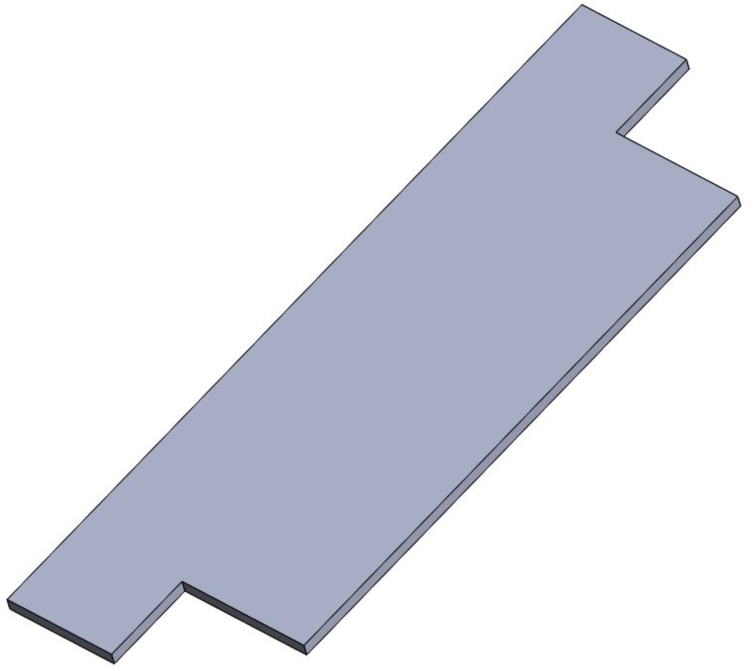

## X Axis Rails

After cleaning & degreasing the parts, glue 1116 mm long IGUS rails to the 1120 mm long 40mm square aluminum extrusions using epoxy. This flexible epoxy from McMaster is recommended (part number <u>75045A65</u>), as it bonds well to anodized aluminum. Other glues may perform well, too. Use masking (or other non-gummy) tape to clamp the rails during glue up.

The width of the base of the supported shaft matches the 40mm Misumi extrusions. Make sure they are lined up during assembly (you can clamp flat parts on either side to ensure alignment).

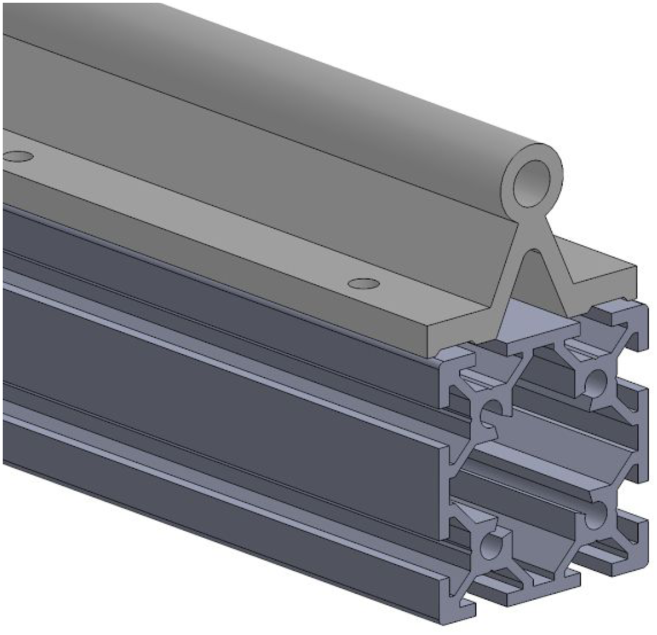

Allow glue to cure fully before assembling the frame.

## X Axis Bearings

Drill out & tap (M5) per drawing.

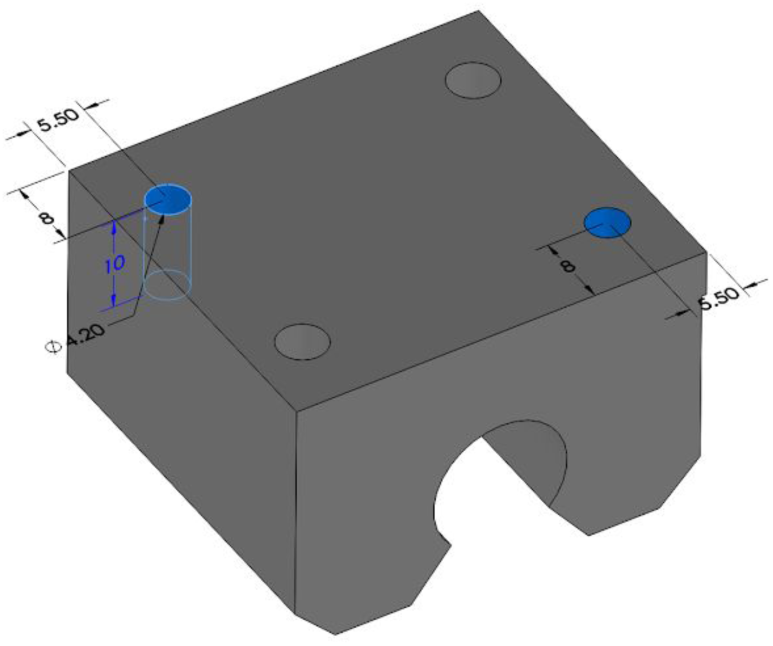

For two of these bearings, drill and tap a 10 mm deep hole on the side with the lip. Make a left and a right version by measuring from different ends for the two blocks.

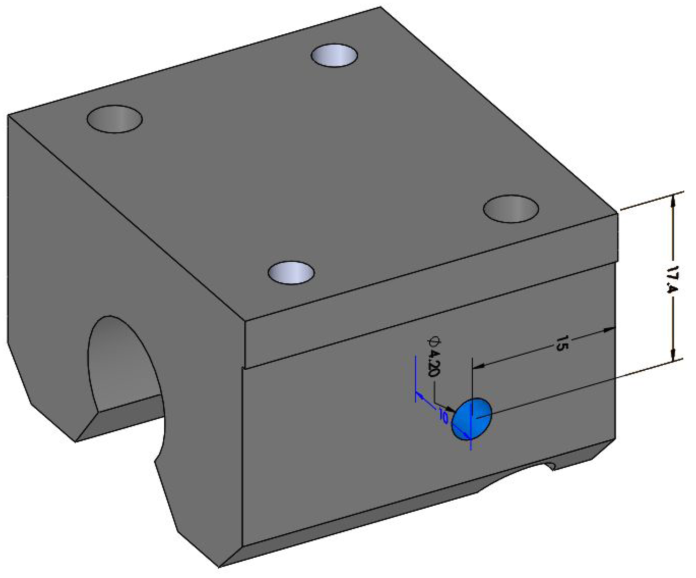

## Y Axis

Drill / mill holes in WS-10–120 part as shown on the PDF drawing:

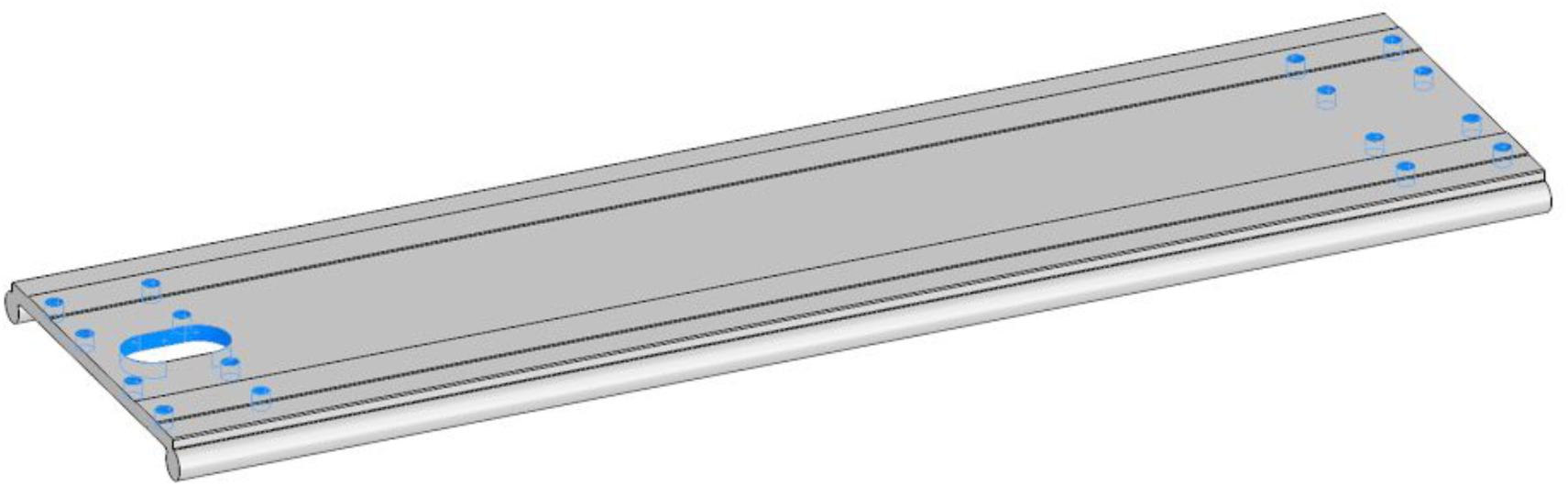

The slot (on the left-hand side of the part in the image above) is for wires and tubes to pass through. Drilling holes and sawing the waste in between them is perfectly acceptable here (if not the most aesthetically appealing), as is using a milling machine, and regardless of how you form that feature, be sure to deburr the edges.

## PCB Assembly

### Smoothieboard

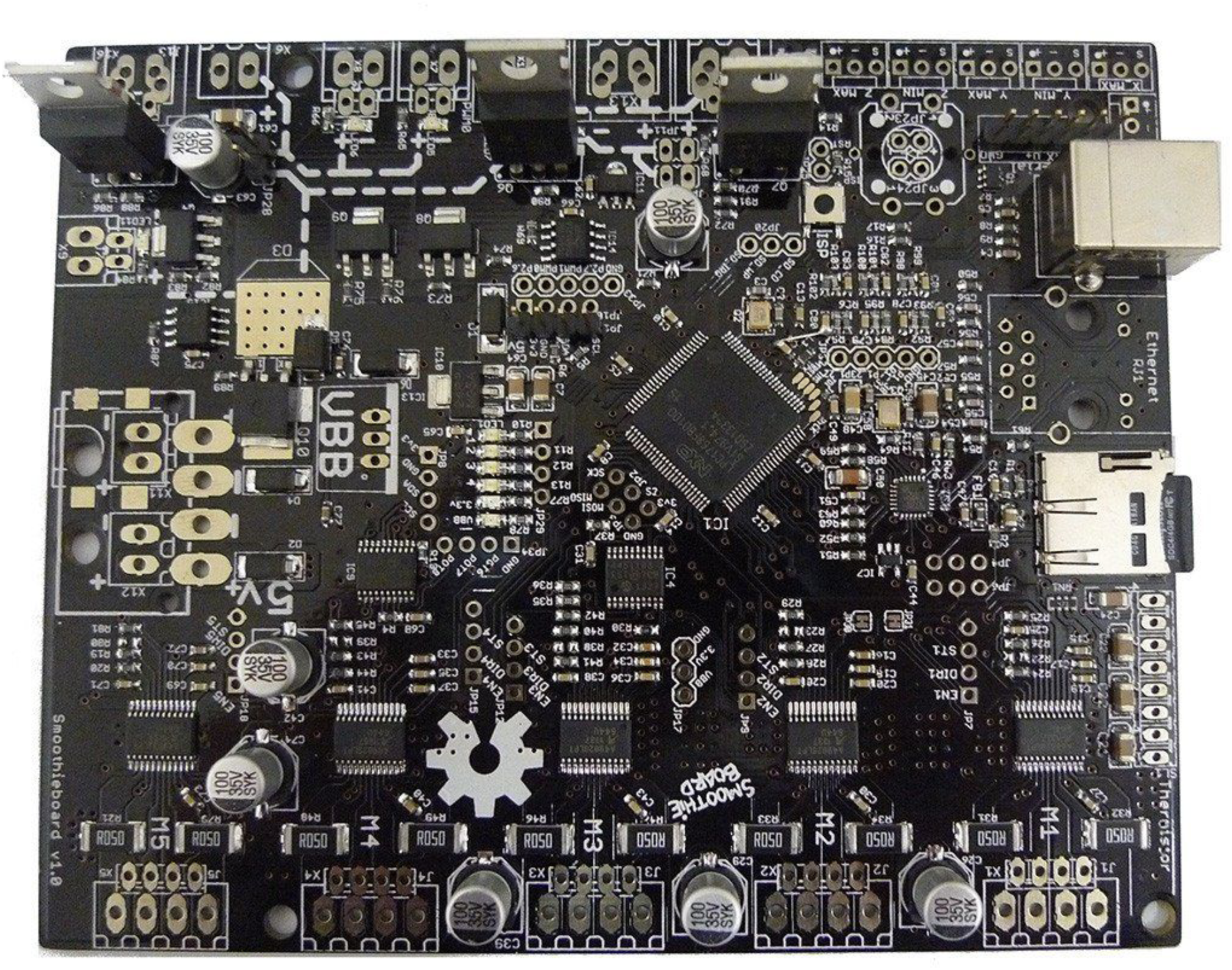

The 5X-xM version of the Smoothieboard does not have any of the connectors soldered in place. We take advantage of this and use header pins to connect an intermediate board (the Smoothieboard-Ribbon PCB). However, some of the connectors are used for MAPLE.

Install the connectors for M1, M2, X MIN and VBB (power in, the screw terminal receptacles). Also install the 5V switching regulator.

**Note:** do not connect or disconnect stepper motors while the Smoothie is powered on.

### Smoothieboard-Ribbon PCB

Solder the ribbon connector receptacle to larger board, as well as the four Schottky diodes (the white stripe of the diodes should be on the left with the board in the orientation shown). Solder the 1 k (R1) and 2k (R2) resistors in place.

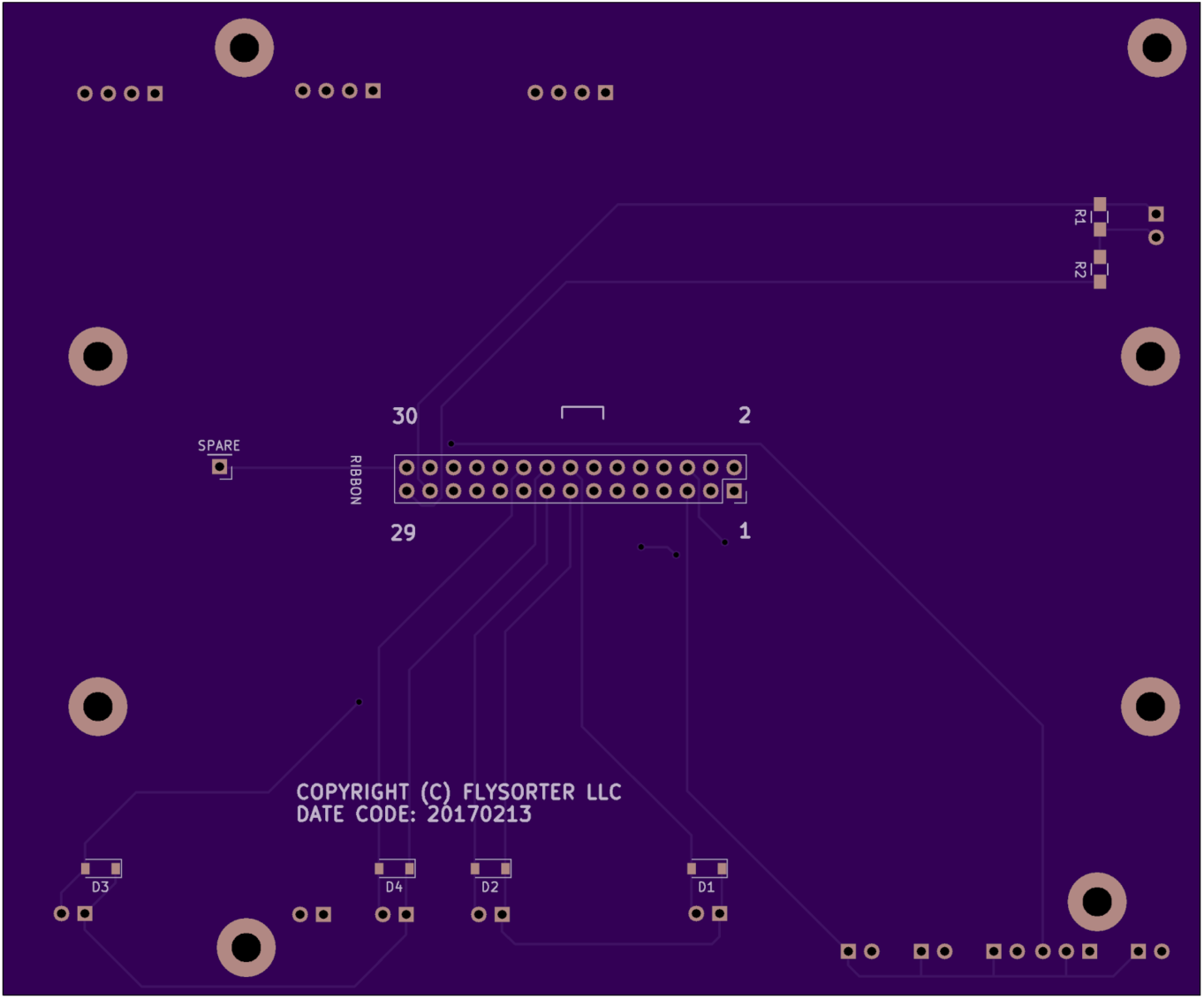

Then, insert the tall header pins between this board and the Smoothieboard and solder in place (the top side of the Smoothieboard should face out from the circuit board sandwich.

### Ribbon Breakout PCB

This PCB is mounted to the back side of the Z-plate. Before installing the board, solder another ribbon connector receptacle to this board, matching the orientation (pin 1) on the Smoothieboard-Ribbon PCB.

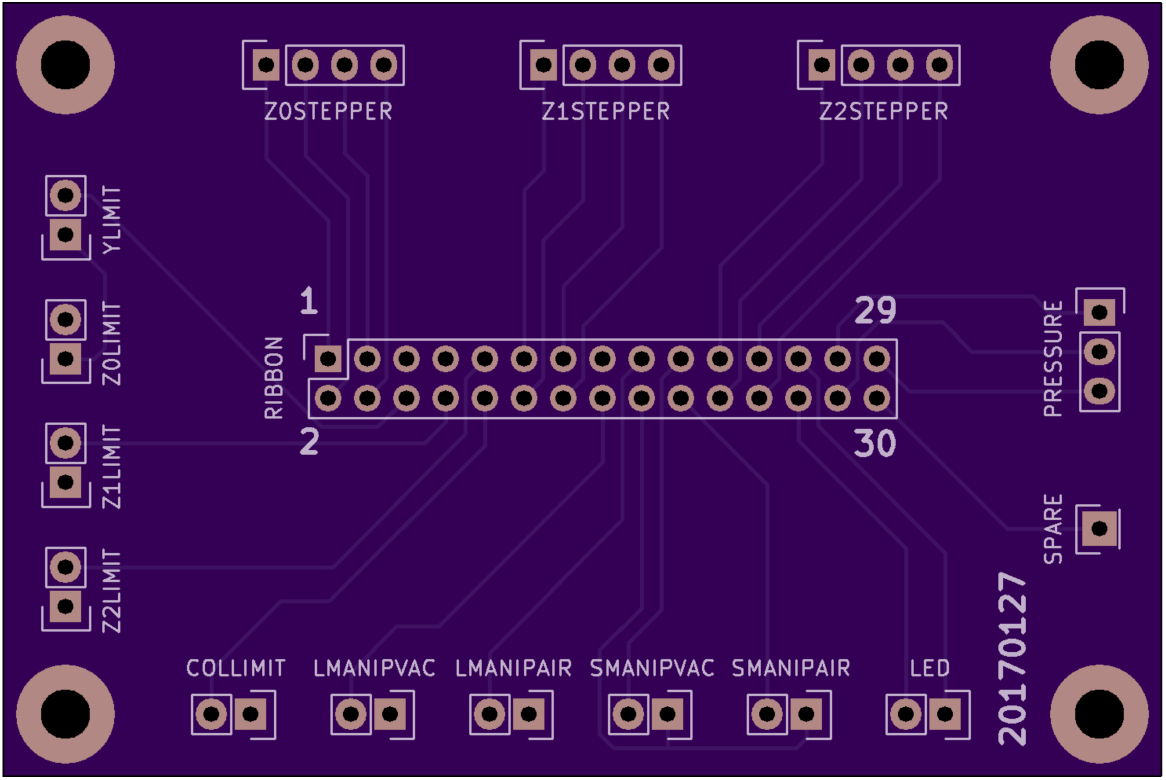

## Frame Assembly

First, assemble two identical rectangles that will form the sides of the frame with two of the 410 mm long 20×40 extrusions (the vertical members), one 1120 mm long 20×40 extrusion (the lower rail) and one 1120 mm long 40×40 extrusion with the linear rail glued to it, using four of the wide corner braces (HBLFSDK5). Be sure to slide the modified bearing blocks onto the rail before assembly, and be sure they are facing the same direction. Make sure the two 0JUM-06–12 blocks with drilled & tapped M5 threads on the side are on the same rail, with the faces that you measured the 15 mm distance from on the inside. Use M5 × 10 mm socket head cap screws and extrusion nuts.

Use the notch in the laser-cut template to set the height from the top of the vertical rail to the top of the supported linear rail. Similarly, use the long section of the laser cut template to set the distance between the top and bottom horizontal rails.

Double check for square by measuring the diagonals with a ruler, tape measure, or string/wire.

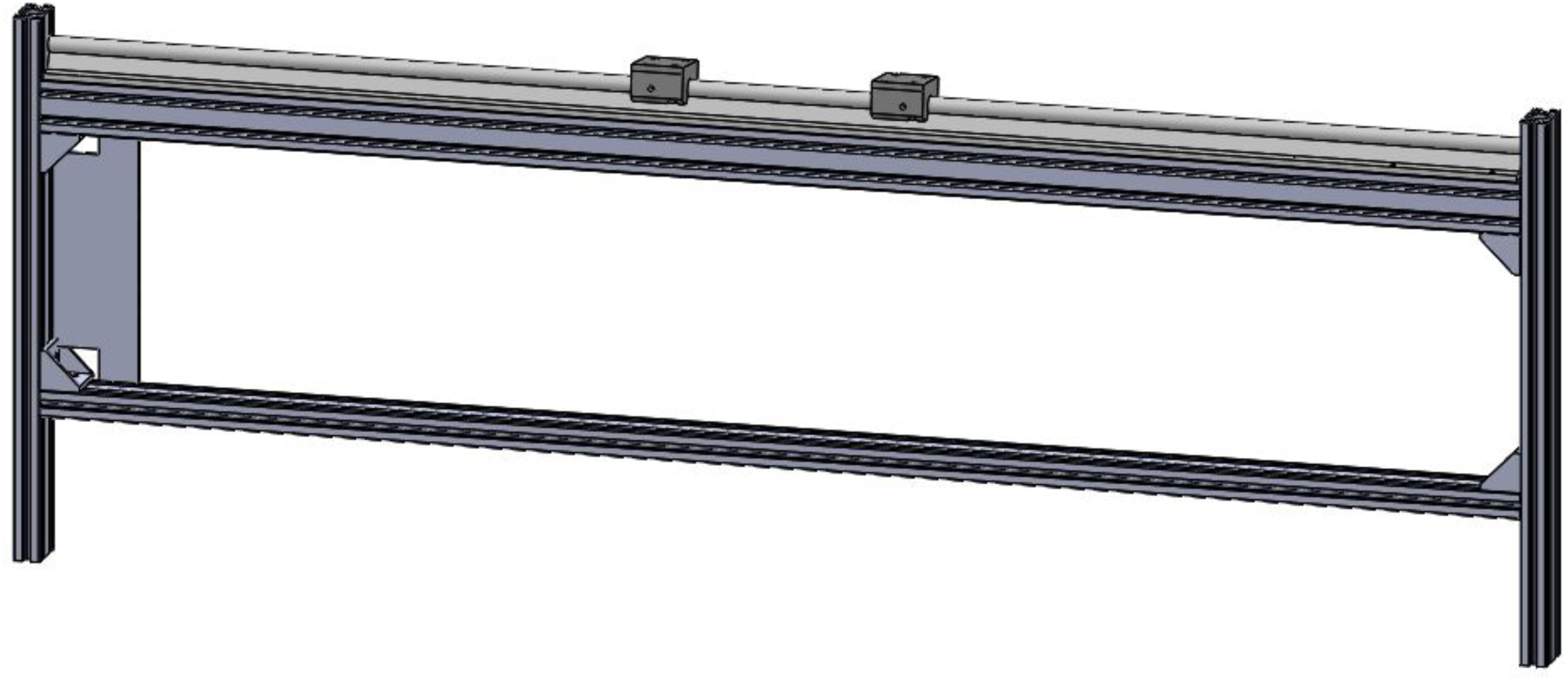

Next, use the triangular sheet metal corners (SHPTCUL5-SET) as well as the narrow corner braces (HBLFSSW5) to join the two sides. Lined up with the bottom rails are two 370 mm long 20×20 extrusions, and at the top are two 370 mm long 20×40 extrusions.Again, check for square using a tape or string to measure across the diagonals (you can also use a large framing square).

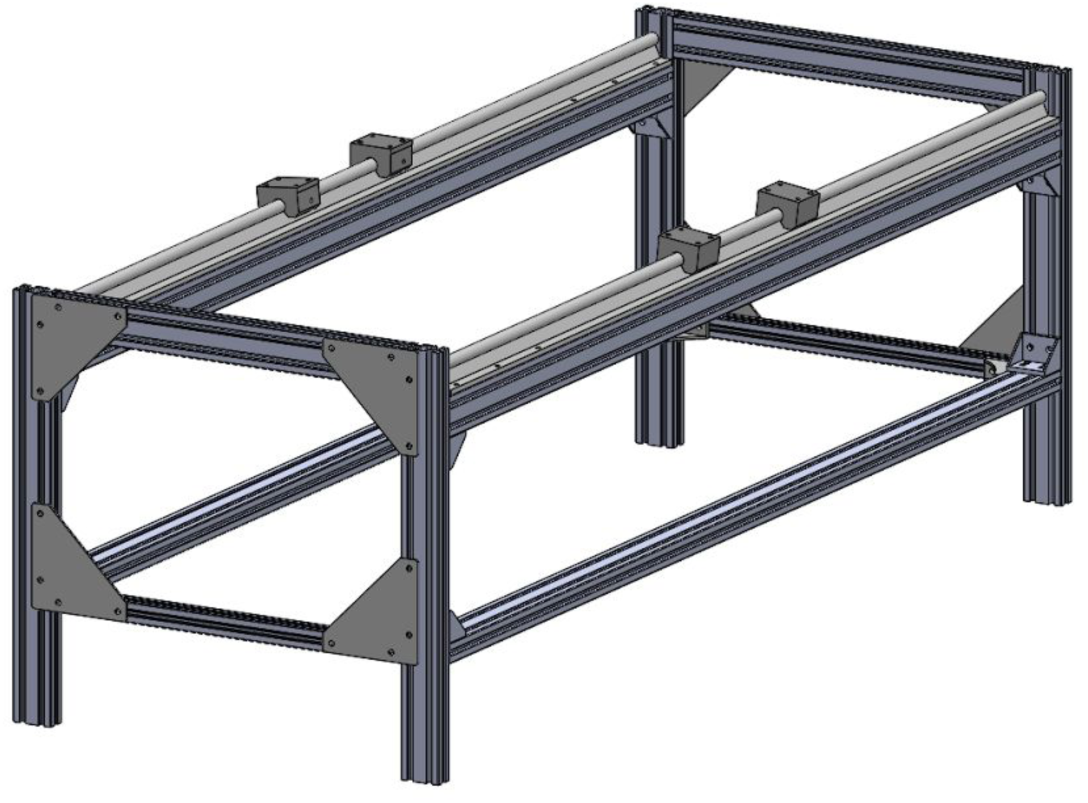

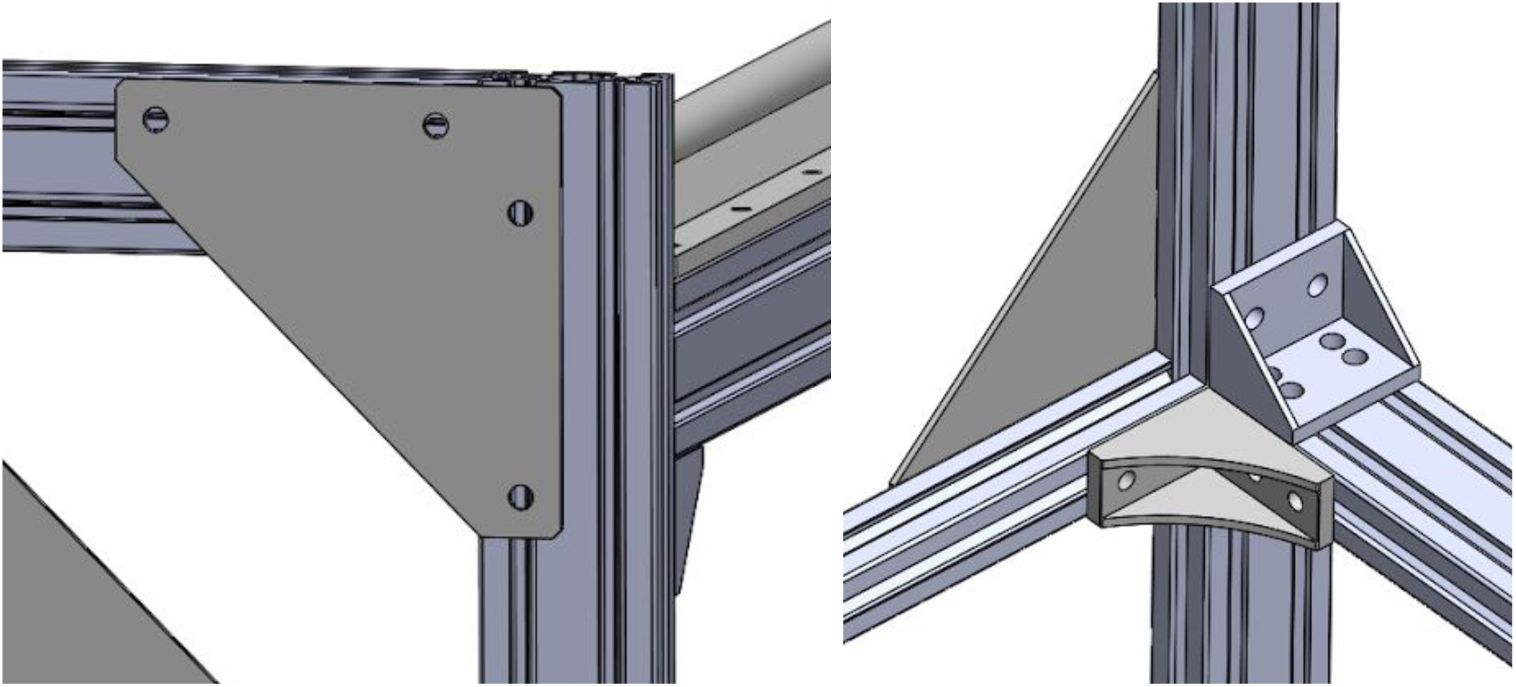

Align the frame with the X axes left-to-right so the rail with the two OJUM-06–12 blocks with holes on the sides is in the back.

Attach the X axis limit switch holder to the rear right upright as shown, using M5 screws and insert nuts:

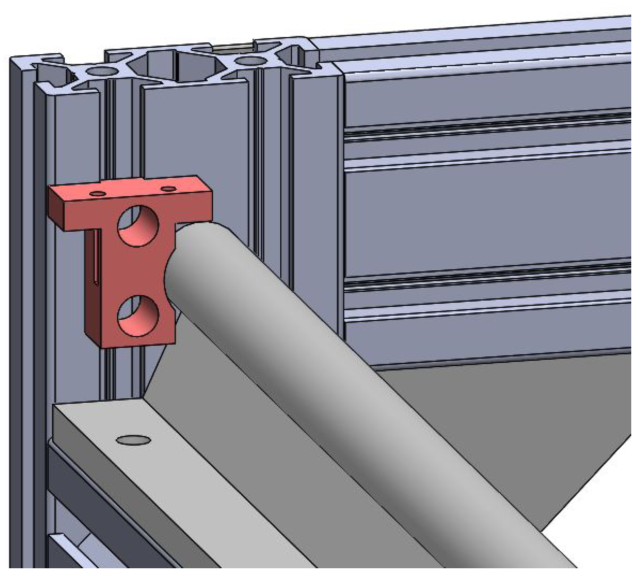

Attach the stepper plates to the top of the 410 mm vertical rails at the right end of the assembly. Attach the idler plates at the other end, with HBLFSSW5 brackets below each one. You will need to add Misumi post-assembly insert nuts on the top horizontal end rails (two per end, one for each plate).

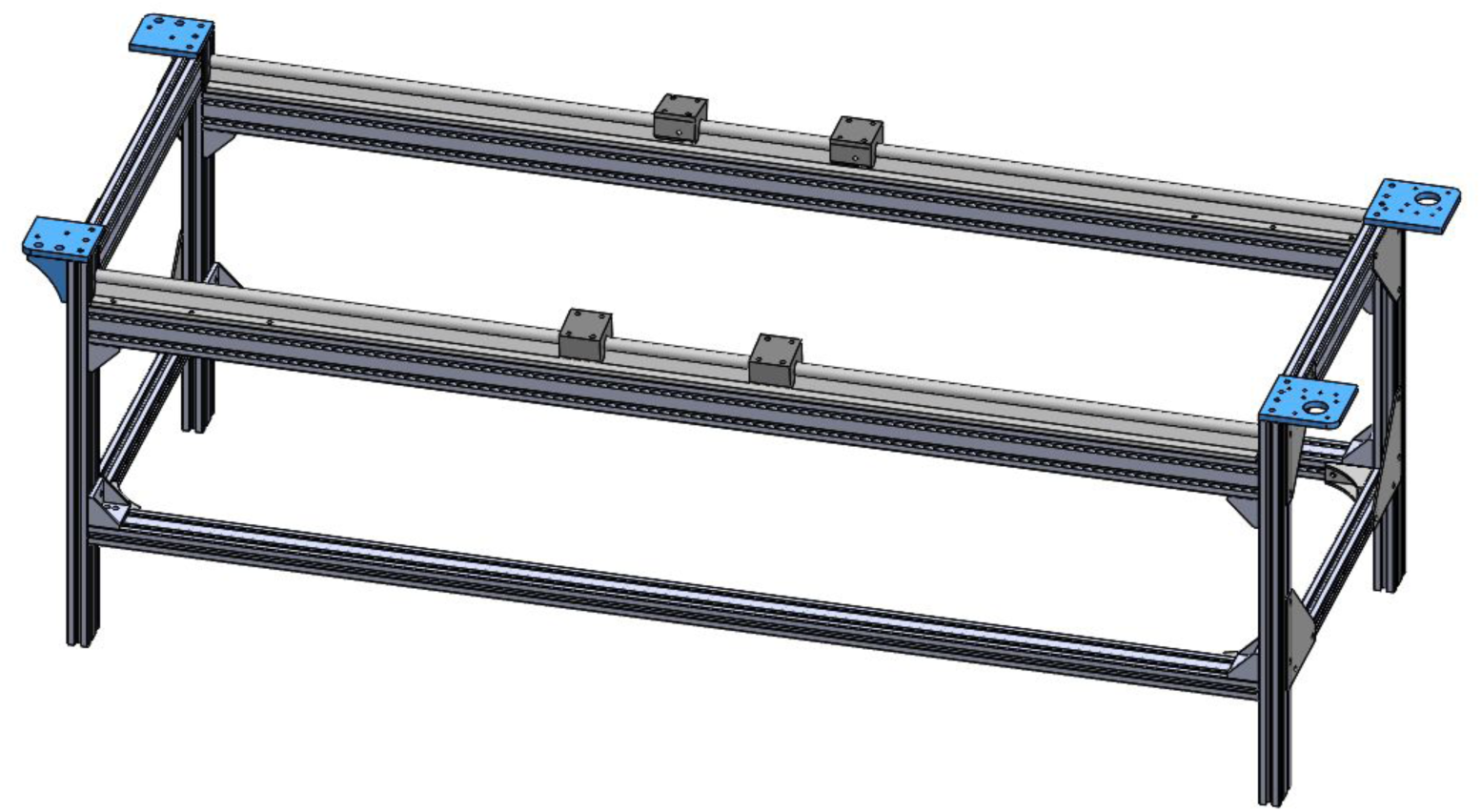

With the stepper motor plates on your right and the idler plates on your left, install the motors (using M3 bolts) and pulleys (using M5 shoulder bolts and 4 mm spacers on top) as shown:

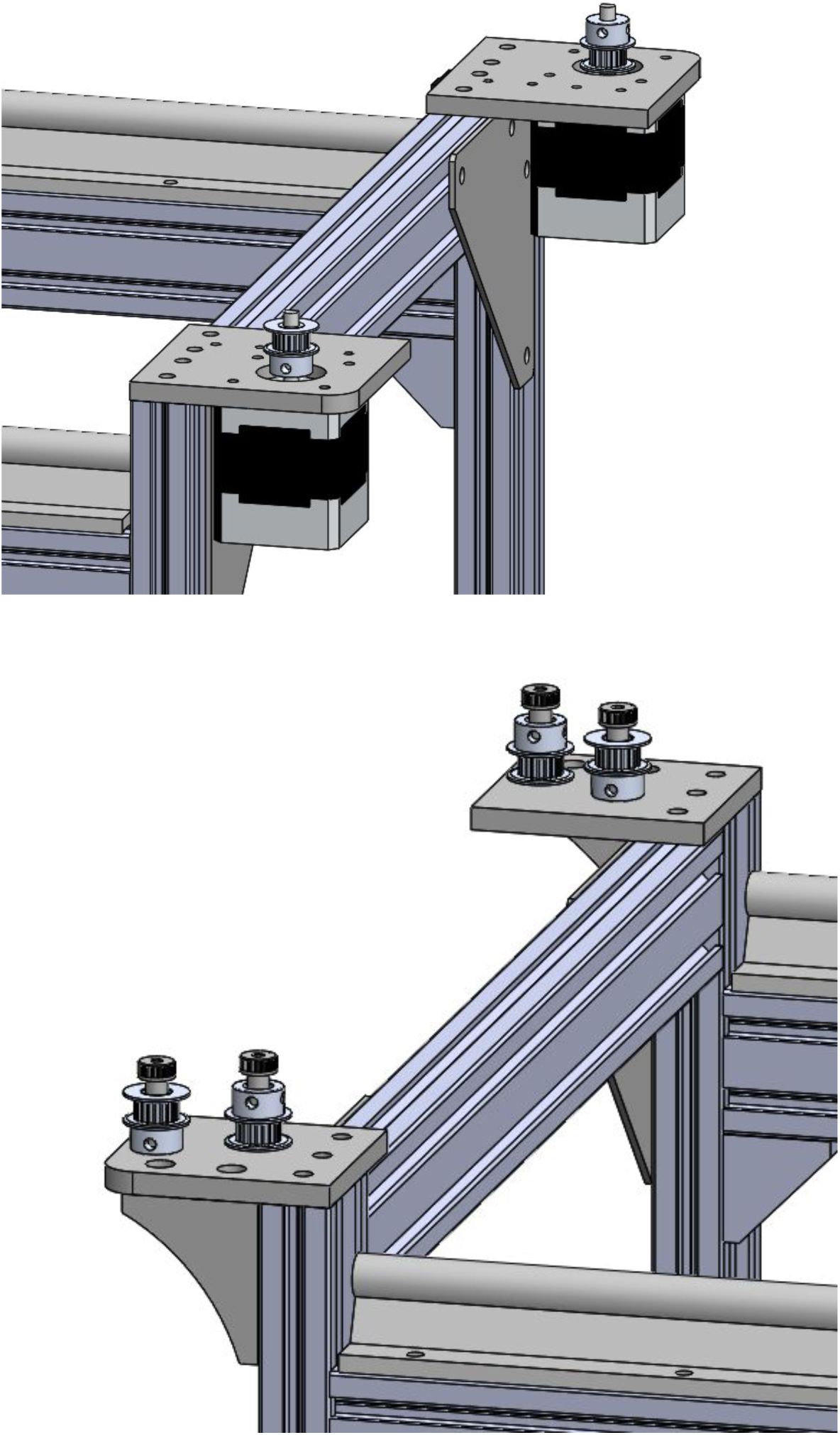

## Y Axis Assembly

Attach the upper and lower Y plates to the WJ200UM-01–10 bearings using M6 screws as shown:

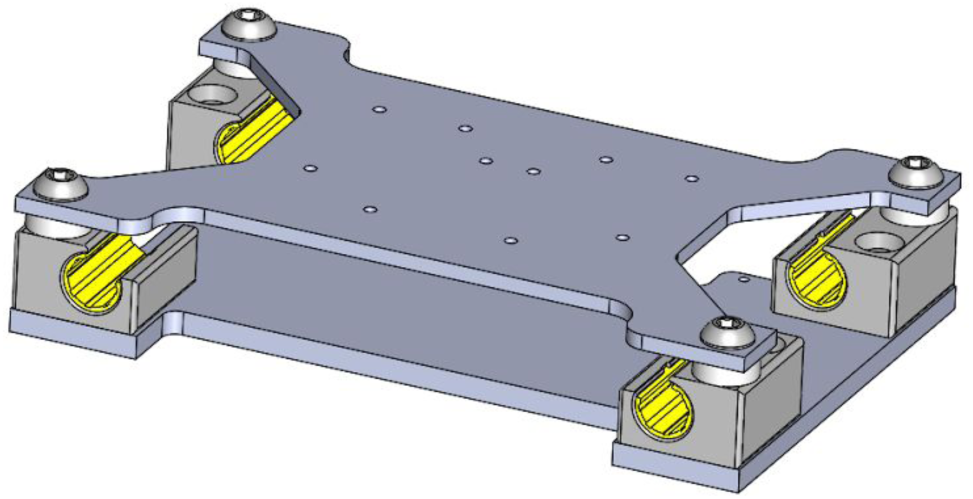

Then attach the two belt clamp parts in the orientation shown below (the two parts are distinct, and must be installed correctly for the belt heights to line up).

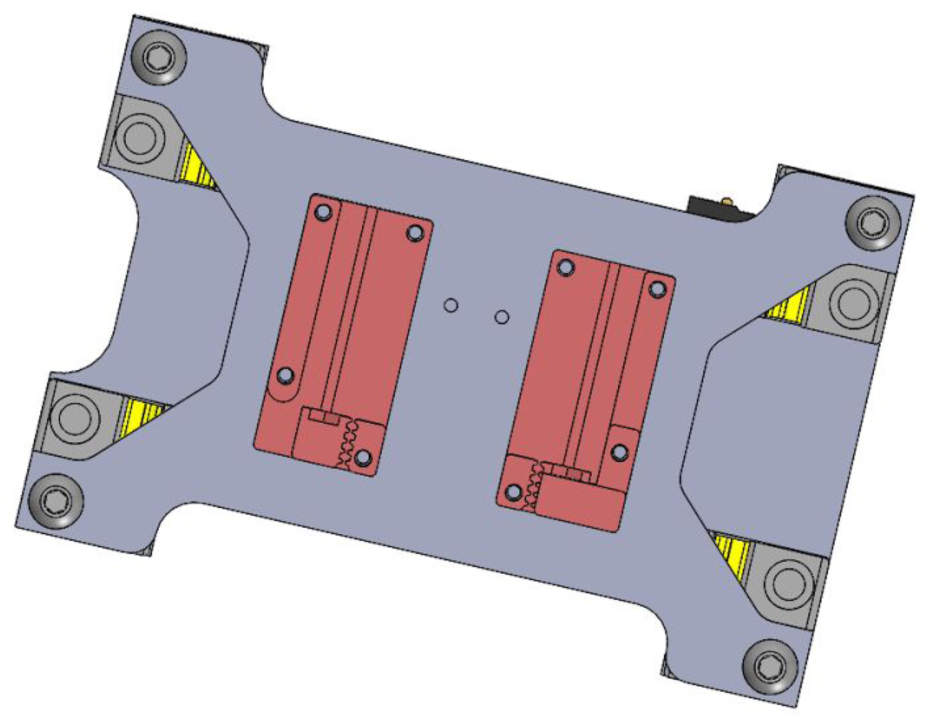

Slide the Y carriage assembly on to the WS-10–120 rail, then install the idler pulleys (above the %” nylon spacers) with M5 shoulder bolts. Note that if the screw threads from the shoulder bolts protrude below the bottom surface of the WS-10–120 rail, you’ll need to cut / grind down the screws so they don’t interfere with the X axis bearing blocks. Again, be careful to get the orientation of the carriage and the pulleys correct.

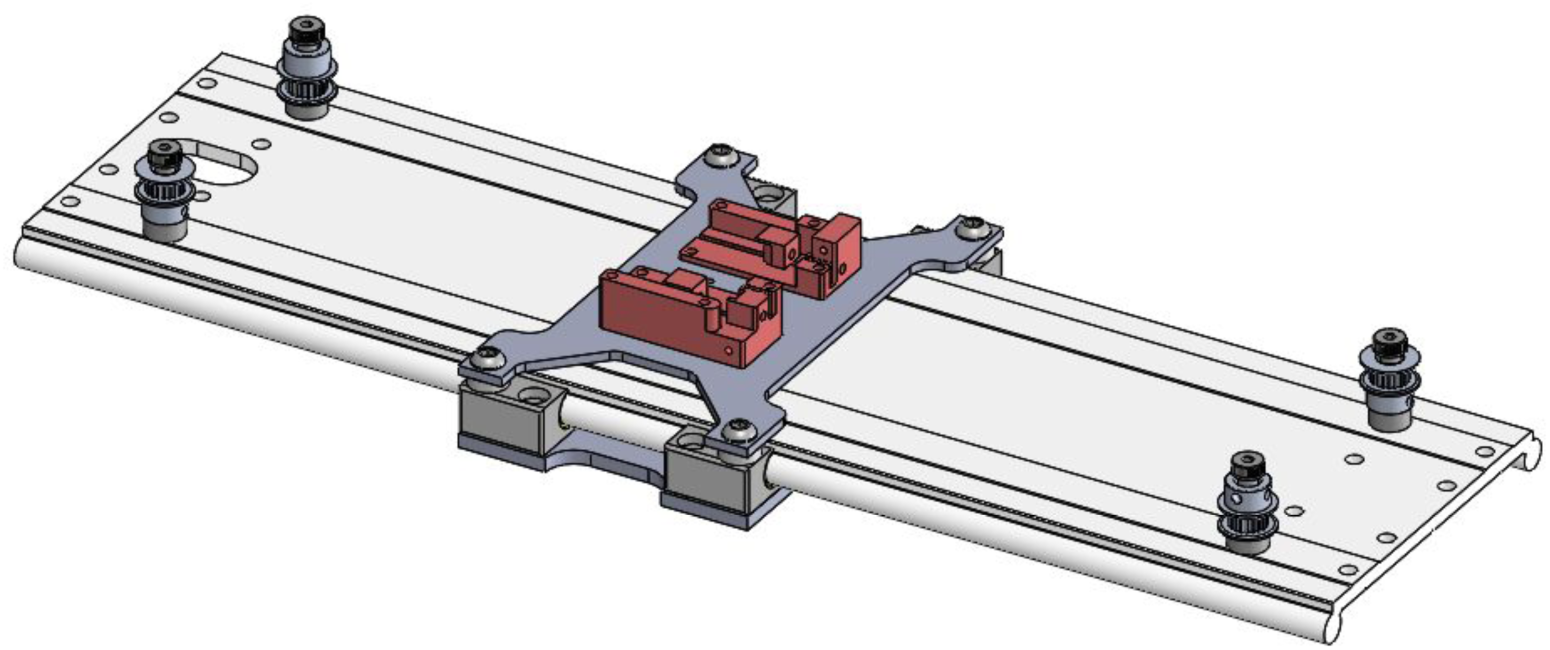

Attach the Y axis assembly to the X axis bearing blocks using M5 screws as shown. Each side requires 6 M5 screws, and the DragChain-YMount part is captured by two of them on the back rail. Do not tighten at this point.

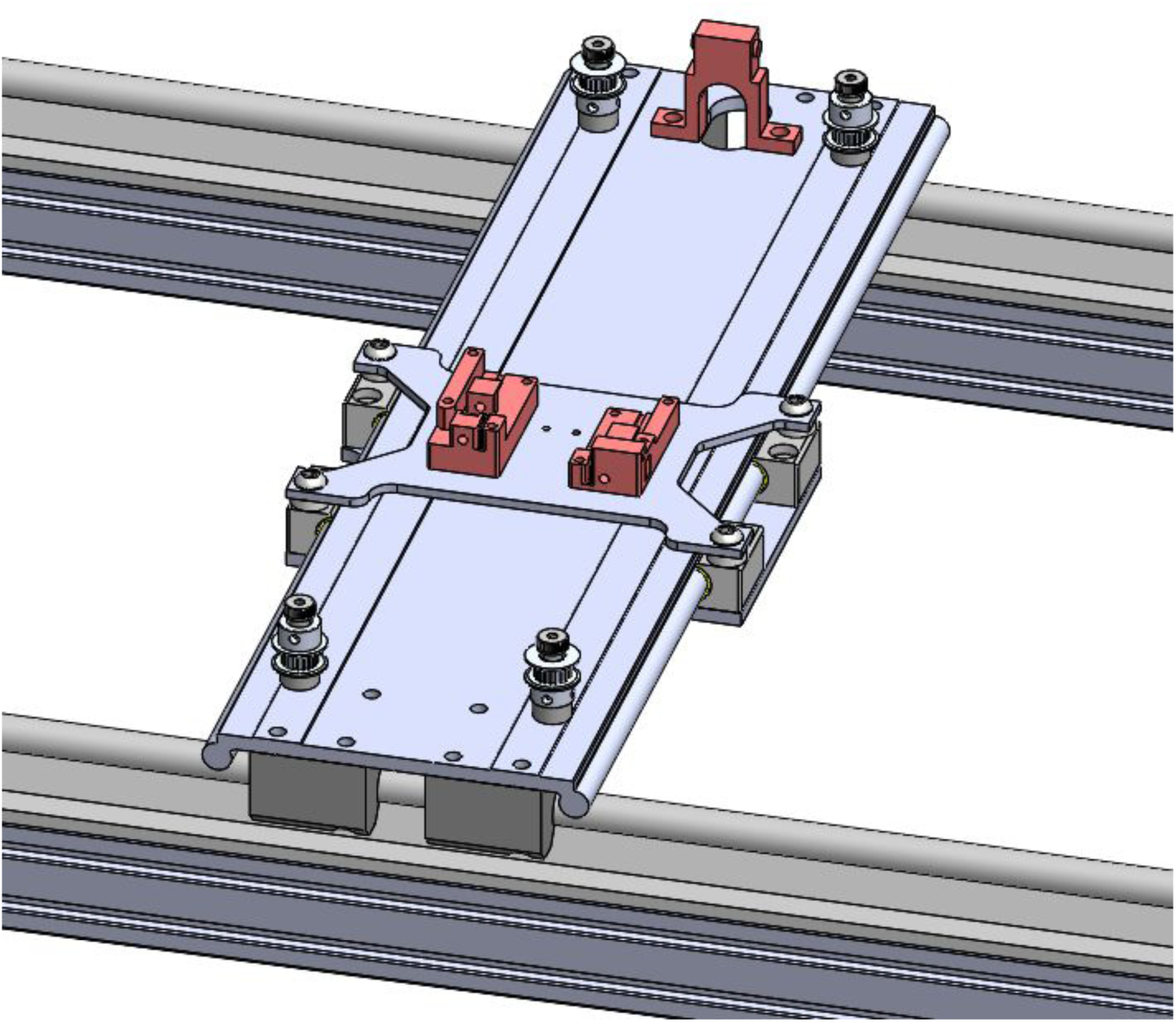

Finish assembling the belt clamps by inserting an M3 nut into each one, and then threading M3x25 screws through. Belt tension will keep the BeltAdjuster parts in place at the end of each screw.

Mount the DragChain-Plate between the two bearing blocks on the back rail:

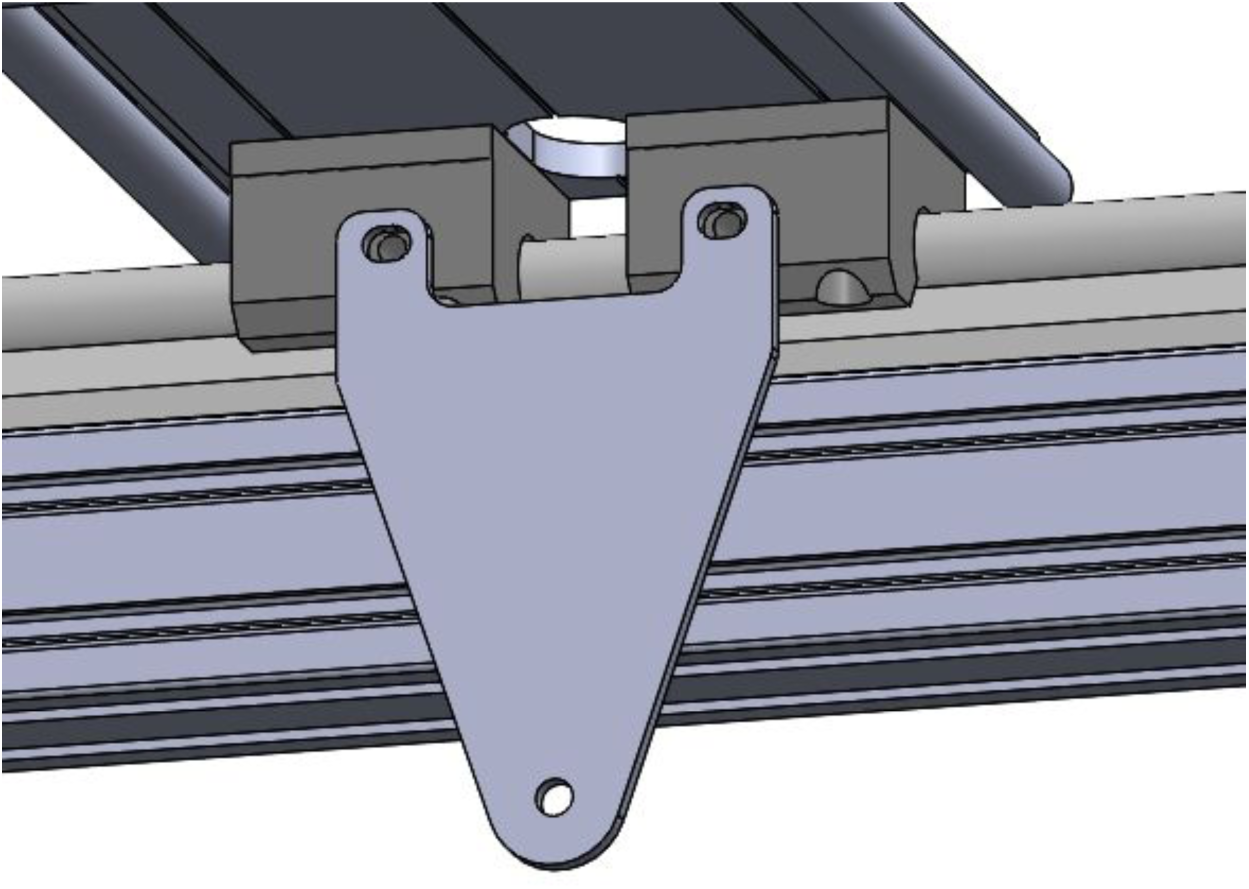

Cut two 3.5 m lengths of GT2 belt. Punch a small hole in one end of each belt, and use an M2.63 plastic screw to attach to the BeltClamps. Thread the belts as shown, snaking them through the belt clamps, and then using the slots in each clamp to hold the end of the belts. Trim excess.

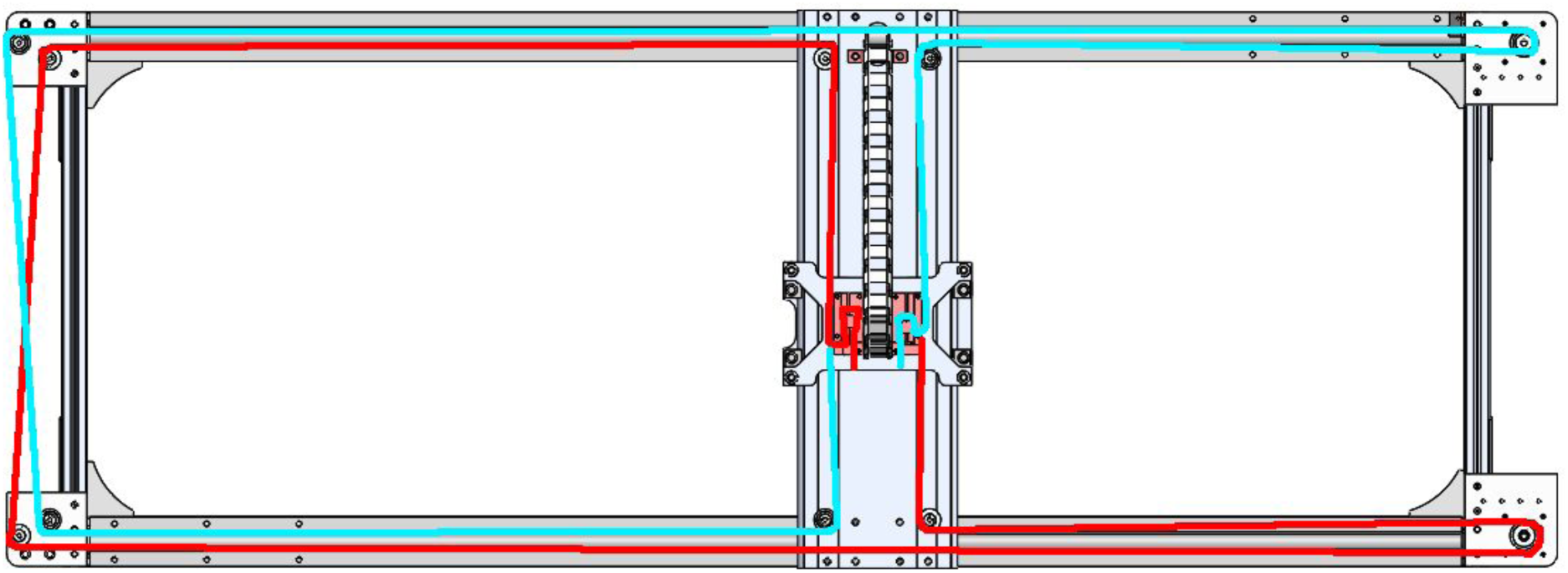

Belt tension and Y axis alignment will be addressed later on.

Attach the smaller drag chain between the Y carriage and DragChain-YMount, and the larger drag chain on the lower back rail, connecting to the newly installed DragChain-Plate with an M5 bolt and a nyloc nut.

## Assembling the Z Axes

Attach the three IGUS SLN-27–14–0050–75–11-G-S-000 linear slides to the Z plate as shown, using 4 M2.5 × 10 mm screws for each slide.

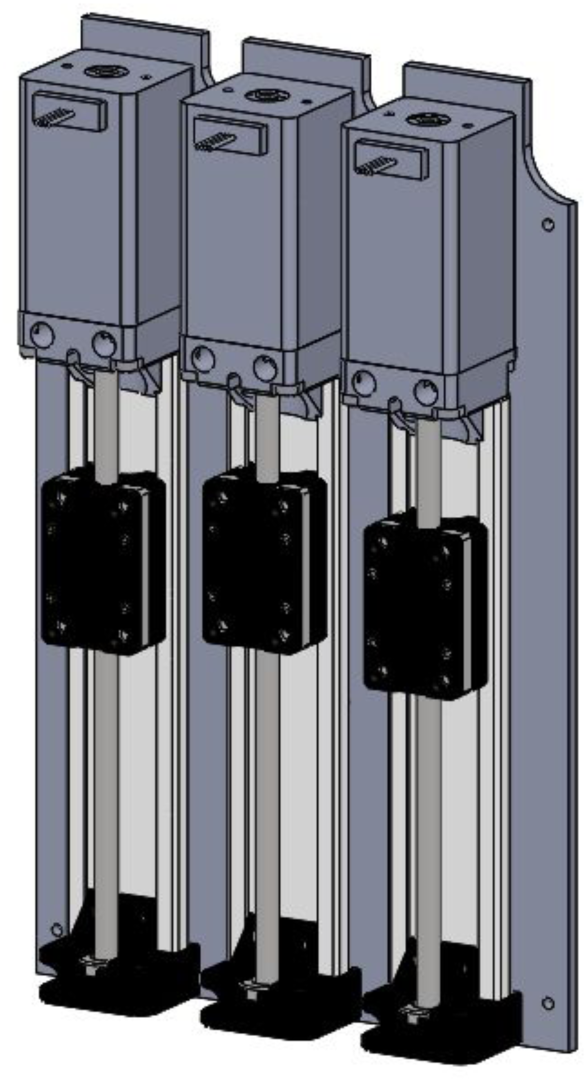

On the reverse side, attach the Clippard manifolds and valves as well as the Ribbon Breakout PCB (using M3 screws and 2 mm spacers). The PCB should be oriented with the stepper connections up.

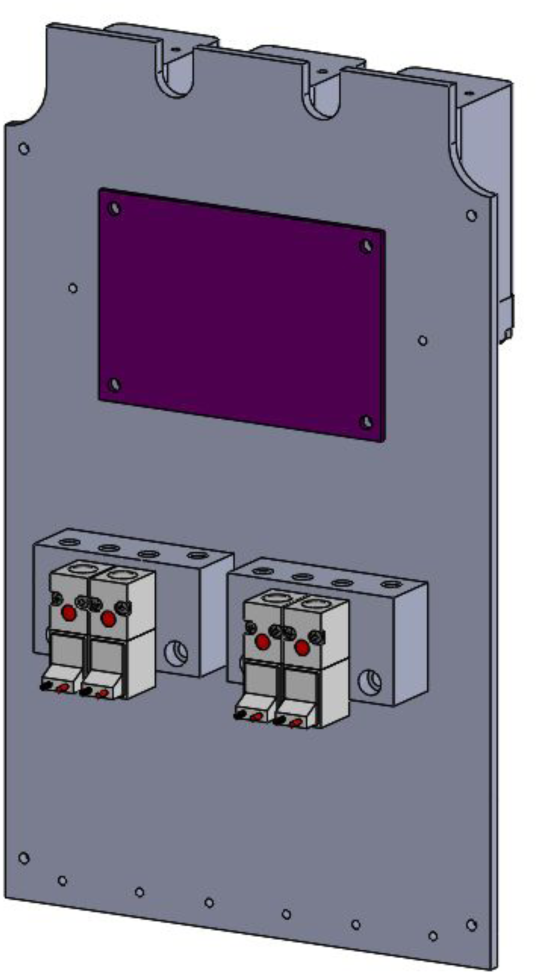

Then screw in 6 4 mm × M5 push-to-connect hose fittings to the Clippard manifolds.

## Z0 - Small Part Manipulator

Attach two 4 mm × M5 push-to-connect fittings to the top of the Large Manipulator Manifold, and the % NPT vacuum cup to the bottom. Attach the manifold block to a Z mount using M3 × 12 mm screws (you may need washers to keep the ends of the screws from protruding through the Z mount).

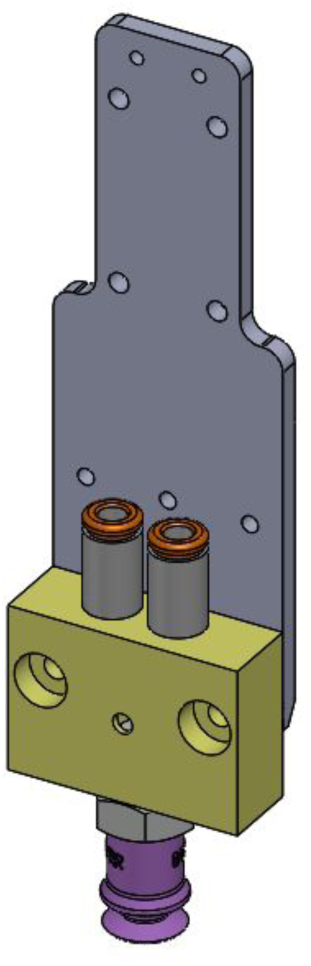

Attach the Z mount to the left-most Z slide using M3 × 10 screws.

## Z1 - Camera

Attach the lens holder to the camera PCB (you may need to remove the original lens holder). Then attach the camera PCB to the IS-Camera-Holder part using the M1.91 screws for plastics and the small, ¼” long nylon spacers to hold the board off the 3d printed part. Make sure the USB plug is in the orientation shown. Screw in the lens.

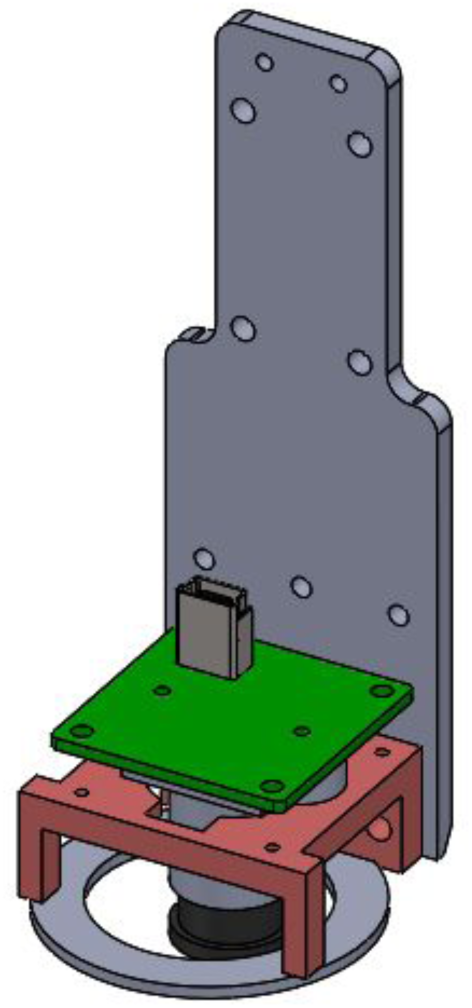

Attach the camera holder to the Z mount, then the assembled Z mount to the middle Z slide.

## Z2 - Fly Manipulator

Using diagonal cutters or a fine jewelers saw, carefully remove two 4 mm sections of the stainless steel tubing portion of the 0.042” OD and 0.028” OD dispensing needles. If needed, use a needle or diamond file to square up one end of each of the four tube sections, and align them so the clean ends face the same way. Insert these tubes as a group into the blunt needle (6710A61) as shown, and push them approximately 4 mm up, forming the fly aspirator:

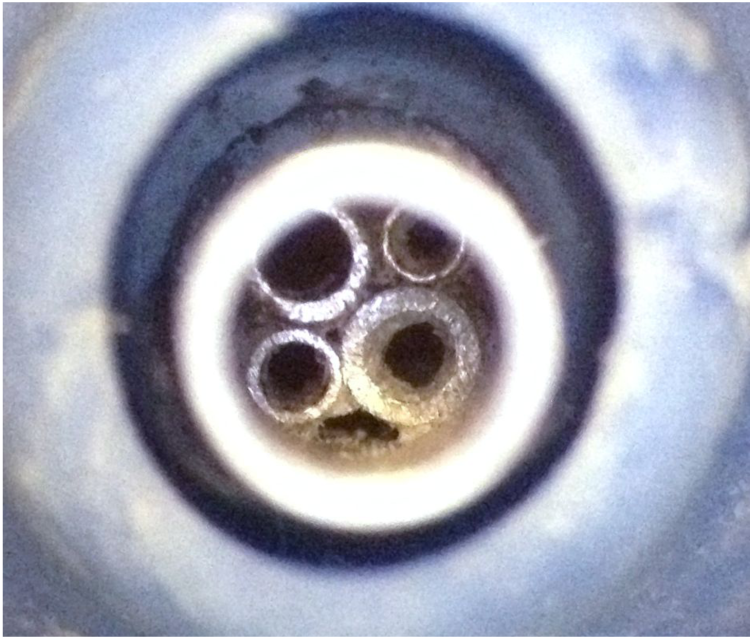

If the small tubes are too loose inside the blunt needle, you can use a little super glue or acrylic dissolved in a solvent (MEK or similar) as a bonding agent. The key is to use a thin liquid so the tubes remain open and air can flow through.

Attach two 4 mm × M5 push-to-connect fittings to the top of the fly manipulator manifold and a plastic Luer-lock quick turn coupling to the bottom.

Using M3 × 12 mm shoulder screws, attach the manifold to the remaining Z mount. The manifold assembly should rotate on the right-hand shoulder screw.

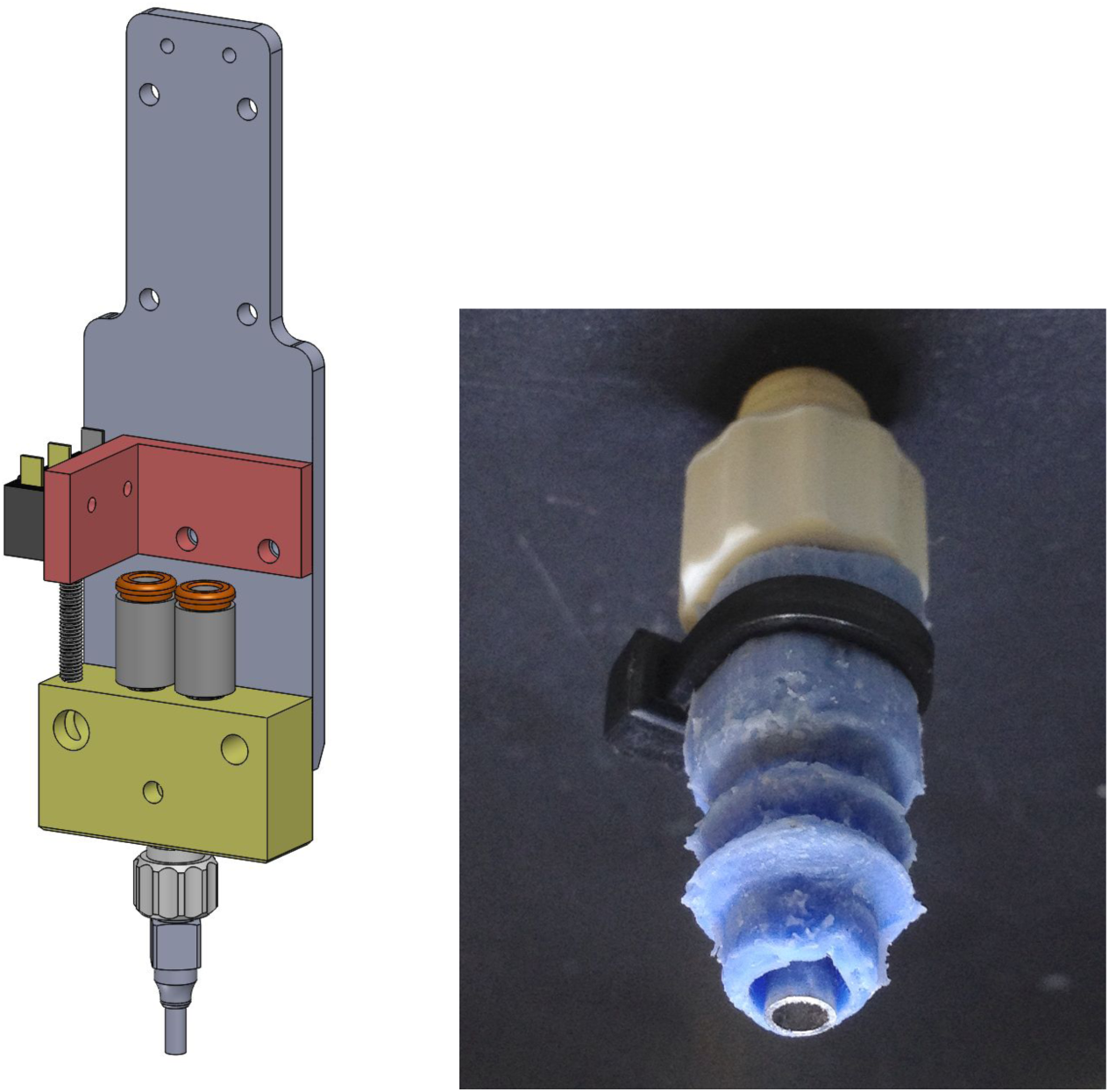

Screw in the 20 mm threaded rod to the top of manifold block, and add an M3 nut to lock in place.

Attach an SS-5 switch to the Z2 switch carrier using M2.5 screws, then attach the switch carrier to the Z mount as shown.

Adjust the M3 threaded rod up or down so that the switch is not triggered when the block is at its lowest point, but the switch triggers as soon as possible when you press up on the bottom of the block. Remove the switch (for now).

Attach the blunt needle (with tubes inserted) to the fitting on the bottom of the manifold block, then attach the silicone bellows with a small zip tie (see above).

Finally, attach the Z2 assembly to the third Z slide.

### Complete Z Assembly

With the 3 end effectors mounted, attach the T brackets to the Z plate as shown using M3 screws:

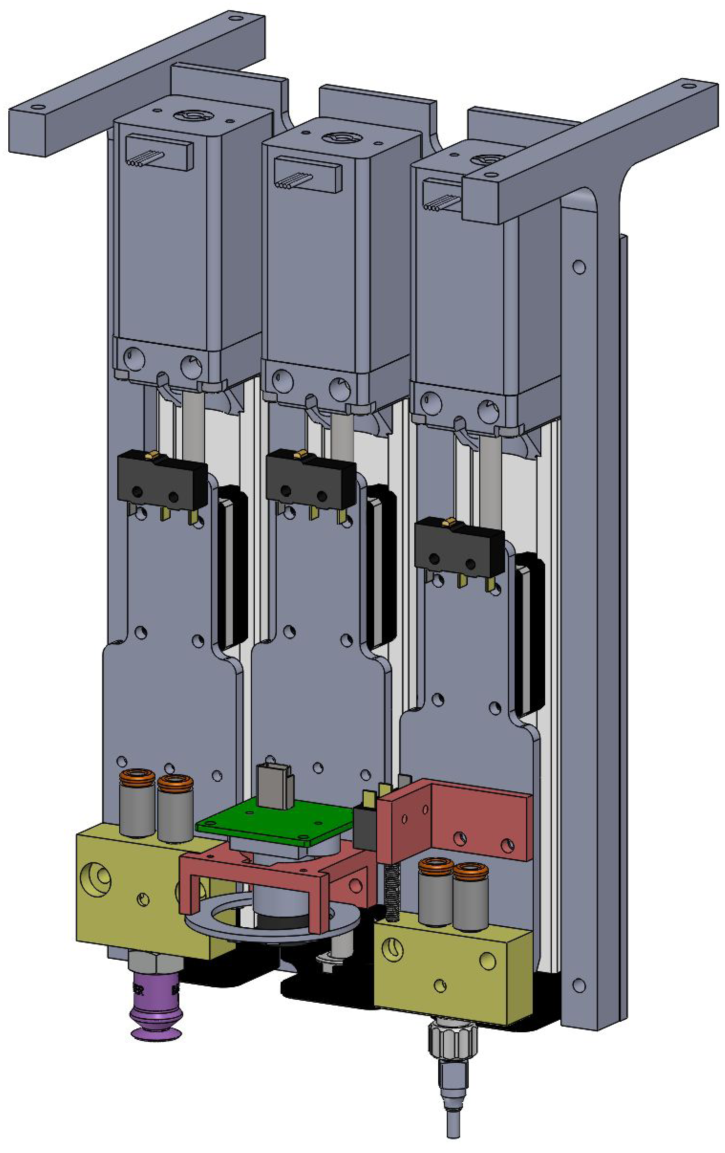

Then, attach the completed Z assembly to the underside of the lower Y plate. Ensure the 3 end effectors face forward (away from the larger drag chain)

## Wiring & Tubing

Unless otherwise noted, use 22 gauge stranded wire for all connections. It’s helpful, but strictly speaking not necessary, to have several colors on hand to help differentiate between signals (i.e. power vs. ground).

For crimping, we like the <u>PA-20 universal crimp tool</u>. But in a pinch, careful use of pliers will do the trick.

All limit switches are wired between the common terminal (labeled ‘C’) and the normally closed terminal (‘NC’). Thus, if they fail for some reason, the result is the same as the switch being depressed, and the software will not allow the robot to move.

### Threading the Drag Chains

It is easiest to thread wires and tubes through the drag chains several at a time (rather than all at once).

#### Large Drag Chain

The large drag chain carries:

- 3x ~9’ 10-pin ribbon cable (it’s helpful to label them at each end)
- 1x 10’ USB A-to-mini-B cable (the camera has a mini-B connector)
- 1x red (pink) 4 mm tube and 1x blue 4 mm tube (length depends on your air & vac connections)

Leave enough cable / tube on the entry end of the drag chain to connect to the Smoothieboard, air & vacuum outlets, and the PC (for the camera’s USB cable).

#### Small Drag Chain

The same cables & tubes exit the large drag chain, snake up through the slot in the Y axis, and then should be threaded through the small drag chain.

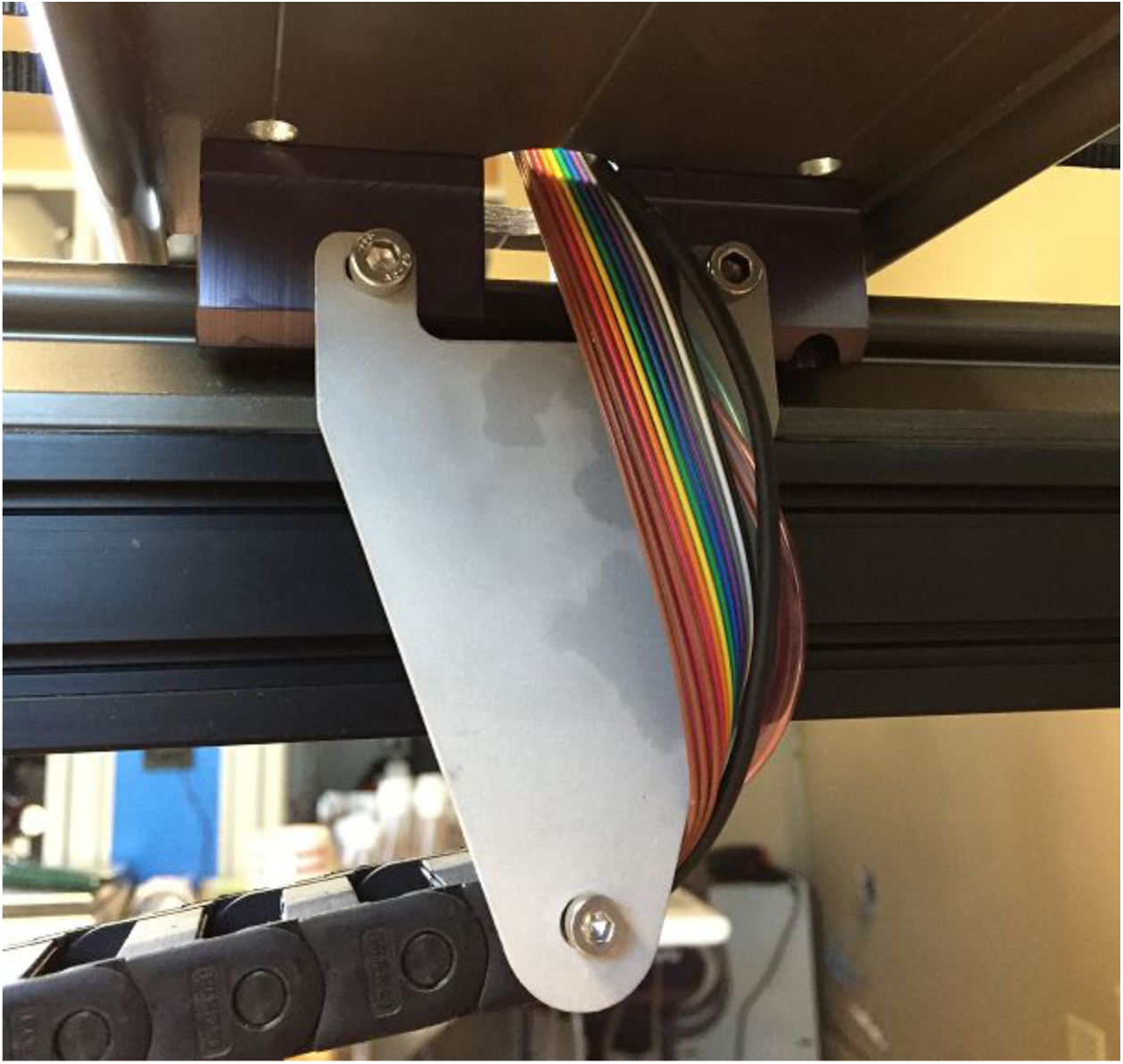

Connect the USB cable to the camera, and the pink and blue tubes to the inlets for the Clippard manifolds.

Use a vise or pliers to attach the ribbon cable connector plug to each end of the ribbon cables (it’s a 30 pin connector and there are 3 10-pin cables). The order and orientation of the cables doesn’t matter, as long as they are the same on each end.

The cables and tubes should pass to the left side of the Y carriage and around the back of the Z plate.

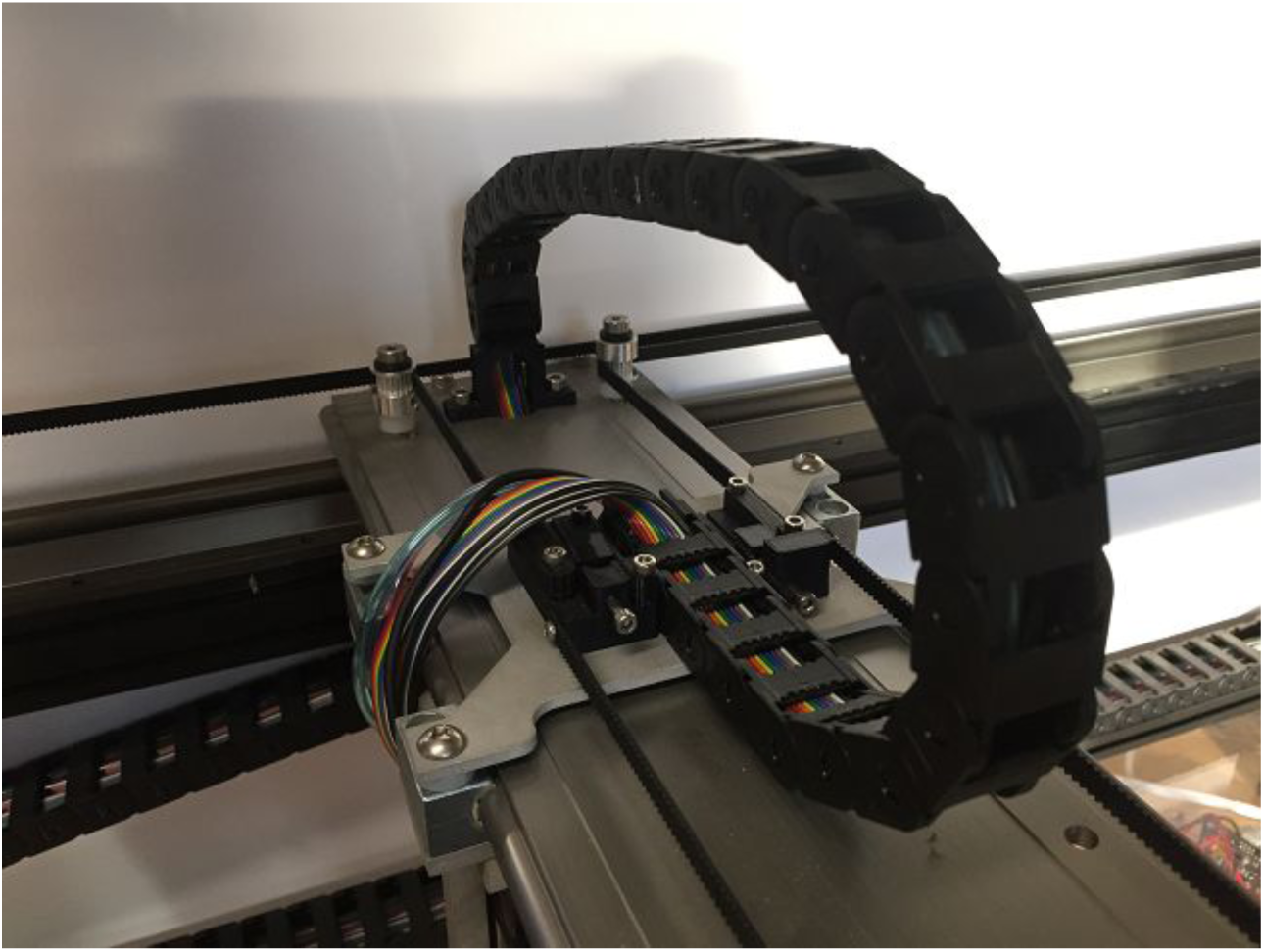

Wait to connect the ribbon cable to the PCB until after you’ve made all the wiring connections to that PCB.

### Power Supply

Connect the line, neutral & ground from the 120V cable to the appropriate terminals on the power supply. If desired, you can add a switch (DPDT) inline with the mains power.

Then connect wires (you should use something heavier than 22 gauge) to the +12 V and GND terminals on the power supply. The other end of those wires should go into the screw terminal block that mates with the power connector on the Smoothieboard for VBB. Be sure to check the polarity of that connector before powering the board.

### Smoothieboard

Solder approximately 1’ of wire to the ‘C’ and ‘NC’ terminals of a limit switch, and crimp Molex pins on to the other end (the pins come with the Smoothieboard). Insert into a 3 pin housing as shown below.

Similarly, crimp pins on to the ends of the 4 wires from each CoreXY stepper, and insert into 4 pin housings.

These components connect directly to the Smoothieboard. The rest connect to the Ribbon Breakout PCB on the back side of the Z plate.

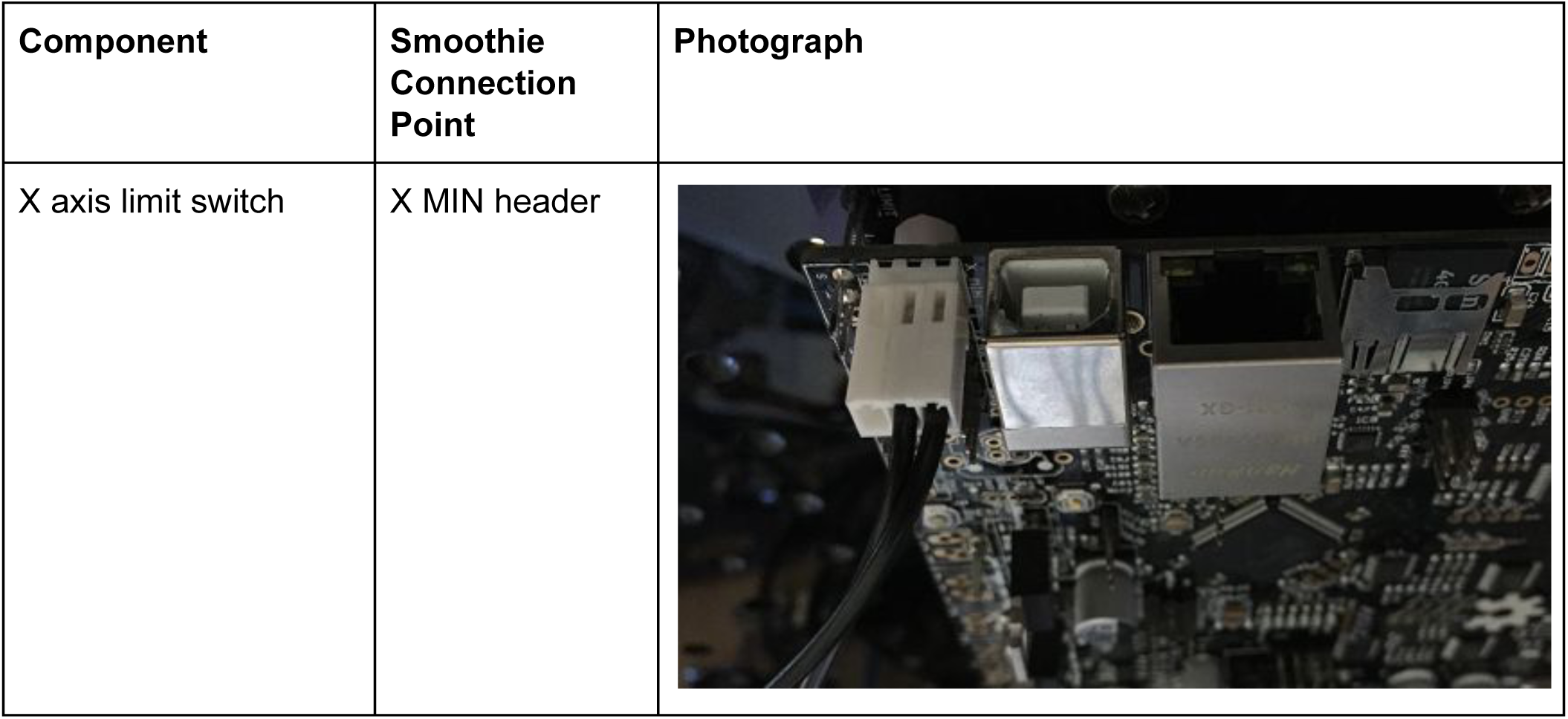

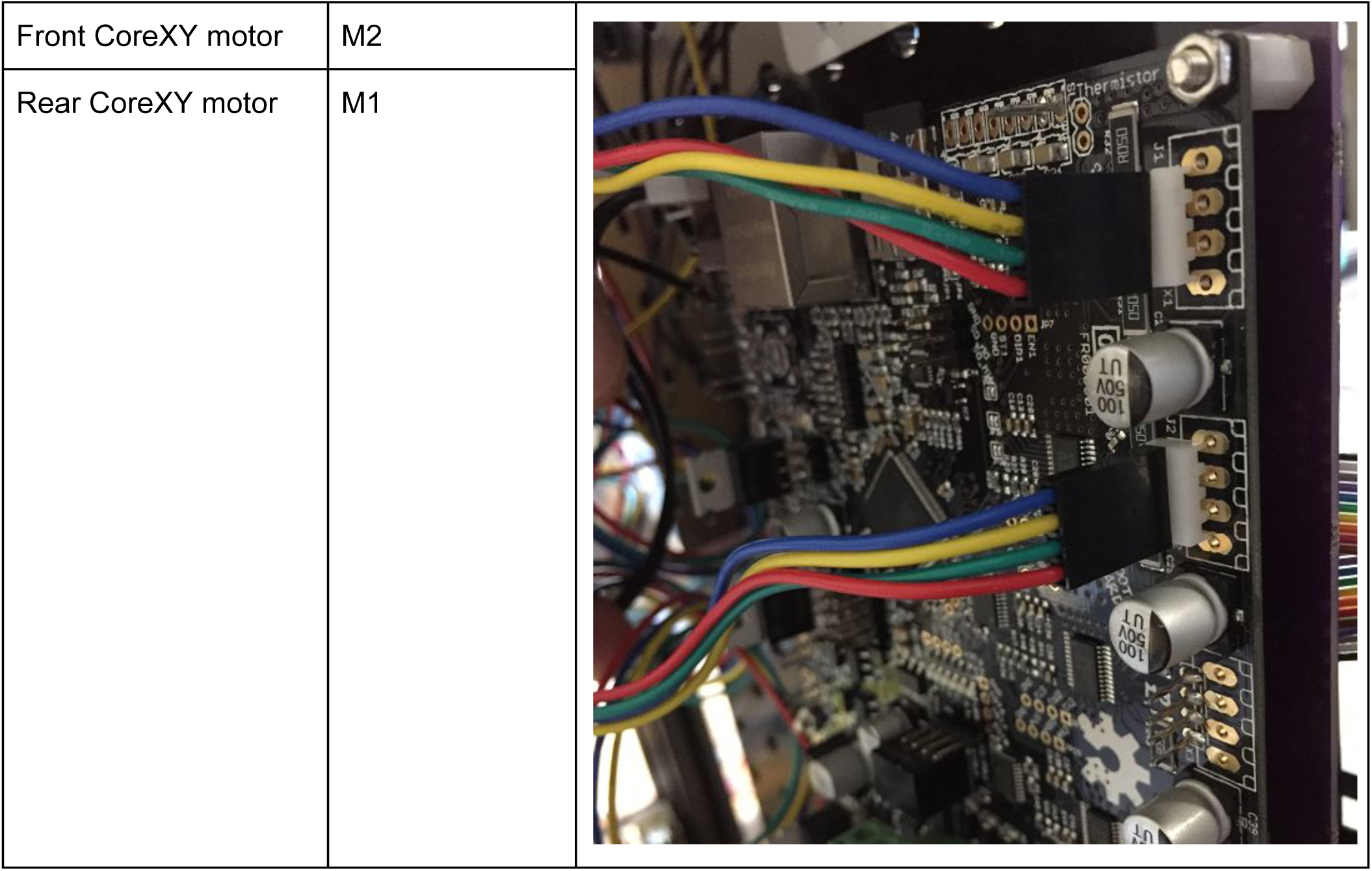

### Z Assembly

Before soldering to the Ribbon Cable Breakout PCB, prepare:

- 5x limit switches, soldering ~1’ lengths of wire to the C and NC terminals.
- The LED circuit board, with ~1’ lengths of wire soldered to the + and - terminals on the board (you may have to desolder the short lengths of wire that come on the board, or you can splice into them).
- The pressure sensor - solder ~1’ wires to pins 2 (red, 5V), 3 (black, GND), and 4 (blue, V_out_)

Use epoxy or hot glue to affix the LED circuit board to the camera holder, run the wires up and thru the U channel to the back side. Solder the ground wire to the square LED pin and the positive wire to the round LED pin.

Similarly, affix the pressure sensor in the small part manipulator manifold (the tube on the sensor fits inside the 3 mm hole in the manifold). Route the wires, then solder to the breakout PCB. The square pin is ground, next is +5 V, then V_out_.

Run the wires from the 3 Z slide stepper motors through the U channels to the back side of the Z plate, and trim them before soldering to the breakout PCB. Solder the wires from the common (‘C’) terminals to the square pads on the board and the ‘NC’ terminals to the round pads.

Attach 3 of the limits switches to 3 Z mounts. You will have to bend the two outside terminals on the switches so they don’t collide with the screws that hold the Z mounts to the slides. Run the wires up and through the U channels, then solder to the breakout board as labeled.

Attach the 4th limit switch to the underside of the lower Y plate, and solder the wires to the appropriate pins on the breakout board.

Finally, attach the 5th limit switch to the Z2 switch carrier, route the wires, and solder to the PCB.

Solder the wires from the solenoid valves to the PCB. The black wires should go to the square pads on the board, and the red wires to the round pads. Keep track of which solenoid is connected to which set of pad (labels might help), so you can connect the correct tubes in the next step.

As shown in the image below, a zip tie anchor can help with cable routing.

Similarly, zip tie anchors can be attached to the front side of the 3 stepper motors at the top of each Z slide, and cables/tubes fixed in place. Be sure to leave enough slack in the cables so that the Z axes can move to their lower limits.

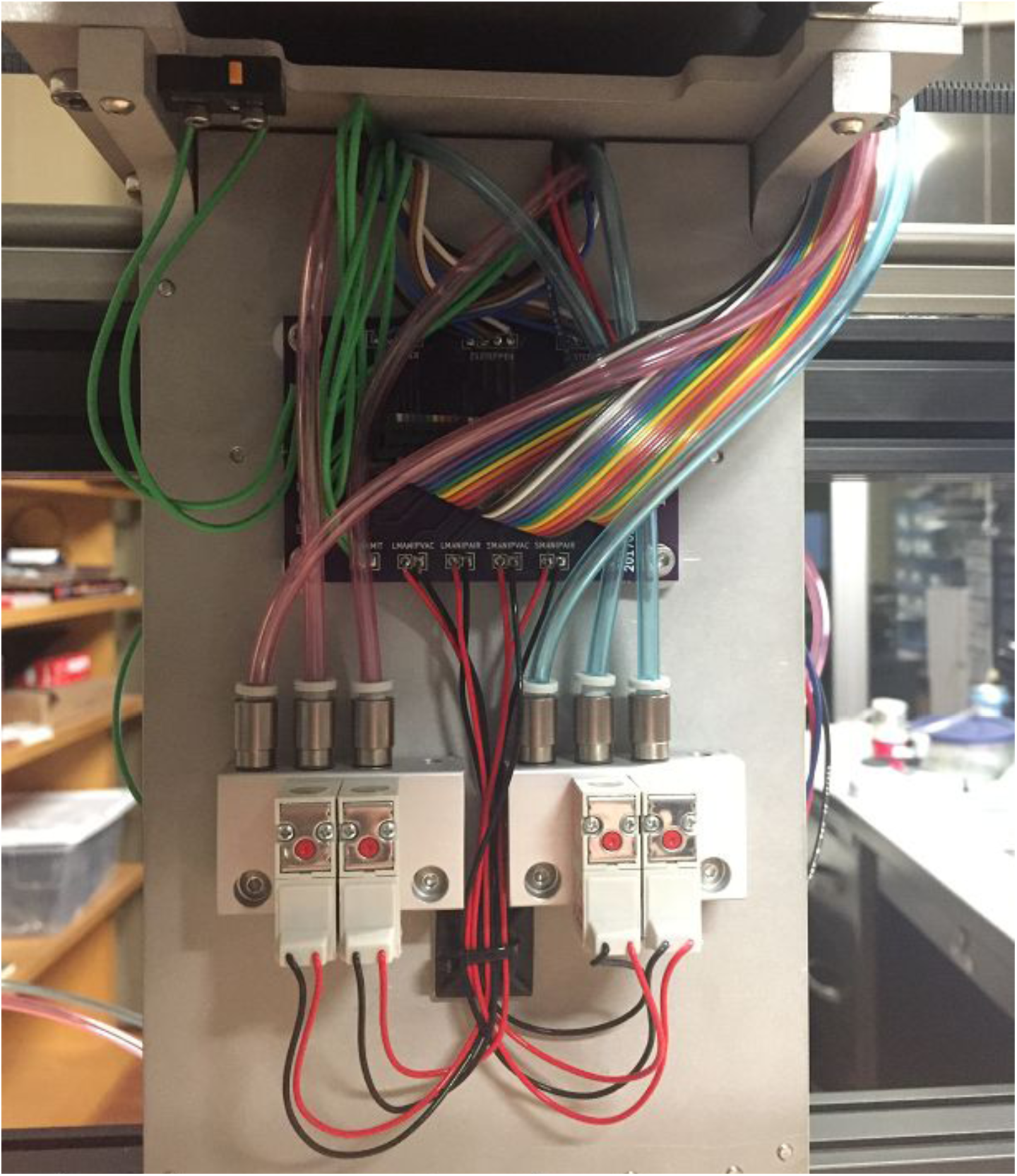

## Air & Vacuum Tubing

We use two different colors of tubing to differentiate positive and negative pressure. The red tubing specified in the BOM is actually a pink color, and we use the mnemonic "Pink is Positive,” leaving the blue tubing for vacuum connections.

Run a short length of pink and of blue tubing through each U-shaped cutout at the top of the Z plate (where it meets the lower Y plate). On the front side, attach the tubes to the small part and fly manipulators. On the back side, connect the two blue tubes to the vacuum manifold block and the pink tubes to the positive pressure manifold block.

## Firmware

Remove the microSD card from the Smoothieboard, mount it on your computer, and copy the *config* and *firmware.bin* files from the <u>Smoothie subdirectory of the MAPLE Control Software github repository</u> to the root directory of the card. Replace the SD card in the Smoothieboard, and when you next power the system, it will load the new firmware and use the updated config file.

## Alignment & Belt Tension

With the Smoothieboard not powered, unplug the CoreXY steppers from the Smoothieboard temporarily.

Use your hand to slide the Y carriage all the way to the right hand side of the robot. Tighten the 6x screws on each side of the Y carriage that fix it to the bearing blocks underneath.

Loosen the two M5 bolts that hold the front, top 40x40 aluminum extrusion to the right angle bracket on the left side of the machine. The slide the Y carriage all the way to the left side, and re-tighten the bolts.

Finally, tighten the M3 screws in the belt tensioner assemblies until the belts are quite snug. You aren’t trying to deform any parts here, but the belts should make a low tone when plucked.

## Testing

Download and install Pronterface - a program to send G-code commands to a COM port - http://www.Dronterface.com/

Slide the carriage to the middle of the workspace. Connect the Smoothieboard to your PC with a USB cable, and power the Smoothieboard. Use the X & Y jog buttons to move the robot 1 or 10 mm in each direction. +X should move the carriage to the left, +Y should move it toward the front. If these directions are reversed, you may need to swap the M1 and M2 connectors, or reverse one or both of them (be sure to power down the Smoothie when making any changes).

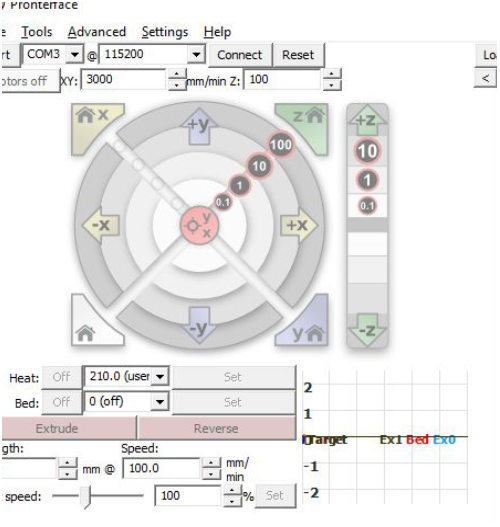

Similarly, jog the Z axis up and down to make sure it goes in the right direction. +Z is towards the bed.

Once you’re sure the robot is wired correctly, you can press the home button (or send the G28 command), which should zero the X, Y, Z (small part manipulator), A (camera) and B (fly manipulator) axes in sequence.

You’re now ready to download and install the MAPLE Control Software. Instructions and code available here: https://github.com/FlySorterLLC/MAPLEControlSoftware.

